# CMTM6 drives cisplatin resistance in OSCC by regulating AKT mediated Wnt signaling

**DOI:** 10.1101/2020.03.18.993774

**Authors:** Pallavi Mohapatra, Omprakash Shriwas, Sibasish Mohanty, Sandeep Rai Kaushik, Rakesh Arya, Rachna Rath, Saroj Kumar Das Majumdar, Dillip Kumar Muduly, Ranjan K Nanda, Rupesh Dash

## Abstract

Chemoresistance is one of the important factors for treatment failure in OSCC, which can culminate in progressive tumor growth and metastatic spread. Rewiring tumor cells to undergo drug-induced apoptosis is a promising way to overcome chemoresistance, which can be achieved by identifying the causative factors for acquired chemoresistance. In this study, to explore the key cisplatin resistance triggering factors, we performed global proteomic profiling of OSCC lines representing with sensitive, early and late cisplatin-resistant patterns. The top ranked up-regulated protein appeared to be CMTM6. We found CMTM6 to be elevated in both early and late cisplatin-resistant cells with respect to the sensitive counterpart. Analyses of OSCC patient samples indicate that CMTM6 expression is upregulated in chemotherapy-non-responder tumors as compared to chemotherapy-naïve tumors. Stable knockdown of CMTM6 restores cisplatin-mediated cell death in chemoresistant OSCC lines. Similarly, upon CMTM6 overexpression in CMTM6KD lines, the cisplatin resistant phenotype was efficiently rescued. Mechanistically, it was found that CMTM6 interacts with membrane bound Enolase-1 and stabilized its expression, which in turn activates the AKT-GSK3β mediated Wnt signaling. CMTM6 triggers the translocation of β-catenin into the nucleus, which elevates the Wnt target pro-survival genes like Cyclin D, c-Myc and CD44. Moreover, incubation with lithium chloride, a Wnt signaling activator, efficiently rescued the chemoresistant phenotype in CMTM6KD OSCC lines. In a patient-derived cell xenograft model of chemoresistant OSCC, knock-down of CMTM6 restores cisplatin induced cell death and results in significant reduction of tumor burden. CMTM6 has recently been identified as a stabilizer of PD-L1 and henceforth it facilitates immune evasion by tumor cells. Herewith for the first time, we uncovered another novel role of CMTM6 as one of the major driver of cisplatin resistance.

## Introduction

Head and neck cancer is the sixth most common cancer worldwide with approximately 53,260 new cases are being reported in United States alone ^1^. Almost 90% of HNSCC cancer cases are Oral squamous cell carcinoma (OSCC) which has emerged as the most common cancer in developing countries. In India, 80000 new OSCC cases are reported in each year with a mortality of ~46000 ^2^. OSCC patients commonly present with locally advanced (stage III or IV) disease. The treatment modalities of advanced OSCC are surgical removal of primary tumor followed by chemo-radiotherapy ^3^. However, neoadjuvant chemotherapy is commonly prescribed for surgically unresectable OSCC tumors that provide more surgical options ^4^. In spite of having all these treatment modalities, the 5-year survival rate of advanced tongue OSCC remains less than 50%, which indicates the development of resistance against existing therapy.

Chemoresistance is one of the major factors for treatment failure in OSCC. The common chemotherapy regimens for OSCC are Cisplatin alone or with 5FU and Docetaxel (TPF) ^5^. The tumor shows initial positive response to chemotherapy, but later it acquires chemoresistance, and patient experience relapse with onset of metastatic diseases. The chemoresistant properties could be attributed to enhanced cancer stem cell population, decreased drug accumulation, reduced drug-target interaction, reduced apoptotic response and enhanced autophagic activities ^6^. These hallmarks present the endpoint events, when cancer cell had already acquired chemoresistance. Few attempts have been made to understand the molecular mechanism of chemoresistance in HNSCC. Peng et al in 2012 demonstrate that tongue cancer chemotherapy resistance-associated protein 1 (TCRP1) is a modulator of cisplatin resistance in OSCC. TCRP1 expression is elevated specifically in cisplatin resistant cells, but not in 5FU resistant cancer cells. Analysis of clinical sample indicates that TCRP1 positive OSCC patients are resistant to cisplatin ^7^. Similarly, a shRNA based human kinome study elucidates that microtubule-associated serine/threonine kinase 1 (MAST1) is a major driver of cisplatin resistance in HNSCC. MAST1 inhibitor lestaurtinib, efficiently sensitized chemoresistant cells to cisplatin. Overall the study suggests that MAST1 is a viable target to overcome cisplatin resistance ^8^.

Here, to elucidate the causative factors those responsible for acquired chemoresistance, we have performed global proteomic profiling of OSCC lines representing with sensitive, early and late cisplatin-resistant patterns. The top ranked up-regulated protein was selected for validation in multiple cell lines and patient derived biopsy samples using qPCR, immunoblotting and immunohistochemistry. ShRNA based stable knock down of the identified important protein in cisplatin-resistant cells restored drug induced phenotype. The PDC based xenograft experiment suggests that knock down of the deregulated protein induces cisplatin-mediated cell death and facilitate significant reduction of tumor burden. Mechanistically, the deregulated molecule interacts with membrane bound Enolase-1 which in turn activates AKT/GSK3β mediated Wnt signaling. The activated Wnt pathway augments the chemoresistant phenotype. The identified deregulated molecule could be useful target to overcome cisplatin resistance in OSCC cells.

## Materials and methods

### Ethics statement

This study was approved by the Institute review Board and Human Ethics committees (HEC) of Institute of Life Sciences, Bhubaneswar (84/HEC/18) and All India Institute of Medical Sciences (AIIMS), Bhubaneswar (T/EMF/Surg.Onco/19/03). The animal related experiments were performed in accordance to the protocol approved by Institutional Animal Ethics Committee of Institute of Life Sciences, Bhubaneswar (ILS/IAEC-190-AH/DEC-19). Approved procedures were followed for patient recruitment and after receiving written informed consent from each patient, tissues samples were collected.

### Cell culture

H357, SCC-9 and SCC-4 (human tongue OSCC) cell lines were obtained from Sigma Aldrich, sourced from European collection of authenticated cell culture. All OSCC cell lines were cultured and maintained as described earlier. A549 (Lung carcinoma) and A375 (Melanoma) cell lines were obtained from NCCS, Pune and were maintained in DMEM supplemented with 10% FBS (Thermo Fisher Scientific) penicillin–streptomycin (Pan Biotech).

### Generation of early and late cisplatin resistance cell lines

For establishment of cisplatin resistant cell cell line, H357, SCC-9 and SCC-4 (OSCC), A549 (Lung carcinoma) and A375 (Melanoma) cell lines were initially treated with cisplatin at 1μM (lower dose) for a week and then the cisplatin concentration was increased gradually up to IC50 value, i.e. 15 μM for H357, SCC-9, SCC-4 and A549 and 10 μM for A375 in a span of 3 months. Parental cells were grouped as sensitive (CisS) and after a period of 4 and 8 months of treatment were termed as early (CisR 4M) and late resistant (CisR 8M) cells respectively.

### iTRAQ based proteomics analysis

Harvested cells (5X10^6^), from three time points (0M, 4M and 8M) were treated with RIPA buffer (Thermo Fisher Scientific, Cat #88665) supplemented with protease and phosphatase inhibitor (Sigma, Cat # P0044,). Extracted cellular proteins from all three time points with appropriate technical and biological replicates were used in an isobaric tag for relative and absolute quantification (iTRAQ) experiment (**Fig. 1B**). Equal amount of proteins (100 μg) from all samples were taken for tryptic protein preparation following manufacturer’s instructions (AB Sciex, USA). Study samples with the tag details used for labelling in iTRAQ experiment are presented in **Fig. 1B.** Trypsin treatment was performed using trypsin supplied by the manufacturer and incubating at 37°C for 16-20 hrs. Tryptic peptides were dried at 40°C using SpeedVac (LabConco, USA). Dried tryptic peptides were dissolved using dissolution buffer and isobaric tags reconstituted with isopropanol were added to be incubated at room temperature for 2 h. After the completion of the reaction, tagged tryptic peptides from all samples were pooled and dried. Tagged tryptic peptides (~250 μg) were subjected to strong cation exchange fractionation using a hand-held ICAT® Cartridge-cation-exchange system (Applied Biosystems, USA). Peptides were eluted using a gradient 30, 50, 80, 120, 250, 300, 400 and 500 mM of ammonium formate solutions with a flow rate of 10 drops/min. Each SCX fraction was dried at 40°C using a Speed Vac (CentriVap, Labconco, USA) and cleaned using Pierce C18 spin column (ThermoFisher Scientific Inc., USA).

**Figure 1:**
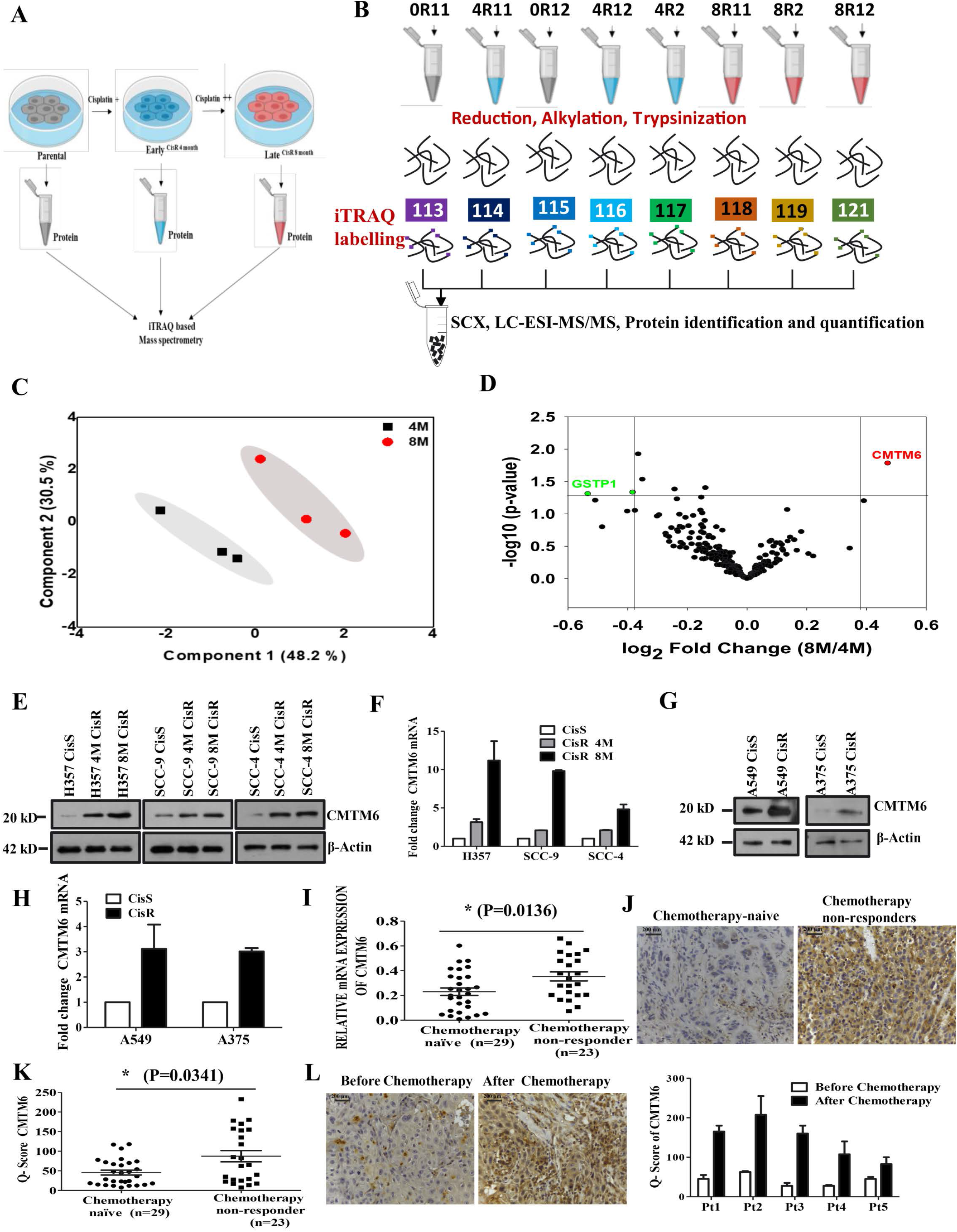
CMTM6 is up regulated in chemoresistant squamous cell carcinomas: **A)** Schematic representation of sensitive, early and late cisplatin resistant OSCC line for global proteomic profiling. The establishment of sensitive, early and late resistant cells are described in materials and method section. **B)** The lysates were isolated from parental sensitive (H3457CisS), early (H357CisR4M) and late (H357CisR8M) cisplatin resistant cells and subjected to global proteomic profiling. The schematic diagram depicts the iTRAQ labelling strategy for proteomic analysis. 0R11 and 0R12 are biological replicates of H357CisS group, 4R11: 4R12 and 4R2 are technical and biological replicates of H357CisR4M group, 8R11: 8R12 and 8R2 are technical and biological replicates of H357CisR8M group. **C)** Principal component analysis of global proteomic profiling sensitive, early (4M) and late resistant cells (8M). **D)** Volcano plot indicating deregulated genes in proteome profiling of sensitive and cisplatin resistant cells. CMTM6 is the top ranked up regulated genes in 4M and 8M cisplatin resistant groups. **E)** Cell lysates from indicated resistant and sensitive OSCC cells were isolated and subjected to immunoblotting against CMTM6 and β-actin antibodies. **F)** Relative mRNA (fold change) expression of CMTM6 was analyzed by qRT PCR in indicated cells (mean ±SEM, n=3). **G)** Cell lysates from indicated resistant and sensitive OSCC cells were isolated and subjected to immunoblotting against CMTM6 and β-actin antibodies. **H)** Relative mRNA (fold change) expression of CMTM6 was analyzed by qRT PCR in indicted cells (mean ±SEM, n=3). **I)** Relative mRNA expression of CMTM6 was analyzed by qRT PCR in different chemotherapy-non-responder OSCC tumors as compared to chemotherapy-naïve tumors (Median, n=29 for chemotherapy-naïve and n=23 for chemotherapy-non-responder). *: P < 0.05. **J)** Protein expression of CMTM6 was analyzed by IHC in chemotherapy-naïve and chemotherapy-non-responder OSCC tumors. **K)** IHC scoring for CMTM6 from panel J (Q Score =Staining Intensity × % of Staining), (Median, n=29 for chemotherapy-naïve and n=23 for chemotherapy-non-responder) *: P < 0.05. **L)** Left panel: Protein expression of CMTM6 was analyzed by immunohistochemistry (IHC) in pre- and post-TPF treated paired tumor samples for chemotherapy-non-responder patients Right panel: Q Score =Staining Intensity × % of IHC Staining.

Each SCX fraction was resuspended in 20 μl of buffer (water with 0.1% formic acid) and introduced to easy-nanoLC 1000 HPLC system (Thermo Fisher Scientific, Waltham, MA) connected to hybrid Orbitrap Velos Pro Mass Spectrometer (Thermo Fisher Scientific, Waltham, MA). The nano-LC system contains the Acclaim PepMap100 C18 column (75 μm × 2 cm) packed with 3 μm C18 resin connected to Acclaim PepMap100 C18 column (50 μm × 15 cm) packed with 2 μm C18 beads. A 120 min gradient of 5% to 90% buffer B (0.1% formic acid in 95% Acetonitrile) and Buffer A (0.1% formic acid in 5% Acetonitrile) was applied for separation of the peptide with a flow-rate of 300 nl/min. The eluted peptides were electrosprayed with a spray voltage of 1.5 kV in positive ion mode. Mass spectrometry data acquisition was carried out using a data-dependent mode to switch between MS1 and MS2.

### Protein Identification and iTRAQ Quantitation

Protein identification and quantification was carried out using SEQUEST search algorithm of Proteome Discoverer Software 1.4 (Thermo Fisher, Waltham, MA, USA). Each MS/MS spectrum was searched against a human proteome database (UniProt, 89,796 total proteins, downloaded in April 2017). Precursor ion mass tolerance (20 ppm), fragmented ion mass tolerance (0.1 Da), missed cleavages (<2) for trypsin specificity, Carbamidomethyl (C), Deamidation (N and Q), Oxidation (M) and 8-plex iTRAQ label (N terminus and K) were set as variable modifications. The false discovery rate (FDR) at both protein and peptide level was calculated at 5%. The identified protein list with fold change values were exported to Microsoft Excel for further statistical analysis. Identified proteins from study samples and relative fold change values were selected for principal component analysis and a partial least square discriminate analysis model was built using MetaboAnalyst 3.0. Proteins with at least 2.0 fold change (log_2_ resistance/sensitive>±1.0, p<0.05, variable important projection value>1.0) were selected as deregulated proteins. All the mass spectrometry data files (.raw and .mgf) with result files were deposited in the ProteomeXchange Consortium (PXD016977). The deregulated proteins, identified from global proteomics analysis, were converted to gene list and a functional analysis was carried out using Ingenuity Pathway Analysis (IPA).

### Lentivirus production and generation of stable CMTM6 KD cell lines

ShRNAs targeting CMTM6 were cloned into pLKO.1 vector as per the protocol mentioned by addgene. Cloning of shRNAs were followed by confirmation using sequencing method. Lentivirus was produced by transfection of pLKO.1-shRNA plasmid along with packaging plasmid psPAX2and envelop plasmid pMD2G into HEK293T cells. Lentivirus particles were generated using protocol as described in Shriwas et al ^9^. Lentivirus infected cells were incubated with puromycin up to 5 μg/ml for two weeks, stable clones were picked and confirmed by immunoblotting. All shRNA sequences used in this study are mentioned in supplementary table 1.

### Transient transfection and rescue of CMTM6 expression in CMTM6 KD cell lines

CMTM6 knockdown cells, stably expressing shRNA#2 targeting 3’ UTR of CMTM6 mRNA, were transiently transfected with pCMV6 CMTM6 (Myc-DDK-tagged) (Origene, Cat #RC201061) using the ViaFect transfection reagent (Promega Cat# E4982). The cells were transfected for 48h, after which they were treated with different concentration of cisplatin followed by flow cytometry analysis (Annexin V PE/7-AAD Assay), Cell viability assay(MTT) and immunoblotting with anti PARP, Anti p^s-139^-H2AX and Anti β-actin. The transfection efficiency was confirmed by immunoblotting against Anti-CMTM6 and Anti-DDK.

### OSCC patient sample

Biopsy samples were collected from clinical sites and divided into two study groups namely chemotherapy-naive patients (n=29, OSCC patients those were never treated with any chemotherapy) and OSCC chemotherapy non-responders (n=23, OSCC patients those were treated with neoadjuvant chemotherapy but never responded or partially responded). The tumor samples were collected from AIIMS, Bhubaneswar and HCG Panda Cancer Centre, Cuttack. The list of patients with treatment modalities and other clinical information are described in table 1.

**Table 1a:** chemotherapy-naïve Patient details.

| Sl No | Tumor samples | Age/Sex | Site of disease | Clinical stage |
| --- | --- | --- | --- | --- |
| 1 | Patient#1 | 32/M | Tongue Lt lateral border | T4N1M0 |
| 2 | Patient#2 | 29/F | Rt- Upper gingivobuccal maxillary antrum | T2N1Mx |
| 3 | Patient#3 | 60/F | Upper alveolus | T4aN1M0 |
| 4 | Patient#4 | 42/M | Tongue Lt lateral border | T4N1M0 |
| 5 | Patient#5 | 60/M | Lt- Buccal mucosa | T4aN1M |
| 6 | Patient#6 | 67/M | Tongue Lt lateral border | T4aN1Mx |
| 7 | Patient#7 | 33/M | Tongue Lt lateral border | T3N1Mx |
| 8 | Patient#8 | 55/F | Rt- Buccal mucosa | T4bN2bM0 |
| 9 | Patient#9 | 50/M | Rt- Buccal mucosa | T4aN2bM0 |
| 10 | Patient#10 | 42/M | Tongue Rt lateral border | T4aN2bM0 |
| 11 | Patient#11 | 52/M | Tongue Lt lateral border | T2N1Mx |
| 12 | Patient#12 | 38/M | Lt-Buccal Mucosa | T4N1M0 |
| 13 | Patient#13 | 75/M | Rt- Buccal mucosa | T4aN2bM0 |
| 14 | Patient#14 | 74/M | Tongue lateral border | T2N0M0 |
| 15 | Patient#15 | 53/M | Tongue Lt lateral border | T4N0M0 |
| 16 | Patient#16 | 34/M | Lt Lt-Buccal Mucosa | T4aN2bMx |
| 17 | Patient#17 | 40/M | Tounge- lateral border | T2N2cM0 |
| 18 | Patient#18 | 74/M | Tounge | T2N0M0 |
| 19 | Patient#19 | 53/M | Oral cavity | T4N0M0 |
| 20 | Patient#20 | 45/M | Tounge | T4aN2cM0 |
| 21 | Patient#21 | 46/M | Tounge | T3N1M0 |
| 22 | Patient#22 | 35/M | Tounge | T4aN2eM0 |
| 23 | Patient#23 | 36/M | Tounge | cT4aN2aM0<br>cT4aN2cM0 |
| 24 | Patient#24 | 48/M | Left Buccal Mucosa |  |
| 25 | Patient#25 | 38/M | SCC Base of Tongue with extension into Oropharynx | T4aN2cMO |
| 26 | Patient#26 | 36/M | Right Buccal Mucosa | T4aN2aM0 |
| 27 | Patient#27 | 34/M | Left Buccal Mucosa | T4aN2bMx |
| 28 | Patient#28 | 38/M | Right Gingiva buccal sulcus | T4N1Mx |
| 29 | Patient#29 | 40/M | Tounge | T2N2M0 |

Lt-Left, Rt-Right

**Table 1b:** Chemotherapy-non-responders patient Details.

| Sl No | Tumor samples | Age /Sex | Site of disease | Clinical stage | Chemotherapy (NACT) | Cycle | Response |
| --- | --- | --- | --- | --- | --- | --- | --- |
| 1 | Patient# 1 (PDC#1) | 76/M | Tongue Rt lateral border | T4N0M0 | Paclitaxel + Cisplatin | 3 | non responder |
| 2 | Patient#2 | 51/M | Rt- Buccal mucosa | T2N2bM0 | Docetaxel + Cisplatin+ 5FU | 2 | non responder |
| 3 | Patient# 3 | 60/M | Tongue Rt lateral border | T3N1M0 | Paclitaxel + Cisplatin +5FU | 3 | non responder |
| 4 | Patient#4 | 33/M | Rt- Lower Alveolar mucosa | T3N1Mx | Docetaxel + Cisplatin | 3 | non responder |
| 5 | Patient#5 | 60/F | Tongue Lt lateral border | T4N0M0 | Docetaxel + Cisplatin+ 5FU | 3 | non responder |
| 6 | Patient#6 | 59/M | Tongue Rt lateral border | T4aN1M0 | Docetaxel + Cisplatin+ 5FU | 3 | non responder |
| 7 | Patient#7 | 46/M | Tongue | T4N3M0 | Docetaxel + Cisplatin+ 5FU | 3 | non responder |
| 8 | Patient#8 | 55/F | Rt- Buccal Mucosa | T4aN2M0 | Docetaxel + Cisplatin+ 5FU | 2 | *Partial responder |
| 9 | Patient#9 | 37/M | Tongue | T4N3M0 | Docetaxel + Cisplatin+ 5FU | 2 | non responder |
| 10 | Patient#10 | 27/M | Lt-Buccal Mucosa | T4N2M0 | Docetaxel + Cisplatin+ 5FU | 2 | *Partial responder |
| 11 | Patient#11 | 46/F | Rt- oral cavity | T4N1M0 | Docetaxel + Cisplatin+ 5FU | 3 | non responder |
| 12 | Patient#12 | 42/M | Rt- Buccal Mucosa | TxN3bM0 | Paclitaxel + Cisplatin+ 5FU | 2 | non responder |
| 13 | Patient#13 | 30/M | Tongue Rt lateral border | T2N0Mx | Paclitaxel + Cisplatin+ 5FU | 3 | non responder |
| 14 | Patient#14 | 52/M | Rt- Buccal Mucosa | T4N2M0 | Docetaxel + Cisplatin+ 5FU | 3 | non responder |
| 15 | Patient#15 | 32/M | Tongue | T3N1M0 | Docetaxel + Cisplatin+ 5FU | 3 | non responder |
| 16 | Patient#16 | 35/M | Tongue | T4aN2aM0 | Docetaxel + Cisplatin+ 5FU | 3 | *Partial responder |
| 17 | Patient#17 | 36/M | Left Buccal Mucosa | T4aN2aM0 | Docetaxel + Cisplatin+ 5FU | 3 | non responder |
| 18 | Patient #18 | 38/M |  | T4bN2bM0 | Docetaxel + Cisplatin+ 5FU | 2 | *Partial responder |
| 19 | Patient # 19 | 55/M | Tounge, Left lateral border, |  | Docetaxel + Cisplatin+ 5FU | 3 | *Partial responder |
| 20 | Patient # 20 | 35/M | Tounge | T4aN2eM0+ | Docetaxel + Cisplatin+ 5FU | 3 | non-Responder |
| 21 | Patient # 21 | 36/M | Left Buccal Mucosa | cT4aN2aM0 | Docetaxel + Cisplatin+ 5FU | 3 | Non Responde |
| 22 | Patient # 22 | 39/M | Right mandible | cT4bN0Mx | Docetaxel + Cisplatin+ 5FU |  | Non Responder |
| 23 | Patient # 23 | 55/M | Tounge, Left lateral border, | cT4aN2cM0 | Docetaxel + Cisplatin+ 5FU | 3 | Non responder |

\*Partial response- Patient initially responded to chemotherapy but after 1-2 cycles became non responded.
Chemotherapy Doses: Cisplatin: 100mg, Paclitaxel: 260 mg, Docetaxel: 100mg, 5FU:1000mg

### Immunoblotting

Cell lysates were used for immunoblotting experiments as described earlier ^10^. For this study, we used antibody against β-actin (Sigma, Cat#A2066), PARP (CST, Cat #9542L), p^s-139^-H2AX (CST, Cat # 9718S), CMTM6 (Sigma, Cat#HPA026980), DDK (CST: Cat#14793), AKT(CST, Cat #9272S), pAKT(Ser473) (CST, Cat #4058S), GSK-3β(CST, Cat#9315s), Phospho-GSK-3β (Ser9) (CST, Cat #9323S), β-Catenin (CST, Cat#9562), Phospho-β-Catenin (Ser552)(CST, Cat #9566), Non-phospho (Active) β-Catenin (Ser33/37/Thr41) (CST, Cat #8814), CD44 (Novus, Cat#NBP1-31488), TCF4/TCF7L2 (C48H11) (CST, Cat#2569S), c-Myc(CST, Cat # 9402), Cyclin D1(CST, Cat #2922S), LRP1(Cloud clone, Cat # PAB010Hu01), PSMD2 (Cloud clone, Cat# PAG279Hu01), α Enolase(L-27)(Sanatcrutz, Cat #sc-100812), Sox2 (CST, Cat #2748), Nanog (CST, Cat #4893S), and Oct-4 (CST, Cat #2750S). Membrane and cytoplasmic fractions were separated using the Mem-PER™ Plus Membrane Protein Extraction Kit (ThermoFisher Scientific, Cat #78833), according to manufacturer’s instructions.

### Co-Immunoprecipitation

For co-immunoprecipitation experiments, cells were lysed in 1% digitonin for 30 min on ice. The lysates were incubated with primary antibody for 1–4 h, followed by addition of Protein A/G PLUS-Agarose beads (Santa cruz, Cat #sc-2003) for overnight at 4 °C. After four washes in 0.2% digitonin, samples were eluted in SDS sample buffer with 50 mM DTT for 10 min at 70 °C, separated by SDS–PAGE and immunoblotted. VeriBlot for IP Detection Reagent (HRP) (Abcam, Cat #ab131366) was used for immunoblotting.

### Patient Derived Xenograft

BALB/C-nude mice (6-8 weeks, male, NCr-Foxn1nu athymic) were purchased from Vivo biotech Ltd (Secunderabad, India). For xenograft model early passage of patient-derived cells (PDC1) established from chemo non-responder patient (treated with TP,50 mg carboplatin and 20 mg paclitaxel for 3 cycles without having any response) was considered. Two million cells were suspended in phosphate-buffered solution-Matrigel (1:1, 100 μl) and transplanted into upper flank of mice. The PDC1 cells stably expressing NtShRNA were injected in right upper flank and PDC1 cells CMTM6ShRNA#1 (PDC1 CMTM6KD) were injected in the left upper flank of same mice. These mice were randomly divided into 2 groups (n=6) once the tumor reached volume of 50 mm^3^ and injected with vehicle or cisplatin (3 mg/Kg) intraperitonially twice a week. Tumor size was measured using digital vernier calliper twice a week until the completion of experiment. Tumor volume was determined using the following formula: Tumor volume (mm^3^) = (minimum diameter)^2^ × (maximum diameter)/2.

### RT-PCR and Real Time Quantitative PCR

RNA mini kit (Himedia, Cat# MB602) was used to isolate total RNA as per manufacturer’s instruction and quantified by Nanodrop. c-DNA was synthesised by reverse transcription PCR using Verso cDNA synthesis kit (ThermoFisher Scientific, Cat # AB1453A) from 300 ng of RNA. qRT-PCR was carried out using SYBR Green master mix (Thermo Fisher scientific Cat # 4367659). GAPDH was used as a loading control. The primer (oloigos) details used for qRT-PCR in this article are listed in supplementary table 1.

### Immunohistochemistry

Immunohistochemistry of formalin fixed paraffin-embedded samples (OSCC patients tumor and Xenograft tumors from mice) were performed as previously described ^11^. Antibodies against CMTM6 (Sigma: Cat#HPA026980), β-Catenin (CST, Cat#9562), Non-phospho (Active) β-Catenin (Ser33/37/Thr41) (CST, Cat #8814), Cyclin D1(CST, Cat #2922S) and Ki67 (Vector, Cat #VPRM04) were used for IHC. Images were obtained using Leica DM500 microscope. Q-score was calculated by multiplying percentage of positive cells with staining (P) and intensity of staining (I). P was determined by the percentage of positively stained cells in the section and I was determined by the intensity of the staining in the section i.e. strong (value=3), intermediate (value=2), weak (value=1) and negative (value=0).

### Annexin V PE/7-ADD Assay

Apoptosis and cell death assay was performed by using Annexin V Apoptosis Detection Kit PE (eBioscience™, USA, Cat # 88-8102-74) as described earlier ^9^and cell death was monitored using a flow cytometer (BD FACS Fortessa, USA).

### Assessment of cell viability

Cell viability was measured by 3-(4, 5-dimethylthiazol-2-yl)-2, 5-diphenyltetrazolium bromide (MTT; Sigma-Aldrich) assay.

### Immunofluorescence

The cells were seeded on lysine coated coverslip and cultured for overnight. Cells were fixed with 4% formaldehyde for 15 min, permeabilized with 1 × permeabilization bufer (eBioscience 00-8333-56) followed by blocking with 3% BSA for 1 h at room temperature. Then the cells were incubated with primary antibody overnight at 4 °C, washed three times with PBST followed by 1hr incubation with Goat anti–Rabbit IgG(H+L) secondary Antibody, Alexa Fluor® 488 conjugate (Invitrogen, Cat #A -11008) and Rabbit anti– Mouse IgG(H+L) secondary Antibody, Alexa Fluor® 647 conjugate (Invitrogen, Cat #A – 21239). After final wash with PBST (thrice) coverslips were mounted with DAPI (Slow Fade ® GOLD Antifade, Thermo Fisher Scientific, Cat # S36938). Images were captured using a confocal microscopy (LEICA TCS-SP8). Anti CMTM6 (Sigma: Cat#HPA026980), Anti α Enolase (L-27)(Sanatcrutz, Cat #sc-100812) and anti Non-phospho (Active) β-Catenin (Ser33/37/Thr41) (CST, Cat #8814) were used in this study.

### Colony formation assay

Colony formation assay was performed as described in Shriwas et al ^9^

### Dual luciferase reporter assay

For this assay cells were co-transfected with M50 Super 8x TOPFlash (which was a kind gift from Randall Moon, Addgene, Cat #12456) ^12^ and pRL-TK (Promega) in a ratio of 100:1 using ViaFect transfection reagent (Promega Cat# E4982). Seven TCF/LEF-binding sites are present upstream of a firefly luciferase gene in the TOPflash vector, whereas pRL-TK provides constitutive expression of Renilla luciferase and was used as an internal control for the experiment. Cells were treated with vehicle control, LiCl (Sigma Aldrich, Cat #62476-100G-F) and Pyrvinium (Sigma Aldrich, Cat #P0027-10MG) 48h after transfection and luciferase reading was taken using the Dual-Glo luciferase assay kit (Promega, Cat # E1910) as per the manufacturer’s instructions.

### Tumorsphere formation assay

1000 cells were seeded on six-well ultralow attachment plates (Corning-Costar Inc.) and were grown with 1×B27 (Invitrogen 17502048), 1×N2 supplement (Invitrogen 17502048), 20 ng/mL of human recombinant epidermal growth factor (Invitrogen PHG0313), 10 ng/mL of basic fibroblast growth factor (Invitrogen PHG0263) in serum-free DMEM-F12 medium (Pan biotech P04-41500). After spheroid formation, treatment was done with DMSO and cisplatin. Images were captured in a microscope (LEICA DMIL).

### ALDH activity assays

The aldehyde dehydrogenase (ALDH) activity was detected using an ALDEFLUOR assay kit (Stem Cell Technologies, Canada, Cat #1700) according to the manufacturer’s protocol. Cells were incubated with ALDH protein substrate (BAAA) in Aldefluor assay buffer for 30 min at 37°C. A specific inhibitor of ALDH (DEAB) was used as a negative control. Fluorescence was measured by BD LSR Fortessa.

### Correlation analysis of β-Catenin target genes and CMTM6

The correlation analysis was done between β-Catenin target genes and CMTM6 in HNSCC patient tumors using GEPIA (http://gepia.cancer-pku.cn/detail.php?gene=CMTM6), online analysis software based on the TCGA database and Genotype, using |log2FC|≥1 and P≤0.05 as the cut-off criteria.

### Statistical analysis

All data points are presented as mean and standard deviation and Graph Pad Prism 5.0 was used for calculation. The statistical significance was calculated by one-way variance (one-way ANOVA), Two-Way ANOVA and considered significance at P≤0.05.

## Results

### Generation and characterization of early and late cisplatin resistant cells

First, we evaluated the cisplatin-induced cell death in sensitive early and late cisplatin resistant pattern (CisS, CisR4M and CisR8M) of H357, SCC-9 and SCC-4 cells by performing MTT (cell viability) assay. The data suggested CisR8M achieved complete acquired resistance and CisR4M achieved partial resistance (Supplementary Fig. 1A-B). In addition to this, we established CisS and CisR 8M pattern of A549 (human lung cancer line) and A375 (human melanoma line). The cell viability (MTT) assay suggests that A549CisR and A375 CisR acquired resistance to cisplatin as compared to their sensitive counterparts i.e. A549CisS and A375 CisS (Supplementary Fig. 1C).

### CMTM6 expression is elevated in chemoresistant cancer cells

To explore the causative factors responsible for acquired cisplatin resistance, we performed global proteomic profiling of H357CisS, H357CisR 4M and H357CisR8M cells. From the proteomic profiling, a set of 247 proteins were identified and 44 showed deregulation (log_2_(resistance/sensitive)>±1.0 and VIP score>1.6) (Supplementary table 2). Principal component analysis (PCA), taking all the identified proteins as variables with their fold change values grouped these samples into three separate clusters (Fig. 1A-C). The dendrogram indicates that CMTM6 is up regulated in early and late resistant cells as compared to sensitive cells (Supplementary Fig. 2). The volcano plot data suggests that CMTM6 is the top ranked upregulated protein in early and late cisplatin resistant cells as compared to sensitive counterpart (Fig. 1D). Based on these data, we selected CKLF-like MARVEL transmembrane domain containing protein 6 (CMTM6) for further validation and to explore its potential role in driving cisplatin-resistance. Henceforth, we evaluated the CMTM6 expression in H357, SCC-9 and SCC-4 cell lines showing early and late chemoresistant patterns (H357 CisR4M, H357 CisR8M, SCC-4 CisR4M, SCC-4 CisR8M, SCC-9 CisR4M and SCC-9 CisR8M). Expressions of CMTM6 at protein and mRNA levels were found to be up-regulated in CisR4M and CisR8M cells with respect to CisS counterparts in all cell lines (Fig. 1E, F). We also observed elevated CMTM6 expression in lung cancer line A549CisR and melanoma line A375CisR cells as compared to their sensitive counterpart (Fig. 1G, H). Here it is important to mention that we indicate CisR for CisR8M or late cisplatin resistant pattern. To evaluate the clinical significance of our observation, we monitored CMTM6 expression in tumors isolated from drug naïve freshly diagnosed OSCC patients and non-responders, i.e. patients not responding or partially responding to neoadjuvant chemotherapy (TPF). Based on qRT-PCR and immunohistochemistry data, higher abundance of CMTM6 was observed in tumor tissues isolated from non-responders to drug naive OSCC patients (Fig. 1I-K). Similarly, we determined the expression of CMTM6 in drug-naive and post chemotherapy treated paired tumor samples not responding to treatment. The post chemotherapy treated tumor samples showed higher expression of CMTM6 (Fig. 1L). Overall, it was found that CMTM6 expression is significantly elevated in chemoresistant lines.

### Targeting CMTM6 reverses cisplatin resistance in squamous cell carcinomas

To delineate the potential role of CMTM6 as a major driver of cisplatin resistance, we generated stable CMTM6 knock down clones in chemoresistant lines using a lentivirus based ShRNA approach. To knock down CMTM6 we used two different ShRNAs, one targeting the coding sequence (CMTM6ShRNA#1) and the other targeting 5’UTR (CMTM6ShRNA#2) of CMTM6 mRNA. Stable clones generated by both the ShRNA showed efficient knock down of CMTM6 (Fig. 2A). Our cell viability and cell death assay data suggests that knock down of CMTM6 significantly sensitized chemoresistant lines to cisplatin (Fig. 2B-D). Similarly, after knocking down CMTM6 in patient-derived tumor primary cells not responding to TP, PDC1 cells reversed resistance and became sensitive to cisplatin (Figure 2B-D). Our immunoblotting data suggests that knocking down CMTM6 in chemoresistant cells results in significant increase in cisplatin induced cleaved caspase and γ-H2AX expression (Fig. 2E). These data suggest CMTM6 dependency of chemoresistant OSCC cells.

**Figure 2:**
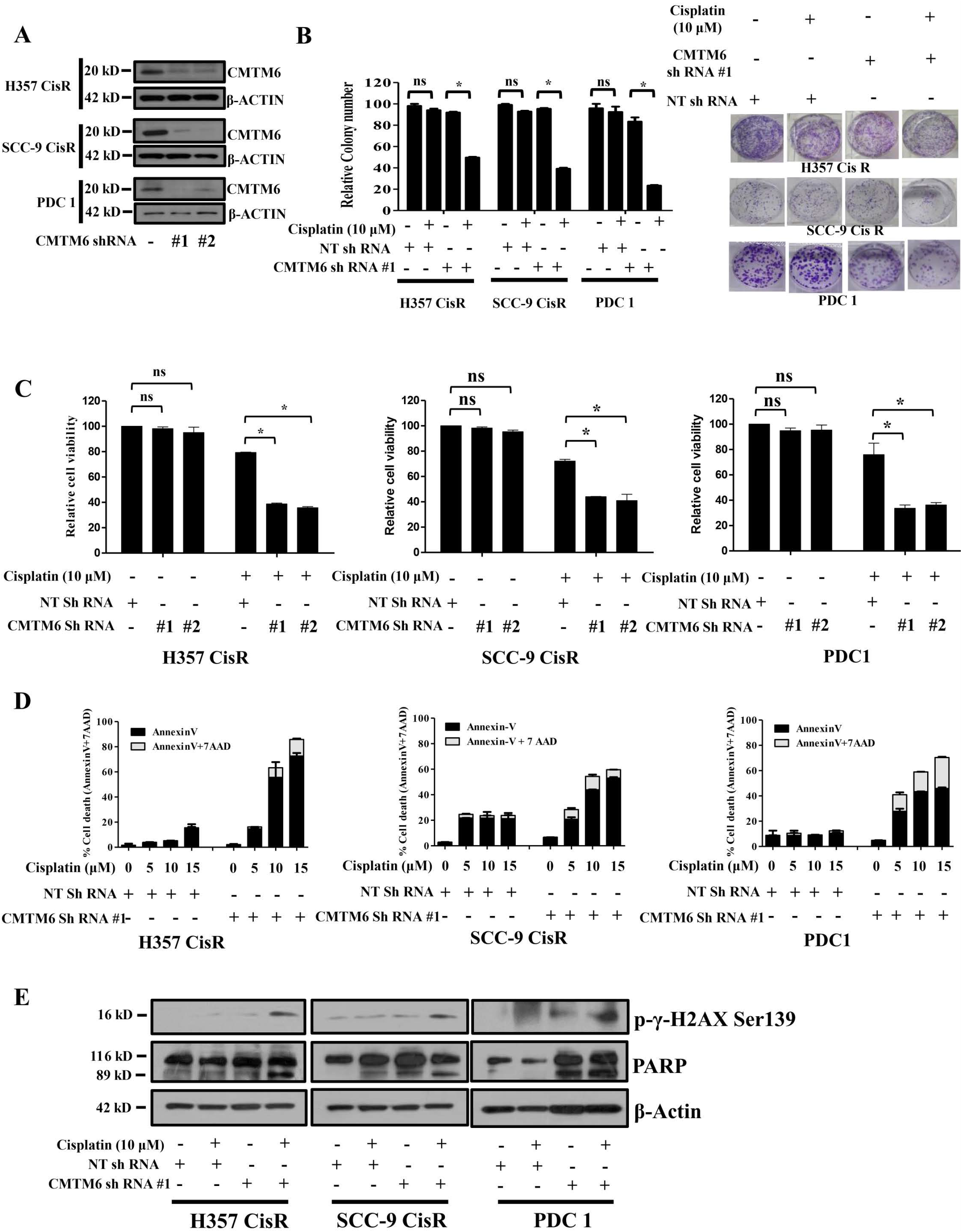
CMTM6 knock down restores cisplatin induced cell death in drug resistant OSCC: **A)** Cisplatin resistant OSCC lines were stable transfected with NTShRNA and CMTM6ShRNA as described in material methods. ShRNA#1 targets CMTM6 mRNA and ShRNA#2 targets 5’UTR of CMTM6 mRNA. Lysates were collected from indicated stable clone and immunoblotting was performed with anti CMTM6 and β-Actin antibodies. **B)** Left panel: Cisplatin resistant cells stably expressing NTShRNA and CMTM6ShRNA were treated with cisplatin for 12 days and colony forming assays as described in method section. Bar diagram indicated the relative colony number (n=3 and *: P < 0.05). Right panel: representating photographs of colony forming assay in each group. **C)** Cisplatin resistant cells stably expressing NTShRNA and CMTM6ShRNA were treated with cisplatin for 48h and cell viability was determined by MTT assay (n=3 and *: P < 0.05). **D)** Cisplatin resistant cells stably expressing NTShRNA and CMTM6ShRNA were treated with cisplatin for 48h and after which cell death was determined by annexin V/7AAD assay using flow cytometer. Bar diagrams indicate the percentage of cell death with respective treated groups (Mean ±SEM, n=2). **E)** Cisplatin resistant cells stably expressing NTShRNA and CMTM6ShRNA were treated with cisplatin for 48h and immunoblotting was performed with indicated antibodies.

### CMTM6 overexpression in CMTM6KD cells rescued the chemoresistant phenotype

To confirm the potential role of CMTM6 in modulating cisplatin, CMTM6 was transiently overexpressed in chemoresistant cells those were stably transfected with CMTM6ShRNA#2 (targets 5’UTR of CMTM6 mRNA)followed by treatment with cisplatin (Fig. 3A). The MTT assay and annexin-V/7AAD staining data suggests that ectopic overexpression of CMTM6 in CMTM6KD drug resistant cells results in rescuing the cisplatin resistant phenotype (Fig. 3B, C). Similarly, transient overexpression of CMTM6 decreases the expression of γ-H2AX in CMTM6 KD cells (Fig. 3D).

**Figure 3:**
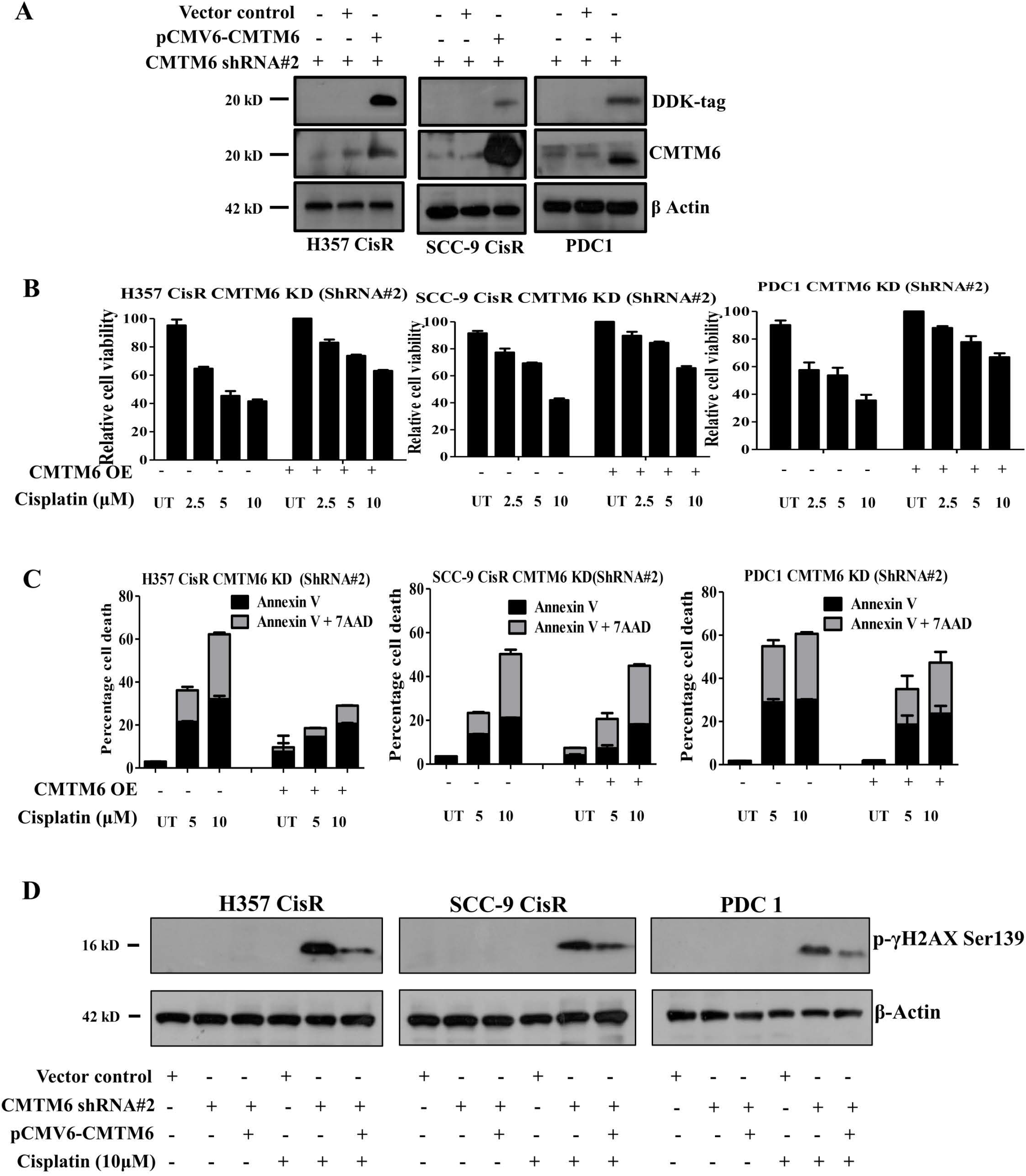
Ectopic overexpression of CMTM6 rescued the drug resistant phenotype in CMTM6KD cells: **A)** For ectopic overexpression, pCMV6-Entry-CMTM6 (MYC-DDK tagged) and control vector were transiently transfected to indicated CMTMKD (ShRNA#2) cells and immunoblotting was performed with indicated antibodies. Efficient overexpression was evident from CMTM6 and DDK expression. **B)** CMTM6 was overexpressed in chemoresistant cells stably expressing CMTM6ShRNA#2 and treated with cisplatin with indicated concentration for 48, after which cell viability was determined by MTT assay (n=3). **C)** Cells were treated as indicated in B panel and cell death was determined by annexin V/7AAD assay using flow cytometer. Bar diagrams indicate the percentage of cell death with respective treated groups (Mean ±SEM, n=2). **D)** CMTM6 was overexpressed in chemoresistant cells stably expressing CMTM6ShRNA#2 followed by cisplatin treatment for 48h and immunoblotting was performed with indicated antibodies.

### CMTM6 regulates AKT/GSK3β signaling by stabilizing membrane Enolase-1 expression

The deregulated proteins, identified from global proteomics analysis, were converted to gene list and a functional analysis was carried out using Ingenuity Pathway Analysis (IPA). This showed deregulation of multiple functional pathways in acquired chemo-resistant cells with significant up-regulation of PI3k/AKT signaling (Fig. 4A). Out of 130 molecules that regulate AKT signaling, in this analysis 30 AKT signaling related genes were deregulated including β-catenin. Henceforth, we evaluated the expression of p-AKT (S437) and p-GSK-3β (S9) in chemoresistant cells. The immunoblotting data suggests that knock down of CMTM6 in chemoresistant cell results in significant reduction of p-AKT and p-GSK3β expression (Fig. 4B). Similarly, CMTM6 overexpression in CMTM6KD chemoresistant cells results in increased p-AKT (S437) and p-GSK-3β (S9) expression (Fig. 4C). Earlier, it is reported in the literature that Enolase-1 can activate AKT signaling ^13^. In a mass spectrometry based analysis by Burr et al., 2017 mentioned that CMTM6 interacts with membrane enolase-1 ^14^. Henceforth, we wanted to observe whether CMTM6 interacts with Enolase-1. Our co-immunoprecipitation and confocal microscopy assay confirmed that CMTM6 and Enolase-1 interact with each other and they co-localized in plasma membrane (Fig. 4D, E). Further, we analyzed the expression of Enolase-1 in cytoplasmic and membrane fraction in chemoresistant cells those stably expressing NTShRNA and CMTM6ShRNA. The data indicates significant reduction in the expression of membrane Enolase-1 in CMTM6KD cells (Fig. 4F).

**Figure 4:**
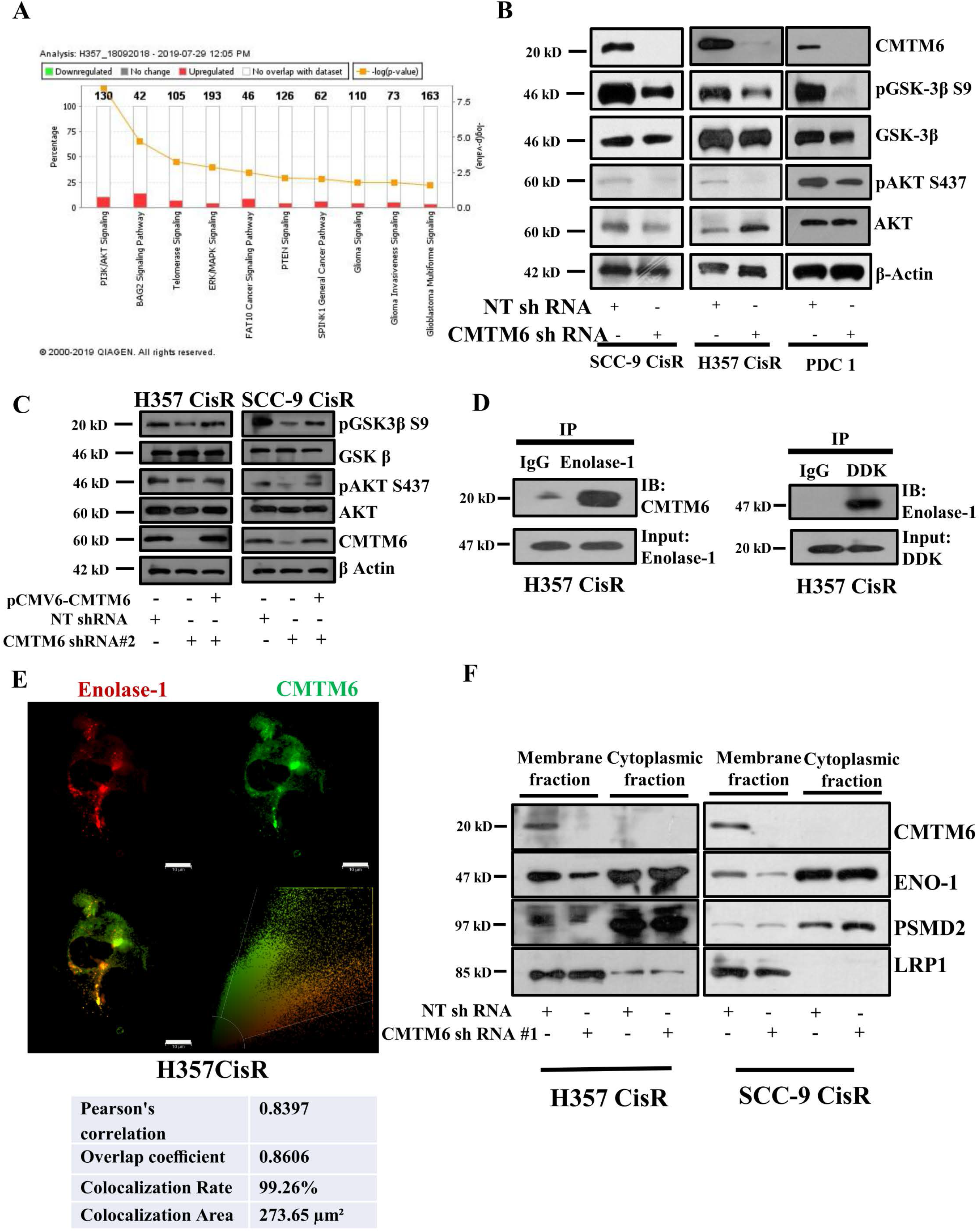
CMTM6 activates AKT/GSK3β signaling by stabilizing membrane enolase-1 expression: **A)** Deregulated genes from proteomic profiling were subjected to Ingenuity Pathway Analysis and list of pathways involved are indicated as bar diagram. **B)** The lysates were isolated and subjected to immunoblotting with indicated antibodies in chemoresistant cells stably expression NTShRNA or CMTM6ShRNA#1. **C)** CMTM6 was overexpressed in chemoresistant cells stably expressing NTShRNA or CMTM6ShRNA#2 and immunoblotting was performed with indicated antibodies. **D)** Left panel: Lysates were isolated from H357CiSR and immunoprecipitated with Enolase-1 and immunoblotting was performed with CMTM6. Right panel: H357CisR was transfected with pCMV6-Entry-CMTM6 (MYC-DDK tagged) after which the lysates were isolated and IP followed by IB was performed with indicated antibodies. **E)** H357CisR cells were subjected to immunostaining with anti-CMTM6 and anti-Enolase-1 using confocal microscope as described in materials and method. **F)** Lysates were isolated from membrane and cytoplasmic fraction and immunoblotting was performed using indicated antibodies.

### CMTM6 modulates cisplatin resistance by regulating β-catenin expression

To know if CMTM6 regulates Wnt signaling, we evaluated the expression of β-catenin in CMTMKD cells. The immunoblotting and immunostaining data indicates that when CMTM6 is knocked down in chemoresistant lines, there is significant downregulation of β-catenin, p-β-catenin (s552) and non-phospho active β-catenin (Fig. 5A, B). It is important to mention here that AKT is known to phosphorylates β-catenin at s552 and this results in translocation of β-catenin from cytoplasm to nucleus ^15^. When Wnt activator lithium chloride (LiCl) was treated with CMTM6KD cells, we observed up regulation of β-catenin and c-Myc expression in chemoresistant cells (Fig. 5C). Similarly, MTT assay data suggests the reversal of chemoresistant phenotype of CMTM6KD cells followed by treatment with Wnt activator lithium chloride (LiCl) (Fig. 5D). Further, we evaluated the expression of β-catenin target genes in CMTM6KD cell. The immunoblotting and qRT-PCR data suggests that in CMTMKD chemoresistant cells, there is a significant down regulation of Wnt target pro-proliferation genes i.e. Cyclin-D, c-Myc and CD44 (Fig. 5E, F). Notably, chemoresistant cells stably transfected with CMTM6ShRNA showed a significantly reduced TOPflash luciferase activity both in presence and absence of LiCl indicating diminished β-catenin/TCF-LEF mediated transcriptional activity (Fig. 5G). Reconstitution of CMTM6 in CMTM6KD cells results in enhanced expression of β-catenin and its target pro-survival genes (Fig. 5H, I). In addition to this, association analysis of CMTM6 m-RNA levels with Wnt target genes (TCF4, LEF1,CD-44, MMP14, ENC1, ID2, PPARA and JAG1) from the cancer genome atlas HNSCC cohort using GEPIA showed a positive correlation (r>0.2) (Supplementary Fig. 3). It is well established from literature that Wnt signaling promotes stemness of cancer cells ^16^. Henceforth, we performed spheroid formation assay and found significantly reduced number of spheroids in cisplatin treated chemoresistant cells stably transfected with CMTM6ShRNA (Fig. 6A). Similarly, the expression of hallmark stem cell markers were found to be reduced in chemoresistant lines stably transfected with CMTM6ShRNA (Fig. 6B). Reconstitution of CMTM6 in CMTM6KD cells results in enhanced expression of hallmark stem cell markers (Fig. 6C). To evaluate if CMTM6 can regulate stem ness in chemoresistant cells, we scored the ALDH activity in CMTM6KD cells. Our data suggests that knock down of CMTM6 results in reduction of ALDH activity in chemoresistant OSCC (Fig. 6D). ABC transporters play important role in acquired chemoresistance in cancer. Henceforth, we measured the expression of ABC transporters in chemoresistant cells stably expressing NTSh or CMTM6ShRNA. The qRT-PCR data indicates reduced expression of major ABC transporters (ABCC1, ABCC2, ABCC3, ABCC4, ABCC5 and ABCG2) in CMTM6 depleted cells. Again, the expression of ABC transporter genes were elevated when we ectopically over expressed CMTM6 in CMTM6 depleted cells (Supplementary Fig. 4A). Association analysis of CMTM6 m-RNA levels with ABC transporters from the cancer genome atlas HNSCC cohort using GEPIA showed a positive correlation (r>0.2) (Supplementary Fig. 4B). Overall, CMTM6 regulates the Wnt signaling and cancer stem ness augmenting chemoresistance in OSCC.

**Figure 5:**
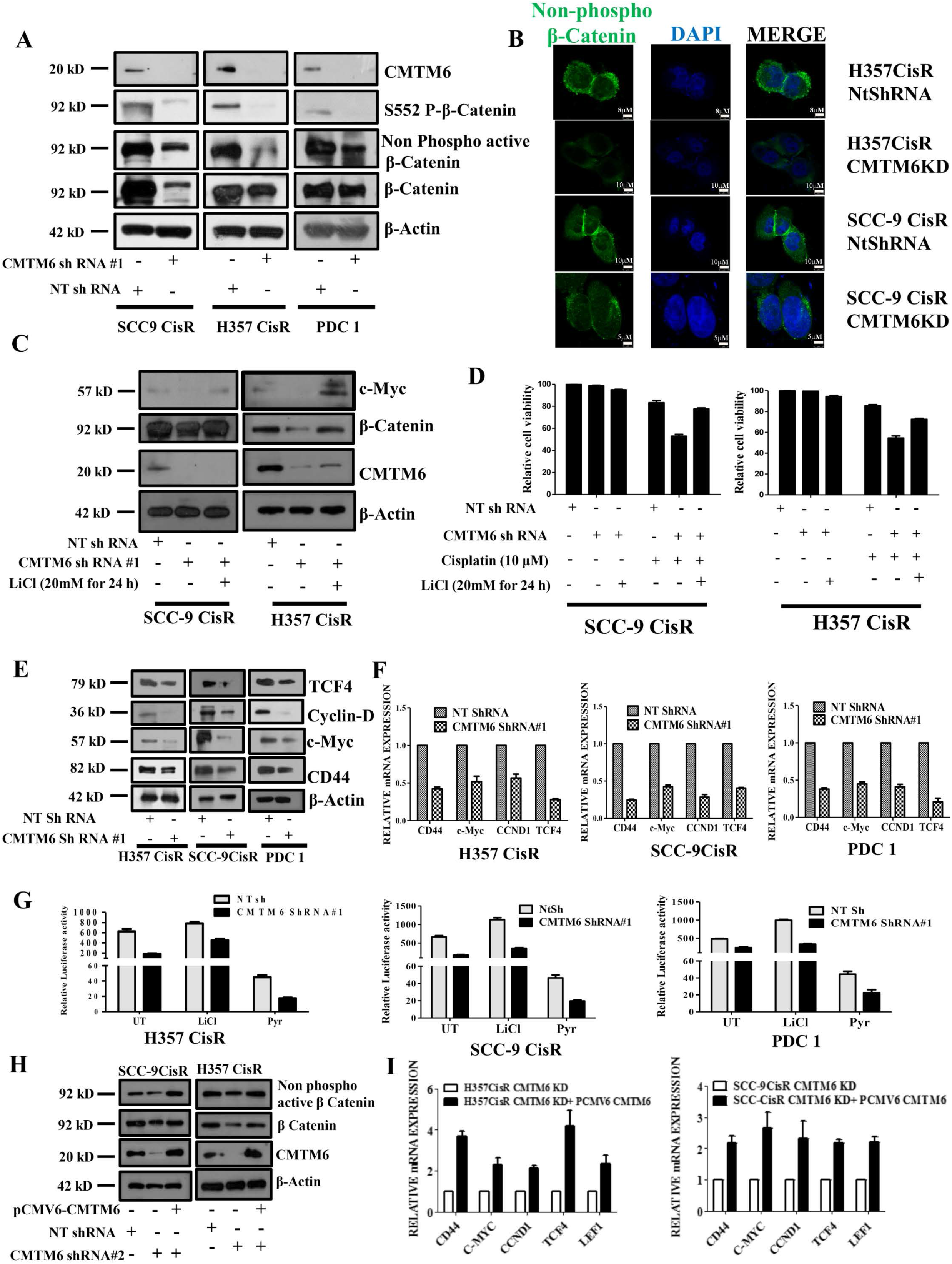
CMTM6 activates Wnt signaling by up regulating β-catenin expression in chemoresistant OSCC. **A)** The lysates were isolated and subjected to immunoblotting with indicated antibodies in chemoresistant cells stably expressing NTShRNA or CMTM6ShRNA#1. **B)** Chemoresistant cells stably expressing NTShRNA or CMTM6ShRNA#1 were subjected to immunostaining and confocal microscopy with indicated antibodies. **C)** Cisplatin resistant cells stably expressing NTShRNA and CMTM6ShRNA#1 were treated with LiCl for 24h and immunoblotting was performed with indicated antibodies **D)** Cisplatin resistant cells stably expressing NTShRNA and CMTM6ShRNA#1 were treated with cisplatin for 48h and LiCl for 24h and cell viability was determined by MTT assay (n=3).**E)** The lysates were isolated and subjected to immunoblotting with indicated antibodies in chemoresistant cells stably expressing NTShRNA or CMTM6ShRNA#1. **F)** Relative mRNA (fold change) expression of indicated genes were analyzed by qRT PCR in indicted cells stably expressing NTShRNA or CMTM6ShRNA#1 (mean ±SEM, n=3). **G)** Chemoresistant cells stably expressing NTShRNA and CMTM6ShRNA#1 were co-transfected with either the TOPflash firefly vector and pRL Renilla control vectors following the treatment with LiCl (20mM) or pyrvinium (50μM) for 12h and luciferase activity was measured as described in materials and methods. The bar diagram indicates the relative luciferase activity in each group (n=3). **H)** CMTM6 was overexpressed in chemoresistant cells stably expressing NTShRNA or CMTM6ShRNA#2 and immunoblotting was performed with indicated antibodies. **I)** CMTM6 was overexpressed in chemoresistant CMTM6KD (ShRNA#2) and relative mRNA expression of indicated genes were determined by qRT PCR (mean ±SEM, n=3).

**Figure 6:**
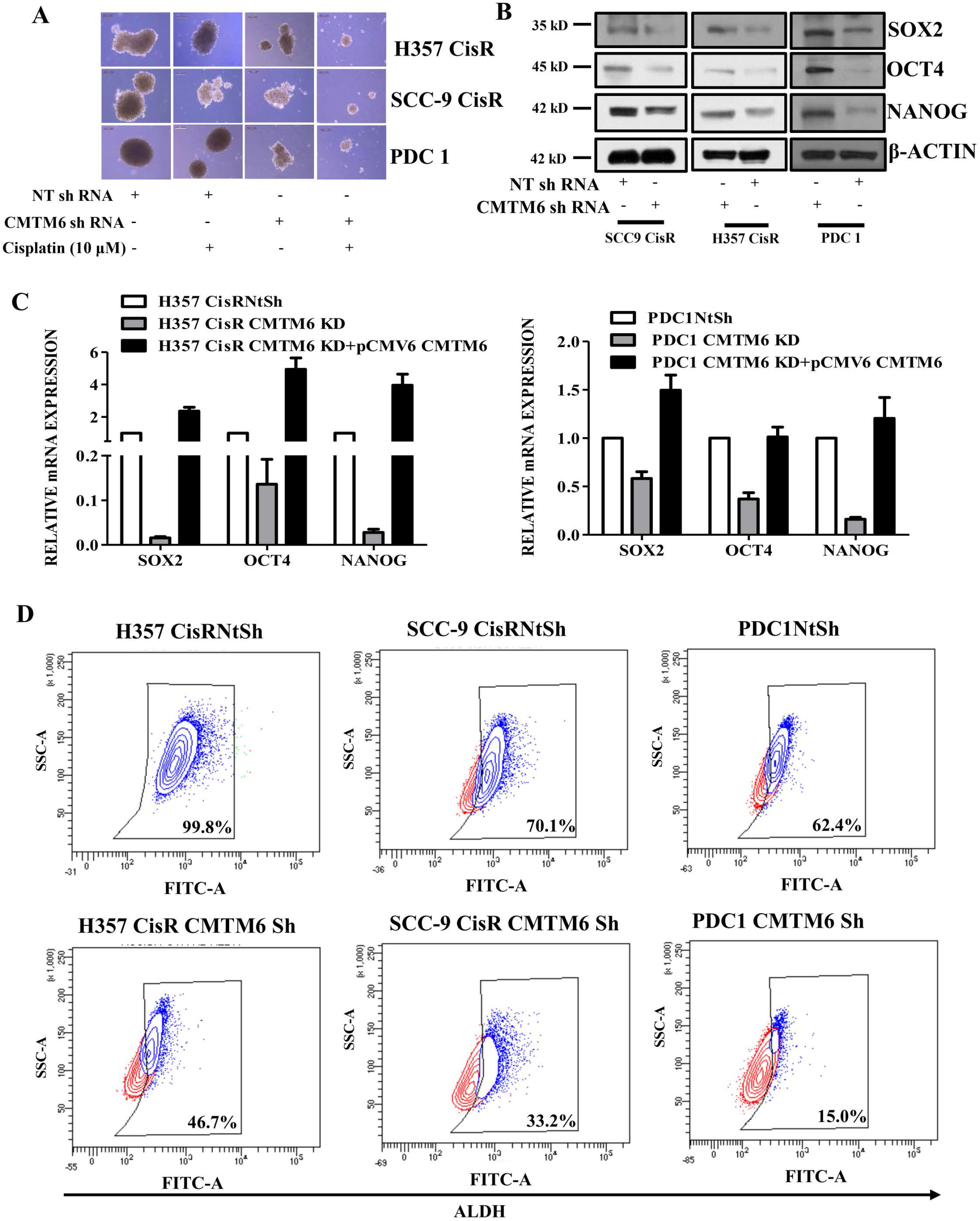
CMTM6 regulates stem ness in chemoresistant cells: **A)** Tumor spheroid assay was performed as described in method section with chemoresistant cells stably expressing NTShRNA and CMTM6ShRNA#1 followed by treatment with indicated concentration of cisplatin for 5 days. At the end of the experiment spheroid photographs were captured using Leica DMIL microscope. **B)** The lysates were isolated and subjected to immunoblotting with indicated antibodies in chemoresistant cells stably expressing NTShRNA or CMTM6ShRNA#1. **C)** CMTM6 was transiently overexpressed in chemoresistant cells stably expressing CMTM6ShRNA#2 and relative mRNA (fold change) expression of indicated genes were analyzed by qRT PCR in indicted cells (mean ±SEM, n=3). **D)** An ALDEFLUOR assay was conducted in chemoresistant cells stably expressing NTShRNA and CMTM6ShRNA#1 and the percentage of ALDH high cells was quantified by flow cytometry as described in method section.

### Knock down of CMTM6 significantly restores cisplatin-mediated cell death in chemoresistant patient-derived xenograft

To evaluate the in vivo efficacy of knocking down CMTM6 in reversing chemoresistance, we generated PDC1 (patient 1 from chemo non-responder group: Table 1) based xenografts using nude mice. Here, we implanted PDC1NtShRNA cells in right upper flank and PDC1CMTM6ShRNA cells in left upper flank of same mice. Treating with cisplatin (3 mg/kg) significantly reduced the tumor burden in case of CMTM6ShRNA group but not in NtShRNA group (Fig. 7 A-C). We also observed significantly decreased cell proliferation in cisplatin treated CMTM6ShRNA tumors with reduced expression of Wnt target pro survival genes (Fig. 7D).

**Figure 7:**
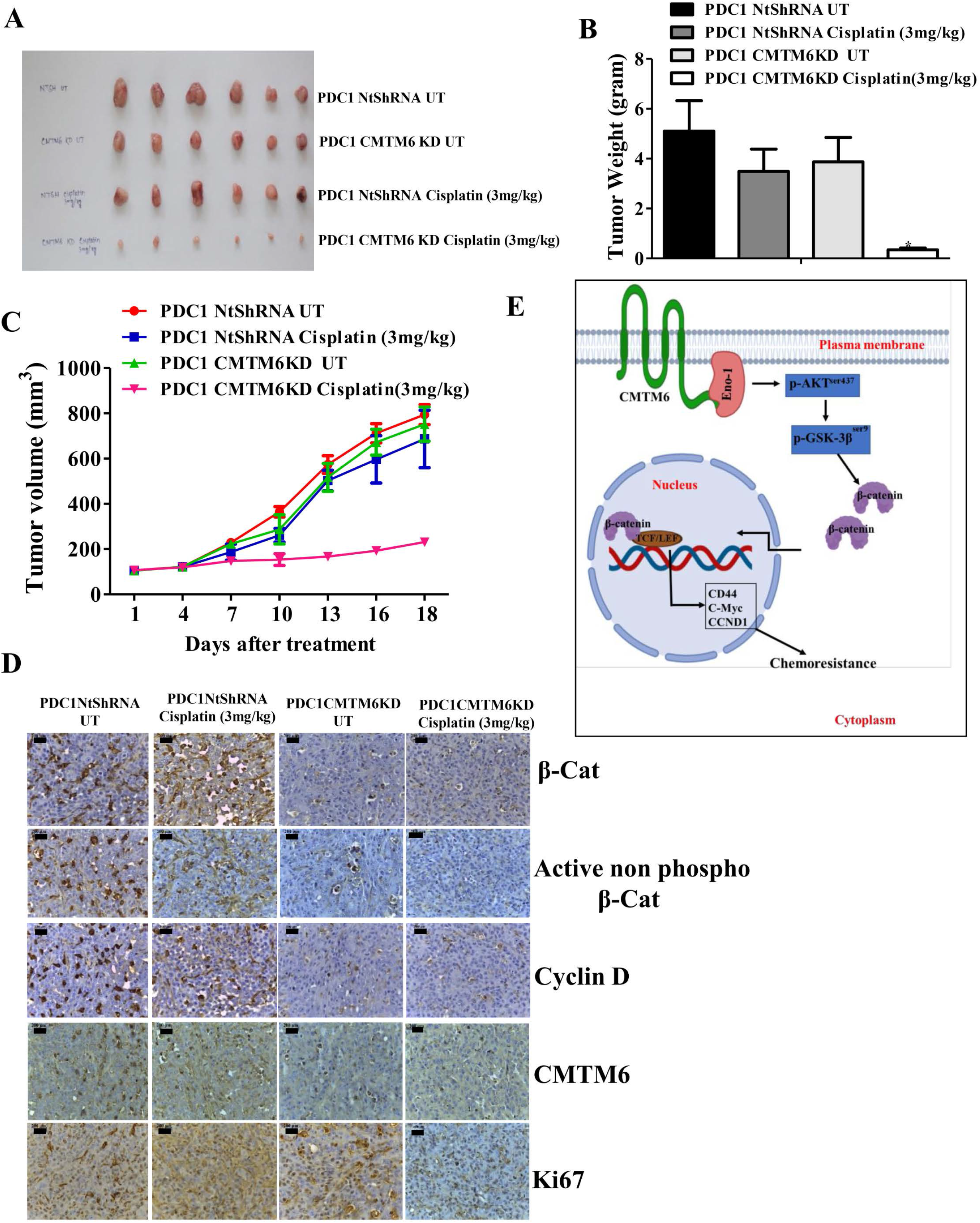
Knock down of CMTM6 reverses cisplatin resistance in chemoresistant xenografts: **A)** Patient derived cells (PDC1) established from tumor of chemo-nonresponder patient. PDC1 cells stably expressing NtShRNA were implanted in right upper flank of athymic male nude mice and PDC1 cells stably expressing CMTM6ShRNA#1 (PDC1 CMTM6KD) were implanted in left upper flank, after which they were treated with cisplatin at indicated concentration. **A)** At the end of the experiment mice were euthanized, tumors were isolated and photographed (n=6). **B)** Bar diagram indicates the tumor weight measured at the end of the experiment (mean ± SEM, *P < 0.05, n = 6). **C)** Tumor growth was measured in indicated time point using digital slide calipers and plotted as a graph (mean ± SEM, n = 6). **D)** After completion of treatment, tumors were isolated and paraffin-embedded sections were prepared as described in materials and methods to perform immunohistochemistry with indicated antibodies. **E)** Schematic presentation of the mechanism by which CMTM6 mediates chemoresistance in OSCC.

## Discussion

Chemotherapy successfully eliminates the rapidly dividing cells in tumor mass but poorly targets the slowly dividing cells. These cells have either inherent resistance properties or they acquired chemoresistance during the treatment of drug. Due to development of chemoresistance, the patient experience continued tumor growth with metastatic disease. The chemoresistance phenotypes can be attributed to reduced apoptosis, enhanced cancer stem cell population, altered metabolic activity and decreased drug accumulation ^6, 17, 18^. All these hallmarks are endpoint events when the tumor cells have already acquired drug resistance. However, the exact causative factors responsible for it are yet to be explored. Till date, most of the study engaged the parental sensitive cells and late drug resistant cells to understand the molecular mechanism for chemoresistance. On the contrary, here in this study, we have performed global proteome profiling of parental sensitive, early and late cisplatin-resistant cells. As per our hypothesis, addition of early resistant group for proteome analysis may enable us to identify the key causative factors responsible for acquired chemoresistance.

Among the set of deregulated proteins in our proteome profiling, CMTM6 was the highest upregulated protein in early and late cisplatin resistant cells as compared to sensitive counterpart. CMTM6 represents the CMTM family member proteins, which consists of eight members ^19^. CMTM6 gene is located at chromosome no 3p22 region ^20^. All the CMTM proteins belong to chemokine-like factor gene superfamily, a superfamily similar to the chemokine and transmembrane 4 superfamilies ^19^. CMTM6 is a type-3 transmembrane protein with a MARVEL domain consisting of three transmembrane helices. It is well established that proteins having MARVEL domain plays important role in regulating the trafficking of transmembrane proteins ^21^. The subcellular localization of CMTM6 is mostly in plasma membrane and it is expressed in several tissues of human body (https://www.proteinatlas.org/ENSG00000091317-CMTM6/tissue). Until 2017, the function of this novel protein was not known. A genome wide CRISPR based screening in pancreatic cancer cell line identified that CMTM6 stabilizes the expression of PD-L1. Interestingly, CMTM6 interacts and co-localized with PD-L1 in plasma membrane. Again, CMTM6 prevents the lysosome mediated degradation of PD-L1, therefore it stabilizes the expression PD-L1 ^14, 22^. Overall, these studies suggest that CMTM6 is an important factor for immune invasion by tumor cells. Further studies suggest that knocking down CMTM6 results in decreased PD-L1 expression and increased infiltration of CD8+ and CD4+ T-cells, that in turn increased the anti-tumor immunity in HNSCC ^23^. CMTM6 expression is also up regulated in high grade malignant glioma and it can be correlated with poor prognosis of glioma patients ^24^. In lung cancer patients, CMTM6 acts as a predictor for PD-1 inhibitor therapy i.e. patients having higher CMTM6 expression, responded well to PD-1 inhibitors ^25, 26^. As the expression of CMTM6 has been detected in several tissues, it is predicted that it might have several biological functions other than triggering immune evasion by tumor cells. Here, in this study for the first time, we uncover another important biological function of CMTM6, i.e. it is a major driver of cisplatin resistance.

In this study, we performed pathway analysis of the set of proteins those were deregulated between sensitive, early and late cisplatin resistant cells. Our data suggests that CMTM6 regulates Wnt signaling through AKT/GSK3β axis. A study by Burr et al in 2017, co-immunoprecipitated CMTM6 from digitonin lysate of pancreatic cancer cells and subjected it to mass spectrometry to explore the potential interacting partners of CMTM6 ^14^. The data suggests that CMTM6 interacts with membrane Enolase-1, but the biological relevance of this interaction was not explored. In this study, the CO-IP and confocal microscopy data suggests that CMTM6 interacts with membrane enolase-1 and co-localized in plasma membrane. We also demonstrate here that knocking down CMTM6 reduced the expression of membrane enolase-1. It is very well documented by other groups that Enolase-1 enhances the phosphorylation of AKT and GSK3β ^13, 27^. Henceforth, we predict that CMTM6 stabilizes the expression of membrane Enolase-1 and activates the AKT/GSK3β mediated Wnt signaling (Fig. 7E). It is very evident from literature that Wnt/β-catenin signaling plays important role in acquiring chemoresistance as it regulates the cancer stemness ^28, 29, 30^. Here, we have also demonstrated that knocking down CMTM6 significantly reduced the stemness properties of chemoresistant cells. Overall in this study, we uncover the novel mechanism by which CMTM6 regulates Wnt signaling and mediates cisplatin resistance in OSCC.

In conclusion, it was earlier established that CMTM6 is a novel protein, which stabilizes PD-L1 and potentiates the immune evasion by tumor cells. Now we for the first time demonstrate that CMTM6 is a major driver of cisplatin resistance. Henceforth, targeting CMTM6 can be a useful strategy to overcome therapy resistance in advanced squamous cell carcinomas.

## Acknowledgment

Grant support: This work is supported by ICMR (5/13/9/2019-NCD-III) and Institute of Life Sciences, Bhubaneswar intramural support. RD is thankful to Ramalingaswami Fellowship. PM is a CSIR-JRF, SM is UGC-JRF, OPS is a UGC-SRF. Core support from International Centre for Genetic Engineering and Biotechnology is highly acknowledged.

## Conflict of interest

The authors have no conflict of interest

## Notes

http://proteomecentral.proteomexchange.org/cgi/GetDataset?ID=PXD016977

