## Supplementary Table 1 for "CMTM6 drives cisplatin resistance in OSCC by regulating AKT mediated Wnt signaling"

| **Sh RNA primers** | **Oligo sequence** |
| --- | --- |
| **pLKO.1 CMTM6 sh RNA F (ShRNA#1)** | **CCGGCTTTCTTCTGAGTCTCCTTATCTCGAGATAAGGAGACTCAGAAGAAAGTTTTTG** |
| **pLKO.1 CMTM6 sh RNA R (ShRNA#1)** | **AATTCAAAAACTTTCTTCTGAGTCTCCTTATCTCGAGATAAGGAGACTCAGAAGAAAG** |
| **pLKO.1 CMTM6 5' UTR sh RNA F (ShRNA#2)** | **CCGGCCCAAGACAGTGAAAGTAATTCTCGAGAATTACTTTCACTGTCTTGGGTTTTTG** |
| **pLKO.1 CMTM6 5' UTR sh RNA R**  **(ShRNA#2)** | **AATTCAAAAACCCAAGACAGTGAAAGTAATTCTCGAGAATTACTTTCACTGTCTTGGG** |

| **qRT PCR Primers** | **Primer sequence** |
| --- | --- |
| **CMTM6 qRT F** | **CGCTGCCTACTTTTTCATGG** |
| **CMTM6 qRT R** | **GAAGAAAGGCACTGCAGCTT** |
| **18S qRT F** | **GTAACCCGTTGAACCCCATT** |
| **18S qRT R** | **CCATCCAATCGGTAGTAGCG** |
| **GAPDH qRT F** | **TCGGAGTCAACGGATTTGGT** |
| **GAPDH qRT R** | **TTGCCATGGGTGGAATCATA** |
| **OCT4 qRT F** | **CGACCATCTGCCGCTTTGAG** |
| **OCT4 qRT R** | **CCCCCTGTCCCCCATTCCTA** |
| **SOX2 qRT F** | **CACCTACAGCATGTCCTACTC** |
| **SOX2 qRT R** | **CATGCTGTTTCTTACTCTCCTC** |
| **Nanog qRT F** | **CAACTGGCCGAAGAATAGCA** |
| **Nanog qRT R** | **GCAGGAGAATTTGGCTGGAA** |
| **CD44 qRT F** | **TGGCACCCGCTATGTCCAG** |
| **CD44 qRT R** | **GTAGCAGGGATTCTGTCTG** |
| **LEF1 qRT F** | **ATCAAGTCTTCCTTGGTGAA** |
| **LEF1 qRT R** | **TATGTACCCGGAATAACTCG** |
| **TCF4 qRT F** | **AGAGCGACAAGCCCCAGAC** |
| **TCF4 qRT R** | **ATTCGCTGCGTCTCCCATC** |
| **CCND1 qRT F** | **TGTGAAGTTCATTTCCAATCC** |
| **CCND1 qRT R** | **GTCACACTTGATCACTCTGG** |
| **c-Myc F** | **CCTGGTGCTCCATGAGGAGAC** |
| **c-Myc R** | **CAGACTCTGACCTTTTGCCAG** |
