## Supplementary Table 2 for "CMTM6 drives cisplatin resistance in OSCC by regulating AKT mediated Wnt signaling"

| Sl No. | Accession | 4R11 | 4R12 | 4R2 | 8R11 | 8R2 | 8R12 | Avg 4M | Avg 8M | FC (4M/8M) | p-value | log2 FC (4M/8M) | -log10 p-value | Gene | Description | Score | Coverage | # Unique Peptides | # AAs | MW [kDa] | calc. pI |
|---|---|---|---|---|---|---|---|---|---|---|---|---|---|---|---|---|---|---|---|---|---|
| 1 | Q8IXM3 | 1.39 | 1.39 | 1.39 | 1.02 | 1.22 | 1.00 | 1.39 | 1.08 | 1.29 | 0.01 | 0.36 | 1.93 | MRPL41 | 39S ribosomal protein L41, mitochondrial OS=Homo sapiens GN=MRPL41 PE=1 SV=1 - [RM41_HUMAN] | 1.72 | 5.11 | 1.00 | 137.00 | 15.37 | 9.57 |
| 2 | Q9NX76 | 1.04 | 1.05 | 1.17 | 1.33 | 1.53 | 1.66 | 1.09 | 1.51 | 0.72 | 0.02 | -0.47 | 1.78 | CMTM6 | CKLF-like MARVEL transmembrane domain-containing protein 6 OS=Homo sapiens GN=CMTM6 PE=1 SV=1 - [CKLF6_HUMAN] | 3.28 | 5.46 | 1.00 | 183.00 | 20.41 | 5.29 |
| 3 | Q5VXV2 | 3.39 | 3.21 | 3.69 | 2.57 | 3.03 | 2.48 | 3.43 | 2.69 | 1.27 | 0.03 | 0.35 | 1.53 | SET | Protein SET OS=Homo sapiens GN=SET PE=2 SV=2 - [Q5VXV2_HUMAN] | 5.67 | 6.72 | 2.00 | 268.00 | 31.25 | 4.21 |
| 4 | B7Z9S8 | 1.26 | 1.21 | 1.28 | 1.09 | 1.19 | 1.12 | 1.25 | 1.13 | 1.10 | 0.04 | 0.14 | 1.40 | | Sodium/potassium-transporting ATPase subunit beta-1 OS=Homo sapiens GN=ATP1B1 PE=2 SV=1 - [B7Z9S8_HUMAN] | 2.42 | 5.67 | 1.00 | 247.00 | 28.61 | 7.59 |
| 5 | P30049 | 1.61 | 1.45 | 1.65 | 1.23 | 1.42 | 1.34 | 1.57 | 1.33 | 1.18 | 0.04 | 0.24 | 1.38 | ATP5F1D | ATP synthase subunit delta, mitochondrial OS=Homo sapiens GN=ATP5D PE=1 SV=2 - [ATPD_HUMAN] | 4.15 | 8.33 | 1.00 | 168.00 | 17.48 | 5.49 |
| 6 | B4DUI5 | 2.00 | 2.46 | 2.49 | 1.76 | 1.97 | 1.61 | 2.32 | 1.78 | 1.30 | 0.05 | 0.38 | 1.33 | | Triosephosphate isomerase OS=Homo sapiens PE=2 SV=1 - [B4DUI5_HUMAN] | 14.39 | 20.19 | 3.00 | 213.00 | 22.86 | 6.92 |
| 7 | P09211 | 1.85 | 1.93 | 2.40 | 1.46 | 1.67 | 1.15 | 2.06 | 1.43 | 1.45 | 0.05 | 0.53 | 1.31 | GSTP1 | Glutathione S-transferase P OS=Homo sapiens GN=GSTP1 PE=1 SV=2 - [GSTP1_HUMAN] | 19.08 | 15.24 | 1.00 | 210.00 | 23.34 | 5.64 |
| 8 | H0YDD8 | 1.82 | 1.82 | 1.79 | 1.55 | 1.76 | 1.57 | 1.81 | 1.63 | 1.11 | 0.06 | 0.15 | 1.26 | RPLP2 | 60S acidic ribosomal protein P2 (Fragment) OS=Homo sapiens GN=RPLP2 PE=4 SV=1 - [H0YDD8_HUMAN] | 8.38 | 48.91 | 2.00 | 92.00 | 9.09 | 4.46 |
| 9 | Q7KZJ3 | 1.72 | 1.66 | 1.84 | 1.34 | 1.63 | 1.46 | 1.74 | 1.48 | 1.18 | 0.06 | 0.23 | 1.23 | | Catenin (Cadherin-associated protein), delta 1 OS=Homo sapiens PE=2 SV=1 - [Q7KZJ3_HUMAN] | 8.88 | 5.90 | 3.00 | 610.00 | 67.96 | 8.13 |
| 10 | Q99714 | 0.94 | 0.63 | 0.79 | 0.53 | 0.59 | 0.54 | 0.79 | 0.55 | 1.42 | 0.06 | 0.51 | 1.21 | HSD17B10 | 3-hydroxyacyl-CoA dehydrogenase type-2 OS=Homo sapiens GN=HSD17B10 PE=1 SV=3 - [HCD2_HUMAN] | 2.75 | 4.60 | 1.00 | 261.00 | 26.91 | 7.78 |
| 11 | Q96T67 | 0.92 | 1.06 | 1.19 | 1.19 | 1.55 | 1.41 | 1.05 | 1.38 | 0.76 | 0.06 | -0.39 | 1.20 | | TOB3 OS=Homo sapiens PE=2 SV=1 - [Q96T67_HUMAN] | 5.54 | 3.98 | 1.00 | 578.00 | 65.07 | 9.33 |
| 12 | P07355 | 1.43 | 1.36 | 1.40 | 1.54 | 1.62 | 1.43 | 1.40 | 1.53 | 0.91 | 0.09 | -0.13 | 1.06 | ANXA2 | Annexin A2 OS=Homo sapiens GN=ANXA2 PE=1 SV=2 - [ANXA2_HUMAN] | 116.60 | 44.84 | 17.00 | 339.00 | 38.58 | 7.75 |
| 13 | Q6E433 | 1.73 | 1.89 | 1.84 | 1.56 | 1.77 | 1.57 | 1.82 | 1.63 | 1.11 | 0.09 | 0.16 | 1.06 | | Activated RNA polymerase II transcription cofactor 4 (Fragment) OS=Homo sapiens PE=2 SV=1 - [Q6E433_HUMAN] | 3.58 | 15.07 | 1.00 | 73.00 | 8.65 | 8.44 |
| 14 | P04083 | 2.45 | 2.32 | 3.17 | 2.08 | 2.11 | 1.92 | 2.64 | 2.04 | 1.30 | 0.09 | 0.38 | 1.05 | ANXA1 | Annexin A1 OS=Homo sapiens GN=ANXA1 PE=1 SV=2 - [ANXA1_HUMAN] | 33.38 | 23.99 | 6.00 | 346.00 | 38.69 | 7.02 |
| 15 | F5H308 | 3.99 | 3.13 | 4.21 | 2.66 | 3.35 | 2.57 | 3.78 | 2.86 | 1.32 | 0.09 | 0.40 | 1.04 | | L-lactate dehydrogenase OS=Homo sapiens GN=LDHA PE=2 SV=1 - [F5H308_HUMAN] | 3.60 | 3.29 | 1.00 | 304.00 | 33.63 | 8.79 |
| 16 | B3GQS7 | 1.73 | 1.71 | 1.91 | 1.60 | 1.70 | 1.52 | 1.79 | 1.61 | 1.11 | 0.10 | 0.15 | 1.01 | HSPD1 | Mitochondrial heat shock 60kD protein 1 variant 1 OS=Homo sapiens GN=HSPD1 PE=2 SV=1 - [B3GQS7_HUMAN] | 80.24 | 35.68 | 18.00 | 569.00 | 60.64 | 6.04 |
| 17 | Q8IUB0 | 1.73 | 1.64 | 2.02 | 1.51 | 1.63 | 1.47 | 1.79 | 1.53 | 1.17 | 0.10 | 0.23 | 0.99 | | Elongation factor 1-alpha OS=Homo sapiens PE=2 SV=1 - [Q8IUB0_HUMAN] | 14.28 | 8.86 | 3.00 | 361.00 | 38.64 | 8.85 |
| 18 | B7Z3V1 | 1.44 | 1.49 | 1.82 | 1.18 | 1.42 | 1.27 | 1.58 | 1.29 | 1.23 | 0.10 | 0.29 | 0.99 | | cDNA FLJ60077, highly similar to Sodium/potassium-transporting ATPase alpha-1 chain (EC 3.6.3.9) (Fragment) OS=Homo sapiens PE=2 SV=1 - | 4.97 | 1.96 | 2.00 | 1020.00 | 112.37 | 5.35 |
| 19 | F8W1A4 | 1.91 | 1.91 | 2.13 | 1.78 | 1.87 | 1.59 | 1.98 | 1.75 | 1.13 | 0.11 | 0.18 | 0.97 | AK2 | Adenylate kinase 2, mitochondrial OS=Homo sapiens GN=AK2 PE=2 SV=1 - [F8W1A4_HUMAN] | 12.69 | 23.28 | 4.00 | 232.00 | 25.61 | 7.83 |
| 20 | B4DMA2 | 2.39 | 2.32 | 2.94 | 2.02 | 2.31 | 1.88 | 2.55 | 2.07 | 1.23 | 0.11 | 0.30 | 0.96 | | cDNA FLJ54023, highly similar to Heat shock protein HSP 90-beta OS=Homo sapiens PE=2 SV=1 - [B4DMA2_HUMAN] | 38.89 | 7.29 | 3.00 | 686.00 | 79.15 | 5.02 |
| 21 | E9PFF2 | 2.41 | 2.47 | 2.66 | 2.12 | 2.46 | 1.96 | 2.52 | 2.18 | 1.15 | 0.11 | 0.21 | 0.94 | TKT | Transketolase OS=Homo sapiens GN=TKT PE=2 SV=1 - [E9PFF2_HUMAN] | 20.86 | 13.47 | 3.00 | 334.00 | 36.43 | 7.77 |
| 22 | Q53HF2 | 2.00 | 1.97 | 2.19 | 1.84 | 2.00 | 1.75 | 2.05 | 1.86 | 1.10 | 0.13 | 0.14 | 0.90 | | Heat shock 70kDa protein 8 isoform 2 variant (Fragment) OS=Homo sapiens PE=2 SV=1 - [Q53HF2_HUMAN] | 36.23 | 15.82 | 4.00 | 493.00 | 53.47 | 5.86 |
| 23 | C9JFR7 | 1.71 | 1.72 | 1.93 | 1.57 | 1.75 | 1.50 | 1.79 | 1.61 | 1.11 | 0.15 | 0.15 | 0.83 | CYCS | Cytochrome c (Fragment) OS=Homo sapiens GN=CYCS PE=2 SV=1 - [C9JFR7_HUMAN] | 102.40 | 49.50 | 5.00 | 101.00 | 11.33 | 9.66 |
| 24 | H0Y7X6 | 1.84 | 1.86 | 2.18 | 1.71 | 1.86 | 1.56 | 1.96 | 1.71 | 1.15 | 0.15 | 0.20 | 0.81 | | Spliceosome RNA helicase DDX39B (Fragment) OS=Homo sapiens GN=DDX39B PE=4 SV=1 - [H0Y7X6_HUMAN] | 1.95 | 8.62 | 3.00 | 325.00 | 38.04 | 8.95 |
| 25 | O15400 | 1.94 | 1.77 | 2.03 | 1.66 | 1.86 | 1.39 | 1.91 | 1.64 | 1.17 | 0.16 | 0.22 | 0.81 | STX7 | Syntaxin-7 OS=Homo sapiens GN=STX7 PE=1 SV=4 - [STX7_HUMAN] | 2.57 | 5.36 | 1.00 | 261.00 | 29.80 | 5.55 |
| 26 | Q9Y2S7 | 1.56 | 1.42 | 1.52 | 1.41 | 1.46 | 1.32 | 1.50 | 1.40 | 1.07 | 0.16 | 0.10 | 0.80 | POLDIP2 | Polymerase delta-interacting protein 2 OS=Homo sapiens GN=POLDIP2 PE=1 SV=1 - [PDIP2_HUMAN] | 1.68 | 2.72 | 1.00 | 368.00 | 42.01 | 8.63 |
| 27 | B7ZW15 | 3.39 | 3.59 | 5.41 | 2.81 | 3.39 | 2.65 | 4.13 | 2.95 | 1.40 | 0.16 | 0.49 | 0.80 | | Putative uncharacterized protein OS=Homo sapiens PE=2 SV=1 - [B7ZW15_HUMAN] | 2.06 | 11.20 | 1.00 | 125.00 | 13.94 | 5.31 |
| 28 | C9J9K3 | 1.55 | 1.61 | 1.82 | 1.50 | 1.58 | 1.42 | 1.66 | 1.50 | 1.11 | 0.16 | 0.15 | 0.79 | RPSA | 40S ribosomal protein SA (Fragment) OS=Homo sapiens GN=RPSA PE=2 SV=1 - [C9J9K3_HUMAN] | 4.20 | 10.61 | 2.00 | 264.00 | 29.49 | 5.25 |
| 29 | H7C144 | 1.78 | 1.60 | 1.93 | 1.52 | 1.71 | 1.34 | 1.77 | 1.52 | 1.16 | 0.16 | 0.22 | 0.79 | ACTN4 | Alpha-actinin-4 (Fragment) OS=Homo sapiens GN=ACTN4 PE=2 SV=1 - [H7C144_HUMAN] | 9.10 | 11.99 | 3.00 | 342.00 | 38.98 | 5.24 |
| 30 | B3KML9 | 2.02 | 1.87 | 2.30 | 1.73 | 1.99 | 1.48 | 2.06 | 1.73 | 1.19 | 0.16 | 0.25 | 0.79 | | cDNA FLJ11352 fis, clone HEMBA1000020, highly similar to Tubulin beta-2C chain OS=Homo sapiens PE=2 SV=1 - [B3KML9_HUMAN] | 52.64 | 14.61 | 5.00 | 397.00 | 44.57 | 4.93 |
| 31 | P62258 | 1.55 | 1.15 | 1.51 | 1.16 | 1.19 | 1.24 | 1.40 | 1.19 | 1.18 | 0.18 | 0.23 | 0.75 | YWHAE | 14-3-3 protein epsilon OS=Homo sapiens GN=YWHAE PE=1 SV=1 - [1433E_HUMAN] | 21.16 | 14.12 | 1.00 | 255.00 | 29.16 | 4.74 |
| 32 | B4DHG0 | 1.80 | 1.75 | 1.82 | 1.70 | 1.79 | 1.59 | 1.79 | 1.69 | 1.06 | 0.18 | 0.08 | 0.75 | DLD | Dihydrolipoyl dehydrogenase, mitochondrial OS=Homo sapiens GN=DLD PE=2 SV=1 - [B4DHG0_HUMAN] | 7.17 | 7.32 | 3.00 | 410.00 | 43.56 | 7.03 |
| 33 | E7EUT5 | 2.07 | 2.10 | 2.60 | 1.91 | 2.14 | 1.72 | 2.26 | 1.92 | 1.17 | 0.19 | 0.23 | 0.73 | GAPDH | Glyceraldehyde-3-phosphate dehydrogenase OS=Homo sapiens GN=GAPDH PE=2 SV=1 - [E7EUT5_HUMAN] | 30.42 | 22.31 | 5.00 | 260.00 | 27.85 | 6.95 |
| 34 | Q09666 | 1.50 | 1.49 | 1.58 | 1.63 | 1.98 | 1.57 | 1.52 | 1.73 | 0.88 | 0.19 | -0.18 | 0.72 | AHNAK | Neuroblast differentiation-associated protein AHNAK OS=Homo sapiens GN=AHNAK PE=1 SV=2 - [AHNK_HUMAN] | 22.16 | 10.54 | 10.00 | 5890.00 | 628.70 | 6.15 |
| 35 | P07195 | 2.27 | 2.25 | 2.54 | 1.95 | 2.34 | 2.06 | 2.36 | 2.12 | 1.11 | 0.19 | 0.15 | 0.72 | LDHB | L-lactate dehydrogenase B chain OS=Homo sapiens GN=LDHB PE=1 SV=2 - [LDHB_HUMAN] | 2.12 | 5.39 | 2.00 | 334.00 | 36.62 | 6.05 |
| 36 | B2R7Y0 | 1.87 | 1.81 | 2.23 | 1.70 | 1.88 | 1.61 | 1.97 | 1.73 | 1.14 | 0.19 | 0.19 | 0.71 | | cDNA, FLJ93654, highly similar to Homo sapiens serpin peptidase inhibitor, clade B (ovalbumin), member 2 (SERPINB2), mRNA OS=Homo sapiens PE=2 SV=1 - [B2R7Y0_HUMAN] | 4.58 | 4.58 | 2.00 | 415.00 | 46.58 | 5.64 |
| 37 | P80723 | 1.83 | 1.67 | 2.03 | 1.63 | 1.76 | 1.61 | 1.84 | 1.67 | 1.11 | 0.20 | 0.15 | 0.71 | BASP1 | Brain acid soluble protein 1 OS=Homo sapiens GN=BASP1 PE=1 SV=2 - [BASP1_HUMAN] | 5.16 | 6.17 | 1.00 | 227.00 | 22.68 | 4.63 |
| 38 | B2R4P2 | 2.35 | 2.51 | 3.18 | 2.10 | 2.54 | 2.02 | 2.68 | 2.22 | 1.21 | 0.20 | 0.27 | 0.70 | | cDNA, FLJ92164, highly similar to Homo sapiens peroxiredoxin 1 (PRDX1), mRNA OS=Homo sapiens PE=2 SV=1 - [B2R4P2_HUMAN] | 5.57 | 10.05 | 2.00 | 199.00 | 22.19 | 8.38 |
| 39 | O14684 | 0.90 | 0.82 | 1.02 | 0.75 | 0.91 | 0.60 | 0.91 | 0.76 | 1.21 | 0.21 | 0.27 | 0.67 | PTGES | Prostaglandin E synthase OS=Homo sapiens GN=PTGES PE=1 SV=2 - [PTGES_HUMAN] | 3.36 | 6.58 | 1.00 | 152.00 | 17.09 | 9.50 |
| 40 | F8VPE8 | 1.57 | 1.53 | 1.77 | 1.43 | 1.60 | 1.42 | 1.62 | 1.48 | 1.09 | 0.22 | 0.13 | 0.67 | RPLP0 | 60S acidic ribosomal protein P0 (Fragment) OS=Homo sapiens GN=RPLP0 PE=2 SV=1 - [F8VPE8_HUMAN] | 11.94 | 7.84 | 1.00 | 153.00 | 16.67 | 9.33 |
| 41 | B7Z800 | 1.63 | 1.60 | 2.22 | 1.48 | 1.63 | 1.44 | 1.82 | 1.52 | 1.20 | 0.23 | 0.26 | 0.65 | ADK | Adenosine kinase OS=Homo sapiens GN=ADK PE=2 SV=1 - [B7Z800_HUMAN] | 2.00 | 3.36 | 1.00 | 327.00 | 36.56 | 6.79 |
| 42 | F8WBR5 | 1.80 | 1.74 | 2.00 | 1.65 | 1.84 | 1.58 | 1.85 | 1.69 | 1.09 | 0.23 | 0.13 | 0.64 | CALM2 | Calmodulin OS=Homo sapiens GN=CALM2 PE=2 SV=1 - [F8WBR5_HUMAN] | 4.83 | 10.77 | 1.00 | 65.00 | 7.37 | 4.01 |
| 43 | H0YGF3 | 1.45 | 1.60 | 1.66 | 1.62 | 1.93 | 1.66 | 1.57 | 1.74 | 0.90 | 0.23 | -0.14 | 0.64 | PRPF19 | Pre-mRNA-processing factor 19 (Fragment) OS=Homo sapiens GN=PRPF19 PE=2 SV=1 - [H0YGF3_HUMAN] | 2.76 | 17.78 | 1.00 | 45.00 | 5.25 | 9.55 |
| 44 | P23284 | 1.62 | 1.52 | 1.71 | 1.49 | 1.61 | 1.37 | 1.61 | 1.49 | 1.08 | 0.23 | 0.11 | 0.63 | PPIB | Peptidyl-prolyl cis-trans isomerase B OS=Homo sapiens GN=PPIB PE=1 SV=2 - [PPIB_HUMAN] | 28.58 | 29.63 | 5.00 | 216.00 | 23.73 | 9.41 |
| 45 | D6RBE9 | 2.35 | 2.23 | 2.01 | 2.20 | 1.95 | 1.75 | 2.20 | 1.97 | 1.12 | 0.24 | 0.16 | 0.63 | ANXA5 | Annexin OS=Homo sapiens GN=ANXA5 PE=2 SV=1 - [D6RBE9_HUMAN] | 4.43 | 3.64 | 1.00 | 220.00 | 24.68 | 4.89 |
| 46 | Q05DN3 | 1.77 | 2.22 | 1.99 | 1.67 | 1.99 | 1.58 | 1.99 | 1.75 | 1.14 | 0.24 | 0.19 | 0.63 | IMMT | IMMT protein (Fragment) OS=Homo sapiens GN=IMMT PE=2 SV=1 - [Q05DN3_HUMAN] | 2.53 | 8.86 | 1.00 | 316.00 | 33.72 | 9.31 |
| 47 | Q96KK5 | 1.21 | 1.15 | 1.38 | 1.34 | 1.56 | 1.28 | 1.25 | 1.39 | 0.90 | 0.25 | -0.16 | 0.60 | HIST1H2AH | Histone H2A type 1-H OS=Homo sapiens GN=HIST1H2AH PE=1 SV=3 - [H2A1H_HUMAN] | 46.82 | 27.34 | 3.00 | 128.00 | 13.90 | 10.89 |
| 48 | D6R904 | 1.31 | 1.30 | 1.35 | 1.35 | 1.51 | 1.33 | 1.32 | 1.40 | 0.95 | 0.26 | -0.08 | 0.59 | TPM3 | Tropomyosin alpha-3 chain OS=Homo sapiens GN=TPM3 PE=2 SV=1 - [D6R904_HUMAN] | 11.20 | 20.00 | 2.00 | 95.00 | 11.01 | 4.79 |
| 49 | A1A5C5 | 1.79 | 1.73 | 1.74 | 1.81 | 2.19 | 1.77 | 1.75 | 1.92 | 0.91 | 0.26 | -0.14 | 0.58 | RRBP1 | RRBP1 protein OS=Homo sapiens GN=RRBP1 PE=2 SV=1 - [A1A5C5_HUMAN] | 2.43 | 1.73 | 1.00 | 751.00 | 84.26 | 5.01 |
| 50 | E5RJD2 | 1.34 | 1.35 | 1.26 | 1.32 | 1.48 | 1.37 | 1.32 | 1.39 | 0.95 | 0.27 | -0.08 | 0.57 | DECR1 | 2,4-dienoyl-CoA reductase, mitochondrial (Fragment) OS=Homo sapiens GN=DECR1 PE=2 SV=1 - [E5RJD2_HUMAN] | 4.38 | 8.61 | 1.00 | 151.00 | 15.89 | 8.85 |
| 51 | Q59GT8 | 1.82 | 1.91 | 2.40 | 1.68 | 2.05 | 1.39 | 2.04 | 1.71 | 1.20 | 0.27 | 0.26 | 0.57 | | BM-010 variant (Fragment) OS=Homo sapiens PE=2 SV=1 - [Q59GT8_HUMAN] | 2.51 | 23.88 | 1.00 | 67.00 | 7.70 | 5.35 |
| 52 | P13639 | 2.27 | 2.00 | 2.78 | 2.01 | 2.25 | 1.79 | 2.35 | 2.01 | 1.17 | 0.28 | 0.22 | 0.56 | EEF2 | Elongation factor 2 OS=Homo sapiens GN=EEF2 PE=1 SV=4 - [EF2_HUMAN] | 13.61 | 3.15 | 3.00 | 858.00 | 95.28 | 6.83 |
| 53 | H0YCG2 | 1.43 | 1.28 | 1.45 | 1.46 | 1.63 | 1.39 | 1.39 | 1.50 | 0.93 | 0.28 | -0.11 | 0.55 | LAMP2 | Lysosome-associated membrane glycoprotein 2 (Fragment) OS=Homo sapiens GN=LAMP2 PE=4 SV=1 - [H0YCG2_HUMAN] | 2.26 | 2.71 | 1.00 | 258.00 | 28.25 | 6.52 |
| 54 | G3V576 | 1.27 | 1.30 | 1.48 | 1.40 | 1.53 | 1.41 | 1.35 | 1.45 | 0.93 | 0.28 | -0.10 | 0.55 | HNRNPC | Heterogeneous nuclear ribonucleoproteins C1/C2 OS=Homo sapiens GN=HNRNPC PE=4 SV=1 - [G3V576_HUMAN] | 19.76 | 15.58 | 4.00 | 231.00 | 25.24 | 9.82 |
| 55 | B4DIW2 | 1.86 | 1.79 | 2.05 | 1.67 | 1.95 | 1.55 | 1.90 | 1.72 | 1.10 | 0.28 | 0.14 | 0.55 | | cDNA FLJ54035, highly similar to Neutral alpha-glucosidase AB OS=Homo sapiens PE=2 SV=1 - [B4DIW2_HUMAN] | 11.87 | 4.10 | 3.00 | 830.00 | 93.93 | 5.87 |
| 56 | A8K5W7 | 1.55 | 1.78 | 1.99 | 1.67 | 1.69 | 1.31 | 1.77 | 1.56 | 1.14 | 0.29 | 0.19 | 0.54 | | cDNA FLJ75180, highly similar to Homo sapiens mitochondrial isoleucine tRNA synthetase, mRNA OS=Homo sapiens PE=2 SV=1 - | 2.43 | 1.28 | 1.00 | 940.00 | 105.87 | 6.42 |
| 57 | P07737 | 2.04 | 2.26 | 2.89 | 1.92 | 2.27 | 2.01 | 2.40 | 2.06 | 1.16 | 0.29 | 0.22 | 0.53 | PFN1 | Profilin-1 OS=Homo sapiens GN=PFN1 PE=1 SV=2 - [PROF1_HUMAN] | 4.05 | 11.43 | 1.00 | 140.00 | 15.04 | 8.27 |
| 58 | Q08211 | 1.52 | 1.43 | 1.85 | 1.45 | 1.53 | 1.33 | 1.60 | 1.43 | 1.12 | 0.30 | 0.16 | 0.53 | DHX9 | ATP-dependent RNA helicase A OS=Homo sapiens GN=DHX9 PE=1 SV=4 - [DHX9_HUMAN] | 7.91 | 2.36 | 2.00 | 1270.00 | 140.87 | 6.84 |
| 59 | J3QSA3 | 1.39 | 1.67 | 1.87 | 1.41 | 1.56 | 1.44 | 1.64 | 1.47 | 1.12 | 0.30 | 0.16 | 0.52 | UBB | Ubiquitin (Fragment) OS=Homo sapiens GN=UBB PE=4 SV=1 - [J3QSA3_HUMAN] | 3.07 | 37.21 | 1.00 | 43.00 | 4.85 | 5.19 |
| 60 | B2R4C1 | 1.30 | 1.17 | 1.36 | 1.27 | 1.39 | 1.51 | 1.28 | 1.39 | 0.92 | 0.30 | -0.12 | 0.52 | | cDNA, FLJ92036, highly similar to Homo sapiens ribosomal protein L31 (RPL31), mRNA OS=Homo sapiens PE=2 SV=1 - [B2R4C1_HUMAN] | 2.29 | 15.20 | 2.00 | 125.00 | 14.49 | 10.54 |
| 61 | B4E2Z3 | 1.61 | 1.53 | 1.79 | 1.47 | 1.68 | 1.34 | 1.64 | 1.50 | 1.10 | 0.30 | 0.13 | 0.52 | | cDNA FLJ54090, highly similar to 4F2 cell-surface antigen heavy chain OS=Homo sapiens PE=2 SV=1 - [B4E2Z3_HUMAN] | 38.57 | 14.48 | 5.00 | 511.00 | 55.90 | 5.17 |
| 62 | P18859 | 1.61 | 1.49 | 1.50 | 1.63 | 1.58 | 1.55 | 1.53 | 1.59 | 0.97 | 0.31 | -0.05 | 0.51 | ATP5PF | ATP synthase-coupling factor 6, mitochondrial OS=Homo sapiens GN=ATP5J PE=1 SV=1 - [ATP5J_HUMAN] | 4.75 | 17.59 | 1.00 | 108.00 | 12.58 | 9.52 |
| 63 | F8VVB9 | 2.30 | 2.15 | 3.03 | 2.00 | 2.49 | 1.81 | 2.49 | 2.10 | 1.19 | 0.32 | 0.25 | 0.50 | TUBA1B | Tubulin alpha-1B chain (Fragment) OS=Homo sapiens GN=TUBA1B PE=2 SV=1 - [F8VVB9_HUMAN] | 46.39 | 11.34 | 2.00 | 247.00 | 27.53 | 5.20 |
| 64 | B3KRY5 | 1.82 | 1.83 | 1.77 | 1.65 | 1.88 | 1.55 | 1.81 | 1.69 | 1.07 | 0.32 | 0.09 | 0.50 | | cDNA FLJ35087 fis, clone PLACE6005546, highly similar to Polymerase I and transcript release factor OS=Homo sapiens PE=2 SV=1 - | 17.71 | 7.83 | 2.00 | 345.00 | 39.02 | 6.51 |
| 65 | F8VYN5 | 1.31 | 1.50 | 1.53 | 1.56 | 1.59 | 1.46 | 1.44 | 1.53 | 0.94 | 0.32 | -0.09 | 0.50 | HNRNPA1 | Heterogeneous nuclear ribonucleoprotein A1 (Fragment) OS=Homo sapiens GN=HNRNPA1 PE=2 SV=1 - [F8VYN5_HUMAN] | 3.14 | 8.85 | 1.00 | 113.00 | 12.89 | 7.64 |
| 66 | D6R9P3 | 1.94 | 1.86 | 2.10 | 1.91 | 1.95 | 1.64 | 1.97 | 1.83 | 1.07 | 0.32 | 0.10 | 0.49 | HNRNPAB | Heterogeneous nuclear ribonucleoprotein A/B OS=Homo sapiens GN=HNRNPAB PE=2 SV=1 - [D6R9P3_HUMAN] | 2.04 | 7.50 | 2.00 | 280.00 | 30.28 | 7.91 |
| 67 | P27348 | 2.20 | 2.53 | 3.00 | 1.75 | 2.55 | 2.32 | 2.57 | 2.20 | 1.17 | 0.33 | 0.22 | 0.48 | YWHAQ | 14-3-3 protein theta OS=Homo sapiens GN=YWHAQ PE=1 SV=1 - [1433T_HUMAN] | 21.16 | 15.92 | 1.00 | 245.00 | 27.75 | 4.78 |
| 68 | B4DE78 | 1.47 | 1.27 | 1.43 | 1.13 | 1.50 | 0.98 | 1.39 | 1.20 | 1.15 | 0.33 | 0.20 | 0.48 | | cDNA FLJ52141, highly similar to 14-3-3 protein gamma OS=Homo sapiens PE=2 SV=1 - [B4DE78_HUMAN] | 24.36 | 28.50 | 2.00 | 207.00 | 23.49 | 4.82 |
| 69 | B4DXF1 | 1.46 | 1.37 | 1.59 | 1.33 | 1.52 | 1.21 | 1.47 | 1.35 | 1.09 | 0.33 | 0.13 | 0.47 | | cDNA FLJ57223 OS=Homo sapiens PE=2 SV=1 - [B4DXF1_HUMAN] | 2.31 | 8.97 | 1.00 | 156.00 | 16.79 | 7.72 |
| 70 | H6VRG2 | 1.73 | 1.62 | 1.80 | 1.90 | 3.02 | 1.63 | 1.72 | 2.18 | 0.79 | 0.34 | -0.34 | 0.47 | KRT1 | Keratin 1 OS=Homo sapiens GN=KRT1 PE=3 SV=1 - [H6VRG2_HUMAN] | 28.67 | 8.70 | 3.00 | 644.00 | 66.03 | 8.12 |
| 71 | B3KP09 | 2.01 | 1.92 | 1.83 | 1.57 | 2.06 | 1.56 | 1.92 | 1.73 | 1.11 | 0.34 | 0.15 | 0.47 | NSUN2 | cDNA FLJ30894 fis, clone FEBRA2005416, highly similar to Homo sapiens NOL1/NOP2/Sun domain family, member 2 (NSUN2), mRNA OS=Homo sapiens PE=2 SV=1 - [B3KP09_HUMAN] | 2.71 | 3.77 | 2.00 | 531.00 | 59.33 | 6.55 |
| 72 | A4UCT0 | 1.58 | 1.35 | 1.33 | 1.39 | 1.31 | 1.29 | 1.42 | 1.33 | 1.07 | 0.34 | 0.09 | 0.46 | | Fructose-bisphosphate aldolase (Fragment) OS=Homo sapiens PE=2 SV=1 - [A4UCT0_HUMAN] | 3.18 | 16.39 | 1.00 | 122.00 | 13.39 | 6.77 |
| 73 | Q5T8M8 | 1.98 | 1.60 | 2.03 | 1.75 | 1.84 | 1.43 | 1.87 | 1.67 | 1.12 | 0.35 | 0.16 | 0.46 | | Actin, alpha skeletal muscle OS=Homo sapiens GN=ACTA1 PE=4 SV=1 - [Q5T8M8_HUMAN] | 596.32 | 35.19 | 2.00 | 287.00 | 32.03 | 5.41 |
| 74 | B4DSH1 | 1.30 | 1.16 | 1.20 | 1.15 | 1.43 | 1.40 | 1.22 | 1.32 | 0.92 | 0.35 | -0.12 | 0.46 | | cDNA FLJ51295, highly similar to Cell division cycle 5-like protein OS=Homo sapiens PE=2 SV=1 - [B4DSH1_HUMAN] | 3.05 | 2.45 | 1.00 | 775.00 | 89.17 | 8.54 |

| | | | | | | | | | | | | | | | | | | | | | |
|---|---|---|---|---|---|---|---|---|---|---|---|---|---|---|---|---|---|---|---|---|---|
| 75 | Q8WUW7 | 1.94 | 1.90 | 2.29 | 1.87 | 2.07 | 1.63 | 2.04 | 1.86 | 1.10 | 0.36 | 0.14 | 0.45 | PKM2 | Pyruvate kinase (Fragment) OS=Homo sapiens GN=PKM2 PE=2 SV=2 - [Q8WUW7_HUMAN] | 9.29 | 9.04 | 3.00 | 343.00 | 37.25 | 8.22 |
| 76 | B4DZZ0 | 1.16 | 1.24 | 1.09 | 1.24 | 0.96 | 0.95 | 1.16 | 1.05 | 1.10 | 0.36 | 0.14 | 0.45 | | cDNA FLJ52128, highly similar to PRA1 family protein 3 OS=Homo sapiens PE=2 SV=1 - [B4DZZ0_HUMAN] | 4.60 | 12.12 | 1.00 | 165.00 | 19.19 | 9.77 |
| 77 | H7C125 | 1.53 | 1.48 | 1.74 | 1.48 | 1.78 | 2.08 | 1.58 | 1.78 | 0.89 | 0.37 | -0.17 | 0.44 | RAB2A | Ras-related protein Rab-2A (Fragment) OS=Homo sapiens GN=RAB2A PE=4 SV=1 - [H7C125_HUMAN] | 4.08 | 13.46 | 1.00 | 104.00 | 12.14 | 5.08 |
| 78 | B4DVQ0 | 1.24 | 1.37 | 1.45 | 1.32 | 1.66 | 1.43 | 1.35 | 1.47 | 0.92 | 0.37 | -0.12 | 0.44 | | cDNA FLJ58286, highly similar to Actin, cytoplasmic 2 OS=Homo sapiens PE=2 SV=1 - [B4DVQ0_HUMAN] | 604.42 | 33.93 | 2.00 | 333.00 | 37.32 | 5.71 |
| 79 | P37802 | 1.94 | 1.66 | 2.23 | 1.65 | 1.99 | 1.54 | 1.94 | 1.73 | 1.12 | 0.37 | 0.17 | 0.43 | TAGLN2 | Transgelin-2 OS=Homo sapiens GN=TAGLN2 PE=1 SV=3 - [TAGL2_HUMAN] | 6.81 | 6.03 | 1.00 | 199.00 | 22.38 | 8.25 |
| 80 | Q9UHS8 | 1.44 | 1.46 | 1.68 | 1.52 | 1.39 | 1.40 | 1.52 | 1.44 | 1.06 | 0.38 | 0.09 | 0.42 | | PRO1975 OS=Homo sapiens PE=2 SV=1 - [Q9UHS8_HUMAN] | 4.59 | 2.54 | 1.00 | 393.00 | 44.12 | 9.03 |
| 81 | F8WCJ1 | 1.75 | 1.81 | 2.19 | 1.72 | 1.96 | 1.50 | 1.91 | 1.73 | 1.11 | 0.38 | 0.15 | 0.42 | EIF5A2 | Eukaryotic translation initiation factor 5A-2 OS=Homo sapiens GN=EIF5A2 PE=2 SV=1 - [F8WCJ1_HUMAN] | 2.02 | 11.43 | 1.00 | 105.00 | 11.68 | 9.14 |
| 82 | B4DG62 | 1.58 | 1.43 | 1.65 | 1.47 | 1.56 | 1.40 | 1.55 | 1.48 | 1.05 | 0.38 | 0.07 | 0.42 | | cDNA FLJ56506, highly similar to Hexokinase-1 (EC 2.7.1.1) OS=Homo sapiens PE=2 SV=1 - [B4DG62_HUMAN] | 5.73 | 1.97 | 2.00 | 915.00 | 102.25 | 6.80 |
| 83 | B4DSW9 | 1.91 | 1.99 | 2.32 | 1.76 | 2.11 | 1.89 | 2.08 | 1.92 | 1.08 | 0.39 | 0.11 | 0.41 | | cDNA FLJ59415, highly similar to Beta-catenin OS=Homo sapiens PE=2 SV=1 - [B4DSW9_HUMAN] | 2.82 | 4.23 | 3.00 | 709.00 | 77.47 | 6.38 |
| 84 | B7Z5V2 | 1.84 | 1.72 | 2.25 | 1.68 | 1.94 | 1.65 | 1.94 | 1.76 | 1.10 | 0.39 | 0.14 | 0.41 | | cDNA FLJ54141, highly similar to Ezrin OS=Homo sapiens PE=2 SV=1 - [B7Z5V2_HUMAN] | 5.88 | 5.96 | 4.00 | 554.00 | 65.53 | 5.91 |
| 85 | G3V2D2 | 1.46 | 1.47 | 1.52 | 1.32 | 1.58 | 1.30 | 1.49 | 1.40 | 1.06 | 0.39 | 0.09 | 0.41 | SHMT2 | Serine hydroxymethyltransferase, mitochondrial OS=Homo sapiens GN=SHMT2 PE=2 SV=1 - [G3V2D2_HUMAN] | 4.72 | 26.92 | 1.00 | 52.00 | 5.77 | 6.74 |
| 86 | Q6NVV1 | 1.80 | 1.65 | 1.83 | 1.70 | 1.74 | 1.66 | 1.76 | 1.70 | 1.03 | 0.40 | 0.05 | 0.40 | RPL13AP3 | Putative 60S ribosomal protein L13a-like MGC87657 OS=Homo sapiens PE=5 SV=1 - [R13AX_HUMAN] | 1.64 | 7.84 | 1.00 | 102.00 | 12.13 | 10.76 |
| 87 | B4E0S6 | 1.58 | 1.36 | 1.51 | 1.43 | 1.66 | 1.61 | 1.48 | 1.57 | 0.94 | 0.40 | -0.09 | 0.40 | | cDNA FLJ55635, highly similar to pre-mRNA-splicing factorATP-dependent RNA helicase DHX15 (EC 3.6.1.-) OS=Homo sapiens PE=2 SV=1 - | 2.39 | 1.28 | 1.00 | 784.00 | 89.49 | 7.46 |
| 88 | Q9BTI9 | 1.89 | 1.95 | 2.09 | 1.82 | 2.06 | 1.74 | 1.98 | 1.87 | 1.06 | 0.40 | 0.08 | 0.40 | NPM1 | NPM1 protein (Fragment) OS=Homo sapiens GN=NPM1 PE=2 SV=2 - [Q9BTI9_HUMAN] | 57.32 | 31.14 | 7.00 | 228.00 | 25.03 | 4.86 |
| 89 | P26599 | 1.43 | 1.40 | 1.59 | 1.35 | 1.53 | 1.29 | 1.47 | 1.39 | 1.06 | 0.40 | 0.09 | 0.40 | PTBP1 | Polypyrimidine tract-binding protein 1 OS=Homo sapiens GN=PTBP1 PE=1 SV=1 - [PTBP1_HUMAN] | 38.11 | 10.55 | 5.00 | 531.00 | 57.19 | 9.17 |
| 90 | P22626 | 1.23 | 1.17 | 1.30 | 1.12 | 1.19 | 1.24 | 1.23 | 1.18 | 1.04 | 0.40 | 0.06 | 0.40 | HNRNPA2B1 | Heterogeneous nuclear ribonucleoproteins A2/B1 OS=Homo sapiens GN=HNRNPA2B1 PE=1 SV=2 - [ROA2_HUMAN] | 28.24 | 19.55 | 6.00 | 353.00 | 37.41 | 8.95 |
| 91 | P27824 | 1.73 | 1.59 | 1.96 | 1.64 | 1.75 | 1.55 | 1.76 | 1.65 | 1.07 | 0.40 | 0.10 | 0.39 | CANX | Calnexin OS=Homo sapiens GN=CANX PE=1 SV=2 - [CALX_HUMAN] | 22.75 | 7.09 | 4.00 | 592.00 | 67.53 | 4.60 |
| 92 | Q8TAS0 | 1.32 | 1.18 | 1.61 | 1.43 | 1.97 | 1.35 | 1.37 | 1.58 | 0.87 | 0.41 | -0.21 | 0.38 | | ATP synthase subunit gamma (Fragment) OS=Homo sapiens PE=2 SV=1 - [Q8TAS0_HUMAN] | 1.62 | 4.47 | 1.00 | 291.00 | 32.23 | 9.11 |
| 93 | B4DUQ1 | 1.72 | 1.35 | 1.64 | 1.44 | 1.59 | 1.29 | 1.57 | 1.44 | 1.09 | 0.41 | 0.12 | 0.38 | | cDNA FLJ54552, highly similar to Heterogeneous nuclear ribonucleoprotein K OS=Homo sapiens PE=2 SV=1 - [B4DUQ1_HUMAN] | 9.92 | 14.58 | 4.00 | 439.00 | 48.48 | 5.92 |
| 94 | A6XGL3 | 1.58 | 1.40 | 1.28 | 1.42 | 1.66 | 1.49 | 1.42 | 1.52 | 0.93 | 0.42 | -0.10 | 0.38 | PRSS1 | Protease serine 1 OS=Homo sapiens PE=2 SV=1 - [A6XGL3_HUMAN] | 5.33 | 7.59 | 1.00 | 237.00 | 25.38 | 7.55 |
| 95 | M0R221 | 1.89 | 1.70 | 2.03 | 1.67 | 1.95 | 1.62 | 1.87 | 1.75 | 1.07 | 0.42 | 0.10 | 0.38 | SNRPA | U1 small nuclear ribonucleoprotein A (Fragment) OS=Homo sapiens GN=SNRPA PE=4 SV=1 - [M0R221_HUMAN] | 2.98 | 9.15 | 1.00 | 142.00 | 15.94 | 10.11 |
| 96 | M0R0N3 | 1.60 | 1.44 | 1.53 | 1.27 | 1.63 | 1.35 | 1.52 | 1.42 | 1.07 | 0.42 | 0.10 | 0.38 | HNRNPM | Heterogeneous nuclear ribonucleoprotein M (Fragment) OS=Homo sapiens GN=HNRNPM PE=4 SV=1 - [M0R0N3_HUMAN] | 8.21 | 20.65 | 2.00 | 276.00 | 30.12 | 8.72 |
| 97 | E9PPU1 | 2.10 | 1.92 | 2.46 | 1.80 | 2.23 | 1.90 | 2.16 | 1.97 | 1.09 | 0.42 | 0.13 | 0.37 | RPS3 | 40S ribosomal protein S3 OS=Homo sapiens GN=RPS3 PE=2 SV=1 - [E9PPU1_HUMAN] | 2.70 | 10.13 | 1.00 | 158.00 | 17.40 | 9.50 |
| 98 | B4E091 | 1.21 | 0.99 | 1.08 | 1.12 | 1.39 | 1.09 | 1.09 | 1.20 | 0.91 | 0.42 | -0.13 | 0.37 | | cDNA FLJ55438, highly similar to Splicing factor 3 subunit 1 OS=Homo sapiens PE=2 SV=1 - [B4E091_HUMAN] | 5.16 | 2.32 | 1.00 | 690.00 | 77.36 | 5.16 |
| 99 | Q7Z612 | 1.67 | 1.72 | 2.03 | 1.59 | 1.86 | 1.58 | 1.81 | 1.68 | 1.08 | 0.43 | 0.11 | 0.37 | | Acidic ribosomal phosphoprotein P1 OS=Homo sapiens PE=2 SV=1 - [Q7Z612_HUMAN] | 17.60 | 14.16 | 1.00 | 113.00 | 11.39 | 4.36 |
| 100 | Q8TCT9 | 1.15 | 1.27 | 1.25 | 1.19 | 1.29 | 1.36 | 1.23 | 1.28 | 0.96 | 0.43 | -0.06 | 0.36 | HM13 | Minor histocompatibility antigen H13 OS=Homo sapiens GN=HM13 PE=1 SV=1 - [HM13_HUMAN] | 2.65 | 3.18 | 1.00 | 377.00 | 41.46 | 6.43 |
| 101 | Q9HCY8 | 1.18 | 1.15 | 1.48 | 1.14 | 1.25 | 1.12 | 1.27 | 1.17 | 1.09 | 0.43 | 0.12 | 0.36 | S100A14 | Protein S100-A14 OS=Homo sapiens GN=S100A14 PE=1 SV=1 - [S10AE_HUMAN] | 2.65 | 25.00 | 2.00 | 104.00 | 11.65 | 5.24 |
| 102 | A8K4W7 | 1.80 | 1.70 | 2.09 | 1.71 | 1.88 | 1.66 | 1.86 | 1.75 | 1.06 | 0.45 | 0.09 | 0.35 | SUCLG1 | cDNA FLJ76284, highly similar to Homo sapiens succinate-CoA ligase, GDP-forming, alpha subunit (SUCLG1), mRNA OS=Homo sapiens PE=2 | 14.91 | 8.11 | 2.00 | 333.00 | 35.03 | 8.79 |
| 103 | P13645 | 1.40 | 1.39 | 1.71 | 1.70 | 2.25 | 1.30 | 1.50 | 1.75 | 0.86 | 0.45 | -0.22 | 0.35 | KRT10 | Keratin, type I cytoskeletal 10 OS=Homo sapiens GN=KRT10 PE=1 SV=6 - [K1C10_HUMAN] | 7.09 | 6.68 | 2.00 | 584.00 | 58.79 | 5.21 |
| 104 | D6RF44 | 1.57 | 1.66 | 1.77 | 1.66 | 1.65 | 1.49 | 1.67 | 1.60 | 1.04 | 0.45 | 0.06 | 0.35 | HNRNPD | Heterogeneous nuclear ribonucleoprotein D0 (Fragment) OS=Homo sapiens GN=HNRNPD PE=2 SV=1 - [D6RF44_HUMAN] | 21.84 | 19.64 | 2.00 | 112.00 | 12.63 | 8.57 |
| 105 | F8VVM2 | 1.67 | 1.53 | 1.77 | 1.57 | 1.70 | 1.46 | 1.66 | 1.58 | 1.05 | 0.45 | 0.07 | 0.34 | SLC25A3 | Phosphate carrier protein, mitochondrial OS=Homo sapiens GN=SLC25A3 PE=2 SV=1 - [F8VVM2_HUMAN] | 12.89 | 8.33 | 3.00 | 324.00 | 36.14 | 9.26 |
| 106 | P26232 | 1.37 | 1.41 | 1.74 | 1.17 | 1.70 | 1.14 | 1.51 | 1.33 | 1.13 | 0.47 | 0.18 | 0.33 | CTNNA2 | Catenin alpha-2 OS=Homo sapiens GN=CTNNA2 PE=1 SV=5 - [CTNA2_HUMAN] | 5.32 | 3.67 | 1.00 | 953.00 | 105.25 | 5.71 |
| 107 | H3BRM5 | 1.40 | 1.44 | 1.45 | 1.40 | 1.58 | 1.45 | 1.43 | 1.47 | 0.97 | 0.48 | -0.04 | 0.32 | COX5A | Cytochrome c oxidase subunit 5A, mitochondrial OS=Homo sapiens GN=COX5A PE=2 SV=1 - [H3BRM5_HUMAN] | 4.82 | 18.84 | 2.00 | 69.00 | 7.77 | 6.02 |
| 108 | Q562M3 | 1.61 | 1.58 | 1.97 | 1.54 | 1.80 | 1.43 | 1.72 | 1.59 | 1.08 | 0.48 | 0.11 | 0.32 | ACT | Actin-like protein (Fragment) OS=Homo sapiens GN=ACT PE=2 SV=1 - [Q562M3_HUMAN] | 13.84 | 17.48 | 1.00 | 103.00 | 11.53 | 6.68 |
| 109 | P30084 | 1.24 | 1.13 | 1.29 | 1.11 | 1.31 | 0.99 | 1.22 | 1.14 | 1.07 | 0.48 | 0.10 | 0.32 | ECHS1 | Enoyl-CoA hydratase, mitochondrial OS=Homo sapiens GN=ECHS1 PE=1 SV=4 - [ECHM_HUMAN] | 4.12 | 4.48 | 1.00 | 290.00 | 31.37 | 8.07 |
| 110 | H0YE40 | 1.44 | 1.56 | 1.96 | 1.52 | 1.71 | 1.28 | 1.65 | 1.50 | 1.10 | 0.49 | 0.14 | 0.31 | CD44 | CD44 antigen (Fragment) OS=Homo sapiens GN=CD44 PE=2 SV=1 - [H0YE40_HUMAN] | 7.36 | 35.37 | 2.00 | 82.00 | 9.10 | 8.16 |
| 111 | D6R9B6 | 1.12 | 0.79 | 1.03 | 0.94 | 0.86 | 0.90 | 0.98 | 0.90 | 1.09 | 0.49 | 0.12 | 0.31 | RPS3A | 40S ribosomal protein S3a OS=Homo sapiens GN=RPS3A PE=4 SV=1 - [D6R9B6_HUMAN] | 7.16 | 8.28 | 1.00 | 145.00 | 16.53 | 9.25 |
| 112 | B4DMD1 | 1.46 | 1.44 | 1.64 | 1.62 | 1.85 | 1.40 | 1.51 | 1.62 | 0.93 | 0.49 | -0.10 | 0.31 | | cDNA FLJ53360, highly similar to Heterogeneous nuclear ribonucleoprotein R OS=Homo sapiens PE=2 SV=1 - [B4DMD1_HUMAN] | 10.87 | 6.98 | 3.00 | 473.00 | 52.95 | 9.32 |
| 113 | B4DM82 | 2.03 | 2.31 | 3.03 | 2.15 | 2.46 | 2.03 | 2.46 | 2.21 | 1.11 | 0.50 | 0.15 | 0.30 | | cDNA FLJ53060, moderately similar to Peptidyl-prolyl cis-trans isomerase A (EC 5.2.1.8) OS=Homo sapiens PE=2 SV=1 - [B4DM82_HUMAN] | 9.02 | 10.08 | 1.00 | 129.00 | 14.08 | 8.47 |
| 114 | B4DNX1 | 1.90 | 2.33 | 2.70 | 2.09 | 2.41 | 1.77 | 2.31 | 2.09 | 1.11 | 0.50 | 0.14 | 0.30 | | cDNA FLJ53752, highly similar to Heat shock 70 kDa protein 1 OS=Homo sapiens PE=2 SV=1 - [B4DNX1_HUMAN] | 24.41 | 11.03 | 1.00 | 417.00 | 45.10 | 5.73 |
| 115 | J3KRY3 | 1.88 | 1.77 | 2.20 | 1.86 | 1.95 | 1.73 | 1.95 | 1.85 | 1.06 | 0.50 | 0.08 | 0.30 | SNRPN | Small nuclear ribonucleoprotein-associated protein N (Fragment) OS=Homo sapiens GN=SNRPN PE=4 SV=1 - [J3KRY3_HUMAN] | 2.31 | 11.86 | 1.00 | 59.00 | 6.96 | 9.52 |
| 116 | E9PP36 | 1.69 | 1.49 | 1.66 | 1.54 | 1.69 | 1.38 | 1.61 | 1.53 | 1.05 | 0.51 | 0.07 | 0.29 | RPL8 | 60S ribosomal protein L8 OS=Homo sapiens GN=RPL8 PE=2 SV=1 - [E9PP36_HUMAN] | 2.96 | 7.43 | 1.00 | 148.00 | 16.17 | 11.90 |
| 117 | Q96HX3 | 1.43 | 1.32 | 1.44 | 1.28 | 1.50 | 1.22 | 1.40 | 1.33 | 1.05 | 0.52 | 0.07 | 0.29 | | Similar to ribophorin I (Fragment) OS=Homo sapiens PE=2 SV=1 - [Q96HX3_HUMAN] | 6.17 | 6.69 | 3.00 | 568.00 | 64.54 | 6.55 |
| 118 | B4DE36 | 1.58 | 1.20 | 1.25 | 1.44 | 1.62 | 1.30 | 1.35 | 1.45 | 0.93 | 0.52 | -0.11 | 0.29 | | Glucose-6-phosphate isomerase OS=Homo sapiens PE=2 SV=1 - [B4DE36_HUMAN] | 3.02 | 4.91 | 2.00 | 530.00 | 60.15 | 8.15 |
| 119 | Q6PKI6 | 1.66 | 1.80 | 1.97 | 1.71 | 1.94 | 1.40 | 1.81 | 1.68 | 1.07 | 0.52 | 0.10 | 0.28 | YBX1 | YBX1 protein (Fragment) OS=Homo sapiens GN=YBX1 PE=2 SV=1 - [Q6PKI6_HUMAN] | 3.23 | 7.14 | 1.00 | 266.00 | 29.36 | 10.23 |
| 120 | B4E0E1 | 1.84 | 1.67 | 1.84 | 1.71 | 1.86 | 1.57 | 1.78 | 1.71 | 1.04 | 0.52 | 0.06 | 0.28 | | cDNA FLJ53442, highly similar to Poly (ADP-ribose) polymerase 1 (EC 2.4.2.30) OS=Homo sapiens PE=2 SV=1 - [B4E0E1_HUMAN] | 2.30 | 1.31 | 1.00 | 993.00 | 111.06 | 8.87 |
| 121 | Q4ZG51 | 1.60 | 1.64 | 1.95 | 1.74 | 1.95 | 1.77 | 1.73 | 1.82 | 0.95 | 0.54 | -0.07 | 0.27 | FNBP3 | Putative uncharacterized protein FNBP3 OS=Homo sapiens GN=FNBP3 PE=2 SV=1 - [Q4ZG51_HUMAN] | 2.00 | 4.88 | 1.00 | 205.00 | 24.28 | 9.72 |
| 122 | H3BRG4 | 1.92 | 1.72 | 1.96 | 1.85 | 2.12 | 1.86 | 1.86 | 1.94 | 0.96 | 0.54 | -0.06 | 0.27 | UQCRC2 | Cytochrome b-c1 complex subunit 2, mitochondrial OS=Homo sapiens GN=UQCRC2 PE=2 SV=1 - [H3BRG4_HUMAN] | 3.82 | 3.88 | 1.00 | 412.00 | 44.61 | 9.00 |
| 123 | Q0QF37 | 1.53 | 1.47 | 1.64 | 1.36 | 1.65 | 1.43 | 1.55 | 1.48 | 1.04 | 0.55 | 0.06 | 0.26 | MDH2 | Malate dehydrogenase (Fragment) OS=Homo sapiens GN=MDH2 PE=2 SV=1 - [Q0QF37_HUMAN] | 73.67 | 34.43 | 9.00 | 305.00 | 31.95 | 7.88 |
| 124 | Q96B49 | 1.21 | 1.22 | 1.57 | 1.34 | 1.59 | 1.35 | 1.33 | 1.43 | 0.93 | 0.55 | -0.10 | 0.26 | TOMM6 | Mitochondrial import receptor subunit TOM6 homolog OS=Homo sapiens GN=TOMM6 PE=1 SV=1 - [TOM6_HUMAN] | 2.42 | 18.92 | 1.00 | 74.00 | 8.00 | 4.89 |
| 125 | H7C4L9 | 1.51 | 1.42 | 1.51 | 1.39 | 1.60 | 1.22 | 1.48 | 1.41 | 1.05 | 0.55 | 0.07 | 0.26 | ATP1B3 | Sodium/potassium-transporting ATPase subunit beta-3 (Fragment) OS=Homo sapiens GN=ATP1B3 PE=2 SV=1 - [H7C4L9_HUMAN] | 2.30 | 40.74 | 1.00 | 27.00 | 2.79 | 8.22 |
| 126 | B3KTN4 | 1.70 | 1.65 | 1.86 | 1.70 | 1.78 | 1.54 | 1.73 | 1.67 | 1.04 | 0.56 | 0.05 | 0.25 | | Citrate synthase OS=Homo sapiens PE=2 SV=1 - [B3KTN4_HUMAN] | 18.37 | 9.50 | 4.00 | 421.00 | 47.05 | 7.24 |
| 127 | Q96Q74 | 1.28 | 1.25 | 1.44 | 1.26 | 1.48 | 1.39 | 1.32 | 1.38 | 0.96 | 0.56 | -0.06 | 0.25 | MRPL7/L12 | Mitochondrial ribosomal protein L7/L12 (Fragment) OS=Homo sapiens GN=MRPL7/L12 PE=2 SV=1 - [Q96Q74_HUMAN] | 6.99 | 42.37 | 2.00 | 59.00 | 6.38 | 9.04 |
| 128 | B7Z254 | 1.62 | 1.57 | 1.87 | 1.62 | 1.76 | 1.45 | 1.69 | 1.61 | 1.05 | 0.57 | 0.07 | 0.24 | PDIA6 | Protein disulfide-isomerase A6 OS=Homo sapiens GN=PDIA6 PE=2 SV=1 - [B7Z254_HUMAN] | 18.06 | 9.84 | 3.00 | 437.00 | 47.81 | 5.08 |
| 129 | H7BY36 | 1.21 | 1.24 | 1.41 | 1.39 | 1.59 | 1.14 | 1.29 | 1.37 | 0.94 | 0.58 | -0.09 | 0.24 | EWSR1 | RNA-binding protein EWS (Fragment) OS=Homo sapiens GN=EWSR1 PE=4 SV=1 - [H7BY36_HUMAN] | 6.15 | 4.55 | 1.00 | 308.00 | 32.16 | 9.82 |
| 130 | B4E0X6 | 1.50 | 1.38 | 1.75 | 1.44 | 1.57 | 1.42 | 1.55 | 1.48 | 1.05 | 0.58 | 0.07 | 0.24 | | Proteasome subunit alpha type OS=Homo sapiens PE=2 SV=1 - [B4E0X6_HUMAN] | 2.52 | 8.46 | 1.00 | 130.00 | 14.61 | 8.51 |
| 131 | C9JK10 | 1.32 | 1.44 | 1.57 | 1.46 | 1.71 | 1.38 | 1.44 | 1.52 | 0.95 | 0.58 | -0.07 | 0.24 | | Integrin alpha-6 heavy chain (Fragment) OS=Homo sapiens GN=ITGA6 PE=2 SV=1 - [C9JK10_HUMAN] | 2.64 | 0.74 | 1.00 | 1086.00 | 121.59 | 7.36 |
| 132 | F8VPV9 | 1.52 | 1.41 | 1.56 | 1.52 | 1.75 | 1.41 | 1.50 | 1.56 | 0.96 | 0.60 | -0.06 | 0.22 | | ATP synthase subunit beta OS=Homo sapiens GN=ATP5B PE=2 SV=1 - [F8VPV9_HUMAN] | 39.23 | 27.03 | 10.00 | 518.00 | 55.27 | 5.40 |
| 133 | F8VUA6 | 1.56 | 1.38 | 1.48 | 1.41 | 1.59 | 1.23 | 1.48 | 1.41 | 1.05 | 0.60 | 0.07 | 0.22 | RPL18 | 60S ribosomal protein L18 (Fragment) OS=Homo sapiens GN=RPL18 PE=2 SV=1 - [F8VUA6_HUMAN] | 10.41 | 7.69 | 1.00 | 130.00 | 14.52 | 11.75 |
| 134 | C9K0U8 | 1.65 | 1.67 | 1.88 | 1.71 | 1.81 | 1.47 | 1.73 | 1.66 | 1.04 | 0.60 | 0.06 | 0.22 | SSBP1 | Single-stranded DNA-binding protein (Fragment) OS=Homo sapiens GN=SSBP1 PE=3 SV=1 - [C9K0U8_HUMAN] | 3.35 | 24.79 | 3.00 | 121.00 | 14.12 | 9.57 |
| 135 | Q15149 | 1.54 | 1.40 | 1.75 | 1.49 | 1.63 | 1.36 | 1.56 | 1.49 | 1.05 | 0.60 | 0.07 | 0.22 | PLEC | Plectin OS=Homo sapiens GN=PLEC PE=1 SV=3 - [PLEC_HUMAN] | 18.64 | 2.07 | 4.00 | 4684.00 | 531.47 | 5.96 |
| 136 | A8K9J7 | 1.52 | 1.40 | 1.74 | 1.35 | 1.73 | 1.31 | 1.55 | 1.46 | 1.06 | 0.60 | 0.09 | 0.22 | | Histone H2B OS=Homo sapiens PE=2 SV=1 - [A8K9J7_HUMAN] | 11.28 | 12.70 | 2.00 | 126.00 | 13.99 | 10.32 |
| 137 | B4DY09 | 1.33 | 1.23 | 1.43 | 1.31 | 1.47 | 1.34 | 1.33 | 1.37 | 0.97 | 0.61 | -0.04 | 0.22 | ILF2 | cDNA FLJ51660, highly similar to Interleukin enhancer-binding factor 2 OS=Homo sapiens PE=2 SV=1 - [B4DY09_HUMAN] | 6.51 | 3.13 | 1.00 | 352.00 | 38.89 | 4.94 |
| 138 | Q86TW7 | 1.59 | 1.66 | 1.95 | 1.73 | 1.87 | 1.79 | 1.73 | 1.80 | 0.96 | 0.61 | -0.05 | 0.22 | DLST | Dihydrolipoamide S-succinyltransferase (E2 component of 2-oxo-glutarate complex), isoform CRA_b (Fragment) OS=Homo sapiens GN=DLST PE=2 SV=1 - [Q86TW7_HUMAN] | 2.28 | 3.19 | 1.00 | 251.00 | 26.87 | 10.08 |
| 139 | B4E364 | 1.79 | 1.70 | 1.91 | 1.60 | 1.94 | 1.66 | 1.80 | 1.73 | 1.04 | 0.61 | 0.05 | 0.21 | | cDNA FLJ54954, highly similar to Ras-related protein Ral-B OS=Homo sapiens PE=2 SV=1 - [B4E364_HUMAN] | 2.53 | 6.93 | 1.00 | 101.00 | 11.83 | 9.51 |
| 140 | P25705 | 1.64 | 1.57 | 1.83 | 1.55 | 1.80 | 1.48 | 1.68 | 1.61 | 1.04 | 0.61 | 0.06 | 0.21 | ATP5F1A | ATP synthase subunit alpha, mitochondrial OS=Homo sapiens GN=ATP5A1 PE=1 SV=1 - [ATPA_HUMAN] | 50.90 | 24.59 | 12.00 | 553.00 | 59.71 | 9.13 |
| 141 | Q96RX5 | 1.13 | 1.60 | 1.65 | 1.40 | 1.47 | 1.23 | 1.46 | 1.36 | 1.07 | 0.61 | 0.10 | 0.21 | NDUFB10 | NADH ubiquinone oxidoreductase PDSW subunit (RH 16p13.3) (Fragment) OS=Homo sapiens GN=NDUFB10 PE=2 SV=1 - [Q96RX5_HUMAN] | 2.12 | 32.56 | 1.00 | 43.00 | 5.08 | 6.60 |
| 142 | Q9BVT0 | 1.84 | 1.84 | 2.44 | 1.78 | 2.16 | 1.80 | 2.04 | 1.91 | 1.07 | 0.62 | 0.09 | 0.21 | ARHA | ARHA protein (Fragment) OS=Homo sapiens GN=ARHA PE=2 SV=2 - [Q9BVT0_HUMAN] | 3.12 | 7.92 | 1.00 | 101.00 | 11.52 | 9.35 |
| 143 | P0CG22 | 1.60 | 1.45 | 1.54 | 1.48 | 1.78 | 1.50 | 1.53 | 1.59 | 0.96 | 0.62 | -0.05 | 0.21 | DHRS4L1 | Putative dehydrogenase/reductase SDR family member 4-like 2 OS=Homo sapiens GN=DHRS4L1 PE=5 SV=1 - [DR4L1_HUMAN] | 2.57 | 6.41 | 1.00 | 281.00 | 30.59 | 9.77 |
| 144 | A8K787 | 1.41 | 1.52 | 1.71 | 1.55 | 1.75 | 1.51 | 1.54 | 1.60 | 0.96 | 0.62 | -0.06 | 0.21 | | cDNA FLJ75273, highly similar to Homo sapiens solute carrier family 25 (mitochondrial carrier; adenine nucleotide translocator), member 4, mRNA OS=Homo sapiens PE=2 SV=1 - [A8K787_HUMAN] | 9.13 | 7.38 | 3.00 | 298.00 | 33.06 | 9.76 |
| 145 | Q7Z4Q5 | 1.29 | 1.26 | 1.36 | 1.27 | 1.42 | 1.31 | 1.30 | 1.33 | 0.98 | 0.62 | -0.03 | 0.21 | HNRPU | Heterogeneous nuclear ribonucleoprotein U (Scaffold attachment factor A), isoform CRA_a OS=Homo sapiens GN=HNRPU PE=2 SV=1 - | 2.06 | 1.36 | 1.00 | 513.00 | 57.63 | 9.36 |
| 146 | Q5JP01 | 1.77 | 1.73 | 2.15 | 1.69 | 2.01 | 1.67 | 1.88 | 1.79 | 1.05 | 0.62 | 0.07 | 0.20 | RBBP7 | Histone-binding protein RBBP7 (Fragment) OS=Homo sapiens GN=RBBP7 PE=2 SV=1 - [Q5JP01_HUMAN] | 2.40 | 4.21 | 1.00 | 285.00 | 31.48 | 6.19 |
| 147 | B2R984 | 1.75 | 1.72 | 2.01 | 1.81 | 1.90 | 1.55 | 1.83 | 1.75 | 1.04 | 0.63 | 0.06 | 0.20 | | cDNA, FLJ94268, highly similar to Homo sapiens histone 1, H1e (HIST1H1E), mRNA OS=Homo sapiens PE=2 SV=1 - [B2R984_HUMAN] | 32.50 | 26.48 | 5.00 | 219.00 | 21.88 | 11.03 |
| 148 | B5MD38 | 1.53 | 1.50 | 1.69 | 1.58 | 1.91 | 1.46 | 1.58 | 1.65 | 0.95 | 0.63 | -0.07 | 0.20 | HADHB | 3-ketoacyl-CoA thiolase OS=Homo sapiens GN=HADHB PE=2 SV=1 - [B5MD38_HUMAN] | 4.67 | 7.41 | 3.00 | 351.00 | 37.90 | 9.45 |
| 149 | C9JW96 | 1.55 | 1.63 | 1.88 | 1.66 | 1.74 | 1.45 | 1.69 | 1.62 | 1.04 | 0.63 | 0.06 | 0.20 | PHB | Prohibitin (Fragment) OS=Homo sapiens GN=PHB PE=2 SV=2 - [C9JW96_HUMAN] | 18.96 | 32.52 | 7.00 | 246.00 | 26.88 | 5.40 |
| 150 | P58107 | 1.02 | 1.00 | 1.26 | 1.07 | 1.14 | 0.90 | 1.09 | 1.04 | 1.06 | 0.63 | 0.08 | 0.20 | EPPK1 | Epiplakin OS=Homo sapiens GN=EPPK1 PE=1 SV=2 - [EPIPL_HUMAN] | 11.84 | 6.35 | 2.00 | 5090.00 | 555.28 | 5.60 |
| 151 | P51148 | 2.09 | 1.82 | 2.03 | 1.98 | 1.99 | 1.83 | 1.98 | 1.93 | 1.03 | 0.64 | 0.04 | 0.20 | RAB5C | Ras-related protein Rab-5C OS=Homo sapiens GN=RAB5C PE=1 SV=2 - [RAB5C_HUMAN] | 2.29 | 6.48 | 1.00 | 216.00 | 23.47 | 8.41 |

| | | | | | | | | | | | | | | | | | | | | | |
|---|---|---|---|---|---|---|---|---|---|---|---|---|---|---|---|---|---|---|---|---|---|
| 152 | Q6PIX2 | 1.79 | 1.63 | 1.94 | 1.54 | 1.99 | 1.58 | 1.79 | 1.70 | 1.05 | 0.64 | 0.07 | 0.20 | SFPQ | SFPQ protein (Fragment) OS=Homo sapiens GN=SFPQ PE=2 SV=1 - [Q6PIX2_HUMAN] | 4.78 | 5.33 | 3.00 | 525.00 | 55.44 | 9.89 |
| 153 | P62424 | 1.61 | 1.68 | 1.83 | 1.64 | 1.80 | 1.51 | 1.71 | 1.65 | 1.03 | 0.64 | 0.05 | 0.20 | RPL7A | 60S ribosomal protein L7a OS=Homo sapiens GN=RPL7A PE=1 SV=2 - [RL7A_HUMAN] | 32.79 | 12.03 | 3.00 | 266.00 | 29.98 | 10.61 |
| 154 | G3V4Y7 | 1.94 | 1.96 | 2.26 | 1.93 | 2.22 | 1.76 | 2.06 | 1.97 | 1.04 | 0.64 | 0.06 | 0.19 | KTN1 | Kinectin OS=Homo sapiens GN=KTN1 PE=2 SV=1 - [G3V4Y7_HUMAN] | 2.65 | 2.02 | 1.00 | 595.00 | 68.90 | 5.22 |
| 155 | H0YEX5 | 1.52 | 1.55 | 1.67 | 1.54 | 1.73 | 1.57 | 1.58 | 1.62 | 0.98 | 0.64 | -0.03 | 0.19 | SF3B2 | Splicing factor 3B subunit 2 (Fragment) OS=Homo sapiens GN=SF3B2 PE=4 SV=1 - [H0YEX5_HUMAN] | 8.84 | 7.62 | 2.00 | 315.00 | 34.99 | 4.81 |
| 156 | Q49AJ9 | 1.42 | 1.38 | 1.66 | 1.51 | 1.60 | 1.49 | 1.49 | 1.53 | 0.97 | 0.64 | -0.04 | 0.19 | RPL3 | RPL3 protein OS=Homo sapiens GN=RPL3 PE=2 SV=1 - [Q49AJ9_HUMAN] | 2.47 | 2.79 | 1.00 | 251.00 | 28.56 | 10.17 |
| 157 | E7EN95 | 1.52 | 1.38 | 1.47 | 1.49 | 1.61 | 1.39 | 1.46 | 1.49 | 0.97 | 0.65 | -0.04 | 0.19 | FLNB | Filamin-B OS=Homo sapiens GN=FLNB PE=2 SV=1 - [E7EN95_HUMAN] | 5.66 | 1.08 | 2.00 | 2409.00 | 256.12 | 5.73 |
| 158 | B3KUZ8 | 1.68 | 1.66 | 1.91 | 1.68 | 1.83 | 1.59 | 1.75 | 1.70 | 1.03 | 0.65 | 0.04 | 0.19 | | Aspartate aminotransferase OS=Homo sapiens PE=2 SV=1 - [B3KUZ8_HUMAN] | 30.59 | 16.98 | 5.00 | 371.00 | 41.30 | 8.84 |
| 159 | Q53G71 | 1.56 | 1.66 | 1.82 | 1.62 | 1.78 | 1.47 | 1.68 | 1.62 | 1.04 | 0.65 | 0.05 | 0.19 | | Calreticulin variant (Fragment) OS=Homo sapiens PE=2 SV=1 - [Q53G71_HUMAN] | 51.75 | 10.59 | 5.00 | 406.00 | 46.89 | 4.45 |
| 160 | Q96AG4 | 1.54 | 1.52 | 1.66 | 1.48 | 1.64 | 1.50 | 1.57 | 1.54 | 1.02 | 0.65 | 0.03 | 0.19 | LRRC59 | Leucine-rich repeat-containing protein 59 OS=Homo sapiens GN=LRRC59 PE=1 SV=1 - [LRC59_HUMAN] | 9.84 | 3.91 | 1.00 | 307.00 | 34.91 | 9.57 |
| 161 | P62304 | 1.42 | 1.36 | 1.49 | 1.40 | 1.82 | 1.30 | 1.42 | 1.50 | 0.95 | 0.65 | -0.08 | 0.19 | SNRPE | Small nuclear ribonucleoprotein E OS=Homo sapiens GN=SNRPE PE=1 SV=1 - [RUXE_HUMAN] | 4.08 | 25.00 | 2.00 | 92.00 | 10.80 | 9.44 |
| 162 | P49411 | 1.53 | 1.57 | 1.81 | 1.56 | 1.70 | 1.50 | 1.64 | 1.59 | 1.03 | 0.65 | 0.04 | 0.18 | TUFM | Elongation factor Tu, mitochondrial OS=Homo sapiens GN=TUFM PE=1 SV=2 - [EFTU_HUMAN] | 25.01 | 12.17 | 5.00 | 452.00 | 49.51 | 7.61 |
| 163 | D6R9X8 | 1.71 | 1.60 | 1.72 | 1.66 | 1.82 | 1.64 | 1.67 | 1.71 | 0.98 | 0.66 | -0.03 | 0.18 | ITGA3 | Integrin alpha-3 OS=Homo sapiens GN=ITGA3 PE=2 SV=2 - [D6R9X8_HUMAN] | 1.91 | 6.34 | 1.00 | 142.00 | 15.22 | 8.21 |
| 164 | K7EMV3 | 1.45 | 1.34 | 1.50 | 1.39 | 1.49 | 1.30 | 1.43 | 1.40 | 1.02 | 0.66 | 0.04 | 0.18 | H3-3B | Histone H3 OS=Homo sapiens GN=H3F3B PE=3 SV=1 - [K7EMV3_HUMAN] | 15.00 | 17.39 | 2.00 | 92.00 | 10.33 | 11.82 |
| 165 | P84090 | 1.36 | 1.38 | 1.51 | 1.45 | 1.60 | 1.32 | 1.41 | 1.46 | 0.97 | 0.66 | -0.04 | 0.18 | ERH | Enhancer of rudimentary homolog OS=Homo sapiens GN=ERH PE=1 SV=1 - [ERH_HUMAN] | 9.62 | 5.77 | 1.00 | 104.00 | 12.25 | 5.92 |
| 166 | H7C469 | 1.35 | 1.23 | 1.47 | 1.35 | 1.37 | 1.22 | 1.35 | 1.31 | 1.03 | 0.67 | 0.04 | 0.17 | | Uncharacterized protein (Fragment) OS=Homo sapiens PE=3 SV=1 - [H7C469_HUMAN] | 9.35 | 7.12 | 2.00 | 379.00 | 40.42 | 5.76 |
| 167 | Q13751 | 1.00 | 1.02 | 1.08 | 0.98 | 1.14 | 0.89 | 1.04 | 1.00 | 1.03 | 0.68 | 0.05 | 0.17 | LAMB3 | Laminin subunit beta-3 OS=Homo sapiens GN=LAMB3 PE=1 SV=1 - [LAMB3_HUMAN] | 5.65 | 2.90 | 2.00 | 1172.00 | 129.49 | 7.21 |
| 168 | B4DFG4 | 1.72 | 1.58 | 1.85 | 1.69 | 1.84 | 1.44 | 1.72 | 1.66 | 1.04 | 0.68 | 0.05 | 0.17 | | cDNA FLJ58801, highly similar to Interleukin enhancer-binding factor 3 OS=Homo sapiens PE=2 SV=1 - [B4DFG4_HUMAN] | 27.45 | 5.14 | 3.00 | 506.00 | 54.67 | 8.81 |
| 169 | P14314 | 1.37 | 1.36 | 1.73 | 1.35 | 1.56 | 1.36 | 1.48 | 1.42 | 1.04 | 0.69 | 0.06 | 0.16 | PRKCSH | Glucosidase 2 subunit beta OS=Homo sapiens GN=PRKCSH PE=1 SV=2 - [GLU2B_HUMAN] | 6.22 | 5.68 | 3.00 | 528.00 | 59.39 | 4.41 |
| 170 | M0QY67 | 1.70 | 1.55 | 1.87 | 1.59 | 1.80 | 1.58 | 1.71 | 1.66 | 1.03 | 0.69 | 0.04 | 0.16 | ETFB | Electron transfer flavoprotein subunit beta (Fragment) OS=Homo sapiens GN=ETFB PE=4 SV=1 - [M0QY67_HUMAN] | 2.67 | 6.74 | 1.00 | 178.00 | 19.51 | 7.74 |
| 171 | H0YCY8 | 1.58 | 1.50 | 1.63 | 1.53 | 1.77 | 1.53 | 1.57 | 1.61 | 0.98 | 0.69 | -0.03 | 0.16 | CTSC | Dipeptidyl peptidase 1 exclusion domain chain (Fragment) OS=Homo sapiens GN=CTSC PE=4 SV=1 - [H0YCY8_HUMAN] | 2.51 | 4.90 | 1.00 | 245.00 | 27.92 | 9.16 |
| 172 | D6REM6 | 1.55 | 1.54 | 1.62 | 1.51 | 1.66 | 1.47 | 1.57 | 1.54 | 1.02 | 0.69 | 0.03 | 0.16 | MATR3 | Matrin-3 OS=Homo sapiens GN=MATR3 PE=2 SV=1 - [D6REM6_HUMAN] | 11.45 | 3.78 | 3.00 | 794.00 | 88.30 | 5.76 |
| 173 | B7Z4V2 | 1.57 | 1.65 | 1.88 | 1.64 | 1.78 | 1.54 | 1.70 | 1.65 | 1.03 | 0.70 | 0.04 | 0.16 | | cDNA FLJ51907, highly similar to Stress-70 protein, mitochondrial OS=Homo sapiens PE=2 SV=1 - [B7Z4V2_HUMAN] | 49.97 | 14.59 | 8.00 | 665.00 | 72.36 | 5.94 |
| 174 | P62805 | 1.66 | 1.48 | 1.87 | 1.53 | 1.80 | 1.50 | 1.67 | 1.61 | 1.04 | 0.70 | 0.05 | 0.16 | H4C1 | Histone H4 OS=Homo sapiens GN=HIST1H4A PE=1 SV=2 - [H4_HUMAN] | 7.27 | 19.42 | 2.00 | 103.00 | 11.36 | 11.36 |
| 175 | P62263 | 1.58 | 1.59 | 1.87 | 1.58 | 1.84 | 1.43 | 1.68 | 1.62 | 1.04 | 0.70 | 0.06 | 0.15 | RPS14 | 40S ribosomal protein S14 OS=Homo sapiens GN=RPS14 PE=1 SV=3 - [RS14_HUMAN] | 2.49 | 8.61 | 1.00 | 151.00 | 16.26 | 10.05 |
| 176 | Q6DC98 | 1.49 | 1.43 | 1.73 | 1.45 | 1.71 | 1.31 | 1.55 | 1.49 | 1.04 | 0.71 | 0.06 | 0.15 | LMNB1 | LMNB1 protein (Fragment) OS=Homo sapiens GN=LMNB1 PE=2 SV=1 - [Q6DC98_HUMAN] | 3.48 | 5.71 | 2.00 | 333.00 | 38.12 | 5.45 |
| 177 | B4E1S3 | 1.25 | 1.31 | 1.41 | 1.32 | 1.44 | 1.30 | 1.33 | 1.35 | 0.98 | 0.71 | -0.03 | 0.15 | | cDNA FLJ57860, highly similar to Transmembrane protein 109 OS=Homo sapiens PE=2 SV=1 - [B4E1S3_HUMAN] | 3.20 | 5.13 | 1.00 | 234.00 | 24.97 | 9.54 |
| 178 | P13667 | 1.70 | 1.59 | 1.78 | 1.63 | 1.84 | 1.44 | 1.69 | 1.64 | 1.03 | 0.71 | 0.04 | 0.15 | PDIA4 | Protein disulfide-isomerase A4 OS=Homo sapiens GN=PDIA4 PE=1 SV=2 - [PDIA4_HUMAN] | 8.66 | 5.27 | 4.00 | 645.00 | 72.89 | 5.07 |
| 179 | P46783 | 2.24 | 1.92 | 2.30 | 1.95 | 2.49 | 2.24 | 2.15 | 2.23 | 0.97 | 0.71 | -0.05 | 0.15 | RPS10 | 40S ribosomal protein S10 OS=Homo sapiens GN=RPS10 PE=1 SV=1 - [RS10_HUMAN] | 5.44 | 5.45 | 1.00 | 165.00 | 18.89 | 10.15 |
| 180 | A8KA83 | 1.50 | 1.35 | 1.69 | 1.50 | 1.55 | 1.36 | 1.51 | 1.47 | 1.03 | 0.71 | 0.04 | 0.15 | | cDNA FLJ78586, highly similar to Homo sapiens VAMP (vesicle-associated membrane protein)-associated protein A, 33kDa (VAPA), mRNA OS=Homo sapiens PE=2 SV=1 - [A8KA83_HUMAN] | 8.54 | 14.05 | 2.00 | 242.00 | 27.30 | 8.62 |
| 181 | B4DEI3 | 1.82 | 1.80 | 2.20 | 1.86 | 2.09 | 1.66 | 1.94 | 1.87 | 1.04 | 0.71 | 0.05 | 0.15 | | cDNA FLJ57715, highly similar to Voltage-dependent anion-selective channel protein 1 OS=Homo sapiens PE=2 SV=1 - [B4DEI3_HUMAN] | 28.21 | 17.95 | 2.00 | 156.00 | 16.68 | 9.39 |
| 182 | B3KQT9 | 1.62 | 1.61 | 1.82 | 1.57 | 1.81 | 1.53 | 1.68 | 1.64 | 1.03 | 0.72 | 0.04 | 0.14 | | cDNA PSEC0175 fis, clone OVARC1000169, highly similar to Protein disulfide-isomerase A3 (EC 5.3.4.1) OS=Homo sapiens PE=2 SV=1 - | 48.28 | 22.29 | 8.00 | 480.00 | 54.07 | 7.21 |
| 183 | Q9UNM1 | 1.53 | 1.43 | 1.55 | 1.45 | 1.63 | 1.32 | 1.50 | 1.47 | 1.02 | 0.73 | 0.03 | 0.14 | EPFP1 | Chaperonin 10-related protein (Fragment) OS=Homo sapiens GN=EPFP1 PE=2 SV=1 - [Q9UNM1_HUMAN] | 26.40 | 55.67 | 5.00 | 97.00 | 10.29 | 9.00 |
| 184 | B2R5H0 | 1.70 | 1.68 | 1.81 | 1.64 | 1.90 | 1.52 | 1.73 | 1.69 | 1.03 | 0.73 | 0.04 | 0.14 | | cDNA, FLJ92471, highly similar to Homo sapiens S100 calcium binding protein A11 (calgizzarin) (S100A11), mRNA OS=Homo sapiens PE=2 SV=1 - | 1.70 | 8.57 | 1.00 | 105.00 | 11.72 | 7.18 |
| 185 | P11021 | 1.52 | 1.39 | 1.73 | 1.42 | 1.74 | 1.31 | 1.55 | 1.49 | 1.04 | 0.74 | 0.05 | 0.13 | HSPA5 | 78 kDa glucose-regulated protein OS=Homo sapiens GN=HSPA5 PE=1 SV=2 - [GRP78_HUMAN] | 98.37 | 32.57 | 15.00 | 654.00 | 72.29 | 5.16 |
| 186 | A2T926 | 1.46 | 1.43 | 1.49 | 1.38 | 1.62 | 1.29 | 1.46 | 1.43 | 1.02 | 0.75 | 0.03 | 0.12 | TMPO | Thymopentin OS=Homo sapiens GN=TMPO PE=2 SV=1 - [A2T926_HUMAN] | 7.93 | 16.53 | 3.00 | 248.00 | 27.37 | 8.48 |
| 187 | Q59H46 | 1.60 | 1.54 | 1.77 | 1.62 | 1.85 | 1.56 | 1.64 | 1.68 | 0.98 | 0.76 | -0.03 | 0.12 | | Integrin beta (Fragment) OS=Homo sapiens PE=2 SV=1 - [Q59H46_HUMAN] | 3.18 | 1.32 | 2.00 | 1515.00 | 169.04 | 6.49 |
| 188 | P49419 | 1.65 | 1.62 | 1.87 | 1.73 | 1.76 | 1.55 | 1.71 | 1.68 | 1.02 | 0.76 | 0.03 | 0.12 | ALDH7A1 | Alpha-aminoadipic semialdehyde dehydrogenase OS=Homo sapiens GN=ALDH7A1 PE=1 SV=5 - [AL7A1_HUMAN] | 3.11 | 3.15 | 1.00 | 539.00 | 58.45 | 7.99 |
| 189 | F8W8J4 | 1.66 | 1.57 | 1.77 | 1.57 | 1.85 | 1.46 | 1.67 | 1.63 | 1.03 | 0.76 | 0.04 | 0.12 | | Myoferlin OS=Homo sapiens GN=MYOF PE=2 SV=1 - [F8W8J4_HUMAN] | 5.91 | 1.12 | 2.00 | 2061.00 | 234.50 | 6.18 |
| 190 | B5BU24 | 1.22 | 1.26 | 1.61 | 1.36 | 1.63 | 1.26 | 1.36 | 1.42 | 0.96 | 0.76 | -0.06 | 0.12 | YWHAB | 14-3-3 protein beta/alpha OS=Homo sapiens GN=YWHAB PE=2 SV=1 - [B5BU24_HUMAN] | 21.16 | 19.11 | 1.00 | 246.00 | 28.09 | 4.83 |
| 191 | P05109 | 1.07 | 1.03 | 1.24 | 1.02 | 1.21 | 1.02 | 1.11 | 1.09 | 1.03 | 0.77 | 0.04 | 0.11 | S100A8 | Protein S100-A8 OS=Homo sapiens GN=S100A8 PE=1 SV=1 - [S10A8_HUMAN] | 2.31 | 8.60 | 1.00 | 93.00 | 10.83 | 7.03 |
| 192 | E7BSV0 | 1.92 | 1.89 | 2.28 | 1.92 | 2.21 | 1.81 | 2.03 | 1.98 | 1.03 | 0.77 | 0.04 | 0.11 | EGFR | Epidermal growth factor receptor variant A OS=Homo sapiens GN=EGFR PE=2 SV=1 - [E7BSV0_HUMAN] | 6.42 | 1.67 | 2.00 | 1136.00 | 125.72 | 6.65 |
| 193 | E5RK64 | 1.71 | 1.76 | 1.83 | 1.64 | 1.90 | 1.67 | 1.77 | 1.74 | 1.02 | 0.77 | 0.02 | 0.11 | VAPB | Vesicle-associated membrane protein-associated protein B/C OS=Homo sapiens GN=VAPB PE=2 SV=1 - [E5RK64_HUMAN] | 4.97 | 19.72 | 1.00 | 71.00 | 7.80 | 9.42 |
| 194 | P02545 | 1.21 | 1.18 | 1.39 | 1.20 | 1.34 | 1.15 | 1.26 | 1.23 | 1.02 | 0.77 | 0.03 | 0.11 | LMNA | Prelamin-A/C OS=Homo sapiens GN=LMNA PE=1 SV=1 - [LMNA_HUMAN] | 171.62 | 40.36 | 23.00 | 664.00 | 74.09 | 7.02 |
| 195 | F5H823 | 1.84 | 1.69 | 2.09 | 1.74 | 2.08 | 1.64 | 1.87 | 1.82 | 1.03 | 0.79 | 0.04 | 0.10 | RAP1B | Ras-related protein Rap-1b (Fragment) OS=Homo sapiens GN=RAP1B PE=2 SV=1 - [F5H823_HUMAN] | 3.04 | 11.65 | 1.00 | 103.00 | 11.91 | 5.55 |
| 196 | Q01650 | 1.21 | 1.37 | 1.71 | 1.13 | 1.71 | 1.26 | 1.43 | 1.37 | 1.05 | 0.79 | 0.07 | 0.10 | SLC7A5 | Large neutral amino acids transporter small subunit 1 OS=Homo sapiens GN=SLC7A5 PE=1 SV=2 - [LAT1_HUMAN] | 5.88 | 5.33 | 2.00 | 507.00 | 54.97 | 7.72 |
| 197 | M0QYT0 | 1.52 | 1.47 | 1.63 | 1.48 | 1.77 | 1.45 | 1.54 | 1.57 | 0.98 | 0.80 | -0.03 | 0.10 | | Uncharacterized protein (Fragment) OS=Homo sapiens PE=4 SV=1 - [M0QYT0_HUMAN] | 11.22 | 4.36 | 1.00 | 321.00 | 35.99 | 7.42 |
| 198 | Q8N183 | 1.81 | 1.80 | 2.03 | 1.76 | 2.32 | 1.73 | 1.88 | 1.93 | 0.97 | 0.80 | -0.04 | 0.10 | NDUFAF2 | Mimitin, mitochondrial OS=Homo sapiens GN=NDUFAF2 PE=1 SV=1 - [MIMIT_HUMAN] | 2.08 | 5.92 | 1.00 | 169.00 | 19.84 | 8.97 |
| 199 | Q53R94 | 1.61 | 1.49 | 1.33 | 1.37 | 1.58 | 1.40 | 1.48 | 1.45 | 1.02 | 0.80 | 0.03 | 0.10 | RTN4 | Putative uncharacterized protein RTN4 (Fragment) OS=Homo sapiens GN=RTN4 PE=2 SV=1 - [Q53R94_HUMAN] | 3.54 | 7.03 | 1.00 | 185.00 | 19.29 | 4.13 |
| 200 | B3KTP9 | 1.78 | 2.14 | 2.27 | 1.88 | 2.21 | 1.96 | 2.06 | 2.02 | 1.02 | 0.80 | 0.03 | 0.09 | | cDNA FLJ38578 fis, clone HCHON2007674, highly similar to NUCLEOLIN OS=Homo sapiens PE=2 SV=1 - [B3KTP9_HUMAN] | 13.15 | 5.37 | 5.00 | 633.00 | 68.40 | 4.65 |
| 201 | P51572 | 1.41 | 1.51 | 1.83 | 1.61 | 1.68 | 1.56 | 1.58 | 1.62 | 0.98 | 0.81 | -0.03 | 0.09 | BCAP31 | B-cell receptor-associated protein 31 OS=Homo sapiens GN=BCAP31 PE=1 SV=3 - [BAP31_HUMAN] | 3.08 | 3.66 | 1.00 | 246.00 | 27.97 | 8.44 |
| 202 | H7BZJ3 | 1.43 | 1.44 | 1.72 | 1.58 | 1.71 | 1.40 | 1.53 | 1.56 | 0.98 | 0.81 | -0.03 | 0.09 | PDIA3 | Thioredoxin (Fragment) OS=Homo sapiens GN=PDIA3 PE=2 SV=1 - [H7BZJ3_HUMAN] | 12.35 | 34.15 | 1.00 | 123.00 | 13.51 | 7.30 |
| 203 | Q15388 | 1.35 | 0.84 | 1.02 | 0.93 | 1.12 | 1.03 | 1.07 | 1.03 | 1.04 | 0.81 | 0.06 | 0.09 | TOMM20 | Mitochondrial import receptor subunit TOM20 homolog OS=Homo sapiens GN=TOMM20 PE=1 SV=1 - [TOM20_HUMAN] | 1.72 | 8.97 | 1.00 | 145.00 | 16.29 | 8.60 |
| 204 | Q6IPH7 | 2.06 | 2.11 | 2.10 | 1.82 | 2.40 | 1.91 | 2.09 | 2.04 | 1.02 | 0.81 | 0.03 | 0.09 | RPL14 | RPL14 protein OS=Homo sapiens GN=RPL14 PE=2 SV=1 - [Q6IPH7_HUMAN] | 4.20 | 13.64 | 2.00 | 220.00 | 23.77 | 10.93 |
| 205 | B4DSR0 | 1.57 | 1.56 | 1.72 | 1.53 | 1.78 | 1.61 | 1.62 | 1.64 | 0.99 | 0.81 | -0.02 | 0.09 | | cDNA FLJ60080, highly similar to 130 kDa leucine-rich protein (LRP 130) (Fragment) OS=Homo sapiens PE=2 SV=1 - [B4DSR0_HUMAN] | 9.51 | 1.38 | 2.00 | 1087.00 | 123.17 | 5.54 |
| 206 | I3L192 | 1.66 | 1.73 | 2.02 | 1.73 | 2.01 | 1.53 | 1.80 | 1.76 | 1.02 | 0.82 | 0.03 | 0.08 | BSG | Basigin (Fragment) OS=Homo sapiens GN=BSG PE=2 SV=1 - [I3L192_HUMAN] | 2.23 | 5.59 | 1.00 | 161.00 | 17.35 | 5.05 |
| 207 | J3QL15 | 1.85 | 1.76 | 1.96 | 1.82 | 2.12 | 1.71 | 1.85 | 1.89 | 0.98 | 0.82 | -0.02 | 0.08 | | Ribosomal protein L19 (Fragment) OS=Homo sapiens GN=RPL19 PE=3 SV=1 - [J3QL15_HUMAN] | 2.29 | 5.47 | 1.00 | 128.00 | 14.99 | 11.97 |
| 208 | O95678 | 1.35 | 1.24 | 1.65 | 1.37 | 1.56 | 1.18 | 1.41 | 1.37 | 1.03 | 0.82 | 0.04 | 0.08 | KRT75 | Keratin, type II cytoskeletal 75 OS=Homo sapiens GN=KRT75 PE=1 SV=2 - [K2C75_HUMAN] | 7.52 | 5.44 | 1.00 | 551.00 | 59.52 | 7.74 |
| 209 | B4DJE3 | 1.45 | 1.49 | 1.67 | 1.45 | 1.67 | 1.43 | 1.54 | 1.51 | 1.02 | 0.83 | 0.02 | 0.08 | DDOST | cDNA FLJ52929, highly similar to Dolichyl-diphosphooligosaccharide--proteinglycosyltransferase 48 kDa subunit (EC2.4.1.119) OS=Homo sapiens PE=2 SV=1 - [B4DJE3_HUMAN] | 5.14 | 2.15 | 1.00 | 419.00 | 46.45 | 7.06 |
| 210 | Q8N1C0 | 1.55 | 1.49 | 1.75 | 1.53 | 1.85 | 1.50 | 1.60 | 1.63 | 0.98 | 0.83 | -0.03 | 0.08 | CTNNA1 | CTNNA1 protein OS=Homo sapiens GN=CTNNA1 PE=2 SV=1 - [Q8N1C0_HUMAN] | 8.82 | 9.33 | 3.00 | 536.00 | 59.51 | 5.60 |
| 211 | P35908 | 1.54 | 1.40 | 1.55 | 1.60 | 1.77 | 1.24 | 1.50 | 1.53 | 0.98 | 0.83 | -0.04 | 0.08 | KRT2 | Keratin, type II cytoskeletal 2 epidermal OS=Homo sapiens GN=KRT2 PE=1 SV=2 - [K22E_HUMAN] | 16.21 | 9.08 | 2.00 | 639.00 | 65.39 | 8.00 |
| 212 | B4E0N9 | 1.46 | 1.57 | 1.73 | 1.53 | 1.69 | 1.47 | 1.59 | 1.56 | 1.01 | 0.84 | 0.02 | 0.08 | | Glutamate dehydrogenase OS=Homo sapiens PE=2 SV=1 - [B4E0N9_HUMAN] | 14.21 | 10.82 | 5.00 | 490.00 | 54.22 | 7.58 |
| 213 | B4E1T1 | 1.24 | 1.22 | 1.47 | 1.19 | 1.60 | 1.24 | 1.31 | 1.34 | 0.98 | 0.84 | -0.04 | 0.07 | | cDNA FLJ54081, highly similar to Keratin, type II cytoskeletal 5 OS=Homo sapiens PE=2 SV=1 - [B4E1T1_HUMAN] | 11.87 | 8.11 | 2.00 | 555.00 | 58.81 | 5.97 |
| 214 | B3KUF6 | 1.17 | 1.15 | 1.19 | 1.09 | 1.19 | 1.26 | 1.17 | 1.18 | 0.99 | 0.84 | -0.01 | 0.07 | | cDNA FLJ39748 fis, clone SMINT2017436, highly similar to Homo sapiens haloacid dehalogenase-like hydrolase domain containing 3 (HDHD3), mRNA OS=Homo sapiens PE=2 SV=1 - [B3KUF6_HUMAN] | 4.75 | 7.97 | 1.00 | 251.00 | 27.95 | 6.71 |
| 215 | B4E2K4 | 2.17 | 1.81 | 1.95 | 2.10 | 1.94 | 1.80 | 1.97 | 1.95 | 1.01 | 0.85 | 0.02 | 0.07 | ASPH | cDNA FLJ54576, highly similar to Aspartyl/asparaginyl beta-hydroxylase (EC 1.14.11.16) OS=Homo sapiens PE=2 SV=1 - [B4E2K4_HUMAN] | 2.35 | 3.16 | 3.00 | 729.00 | 83.23 | 4.94 |
| 216 | P30050 | 1.36 | 1.44 | 1.45 | 1.31 | 1.58 | 1.31 | 1.42 | 1.40 | 1.01 | 0.85 | 0.02 | 0.07 | RPL12 | 60S ribosomal protein L12 OS=Homo sapiens GN=RPL12 PE=1 SV=1 - [RL12_HUMAN] | 4.80 | 9.09 | 1.00 | 165.00 | 17.81 | 9.42 |
| 217 | Q15365 | 1.35 | 1.38 | 1.54 | 1.38 | 1.67 | 1.29 | 1.42 | 1.44 | 0.98 | 0.86 | -0.02 | 0.06 | PCBP1 | Poly(rC)-binding protein 1 OS=Homo sapiens GN=PCBP1 PE=1 SV=2 - [PCBP1_HUMAN] | 5.01 | 9.27 | 2.00 | 356.00 | 37.47 | 7.09 |
| 218 | Q04695 | 1.28 | 1.38 | 1.67 | 1.31 | 1.59 | 1.35 | 1.44 | 1.41 | 1.02 | 0.86 | 0.03 | 0.06 | KRT17 | Keratin, type I cytoskeletal 17 OS=Homo sapiens GN=KRT17 PE=1 SV=2 - [K1C17_HUMAN] | 22.10 | 18.98 | 1.00 | 432.00 | 48.08 | 5.02 |
| 219 | P47914 | 1.49 | 1.46 | 1.72 | 1.37 | 1.81 | 1.41 | 1.55 | 1.53 | 1.02 | 0.87 | 0.03 | 0.06 | RPL29 | 60S ribosomal protein L29 OS=Homo sapiens GN=RPL29 PE=1 SV=2 - [RL29_HUMAN] | 3.22 | 14.47 | 2.00 | 159.00 | 17.74 | 11.66 |
| 220 | Q59FZ0 | 1.27 | 1.21 | 1.44 | 1.31 | 1.36 | 1.29 | 1.31 | 1.32 | 0.99 | 0.87 | -0.01 | 0.06 | | LAMP1 protein variant (Fragment) OS=Homo sapiens PE=2 SV=1 - [Q59FZ0_HUMAN] | 4.97 | 2.30 | 1.00 | 392.00 | 41.80 | 7.87 |
| 221 | Q2VPJ6 | 1.20 | 0.98 | 1.40 | 1.16 | 1.39 | 0.93 | 1.19 | 1.16 | 1.03 | 0.88 | 0.04 | 0.06 | HSP90AA1 | HSP90AA1 protein (Fragment) OS=Homo sapiens GN=HSP90AA1 PE=2 SV=1 - [Q2VPJ6_HUMAN] | 35.11 | 8.03 | 2.00 | 585.00 | 68.33 | 5.19 |
| 222 | Q5CAQ5 | 1.63 | 1.61 | 1.84 | 1.62 | 1.88 | 1.53 | 1.69 | 1.67 | 1.01 | 0.88 | 0.02 | 0.05 | TRA1 | Tumor rejection antigen (Gp96) 1 OS=Homo sapiens GN=TRA1 PE=2 SV=1 - [Q5CAQ5_HUMAN] | 42.15 | 11.47 | 10.00 | 802.00 | 92.28 | 4.86 |
| 223 | P40939 | 1.58 | 1.31 | 1.91 | 1.62 | 1.78 | 1.49 | 1.60 | 1.63 | 0.98 | 0.88 | -0.03 | 0.05 | HADHA | Trifunctional enzyme subunit alpha, mitochondrial OS=Homo sapiens GN=HADHA PE=1 SV=2 - [ECHA_HUMAN] | 12.56 | 9.96 | 5.00 | 763.00 | 82.95 | 9.04 |
| 224 | P63104 | 1.87 | 1.57 | 1.92 | 1.66 | 1.97 | 1.66 | 1.79 | 1.76 | 1.01 | 0.88 | 0.02 | 0.05 | YWHAZ | 14-3-3 protein zeta/delta OS=Homo sapiens GN=YWHAZ PE=1 SV=1 - [1433Z_HUMAN] | 37.65 | 33.06 | 3.00 | 245.00 | 27.73 | 4.79 |
| 225 | M0QYG1 | 1.67 | 1.88 | 1.94 | 1.81 | 2.01 | 1.73 | 1.83 | 1.85 | 0.99 | 0.88 | -0.01 | 0.05 | KHSRP | Far upstream element-binding protein 2 (Fragment) OS=Homo sapiens GN=KHSRP PE=4 SV=1 - [M0QYG1_HUMAN] | 2.52 | 15.79 | 1.00 | 114.00 | 12.45 | 7.84 |
| 226 | C9J4S4 | 1.50 | 1.40 | 1.66 | 1.32 | 1.70 | 1.49 | 1.52 | 1.50 | 1.01 | 0.89 | 0.02 | 0.05 | RAB7A | Ras-related protein Rab-7a OS=Homo sapiens GN=RAB7A PE=2 SV=1 - [C9J4S4_HUMAN] | 3.92 | 14.29 | 1.00 | 98.00 | 10.95 | 8.78 |
| 227 | B4E2M8 | 1.50 | 1.24 | 1.41 | 1.34 | 1.56 | 1.30 | 1.38 | 1.40 | 0.99 | 0.89 | -0.02 | 0.05 | | cDNA FLJ61076, highly similar to UBX domain-containing protein 8 OS=Homo sapiens PE=2 SV=1 - [B4E2M8_HUMAN] | 2.49 | 5.66 | 1.00 | 212.00 | 24.73 | 5.74 |

| | | | | | | | | | | | | | | | | | | | | | |
|---|---|---|---|---|---|---|---|---|---|---|---|---|---|---|---|---|---|---|---|---|---|
| 228 | I3L2P8 | 1.53 | 1.47 | 1.74 | 1.51 | 1.77 | 1.39 | 1.58 | 1.56 | 1.01 | 0.90 | 0.02 | 0.04 | | Protein disulfide-isomerase OS=Homo sapiens GN=P4HB PE=2 SV=1 - [I3L2P8_HUMAN] | 60.31 | 17.78 | 8.00 | 450.00 | 50.99 | 4.91 |
| 229 | B4E1K7 | 0.98 | 0.81 | 0.86 | 0.96 | 0.91 | 0.81 | 0.88 | 0.89 | 0.99 | 0.90 | -0.01 | 0.04 | STOML2 | Stomatin-like protein 2 OS=Homo sapiens GN=STOML2 PE=2 SV=1 - [B4E1K7_HUMAN] | 1.64 | 9.97 | 2.00 | 311.00 | 33.32 | 8.25 |
| 230 | Q9NS69 | 1.49 | 1.43 | 1.73 | 1.55 | 1.73 | 1.42 | 1.55 | 1.56 | 0.99 | 0.91 | -0.01 | 0.04 | TOMM22 | Mitochondrial import receptor subunit TOM22 homolog OS=Homo sapiens GN=TOMM22 PE=1 SV=3 - [TOM22_HUMAN] | 4.19 | 7.75 | 1.00 | 142.00 | 15.51 | 4.34 |
| 231 | E9PKV2 | 1.58 | 1.30 | 1.58 | 1.64 | 1.60 | 1.27 | 1.49 | 1.50 | 0.99 | 0.91 | -0.02 | 0.04 | MRPL17 | 39S ribosomal protein L17, mitochondrial (Fragment) OS=Homo sapiens GN=MRPL17 PE=2 SV=1 - [E9PKV2_HUMAN] | 2.53 | 5.63 | 1.00 | 142.00 | 16.36 | 10.48 |
| 232 | P16401 | 1.75 | 1.75 | 2.05 | 1.80 | 2.09 | 1.60 | 1.85 | 1.83 | 1.01 | 0.92 | 0.02 | 0.04 | H1-5 | Histone H1.5 OS=Homo sapiens GN=HIST1H1B PE=1 SV=3 - [H15_HUMAN] | 16.92 | 12.39 | 1.00 | 226.00 | 22.57 | 10.92 |
| 233 | E9PLD0 | 1.70 | 1.77 | 1.91 | 1.76 | 2.00 | 1.58 | 1.79 | 1.78 | 1.01 | 0.92 | 0.01 | 0.04 | RAB1B | Ras-related protein Rab-1B OS=Homo sapiens GN=RAB1B PE=2 SV=1 - [E9PLD0_HUMAN] | 3.91 | 10.06 | 1.00 | 169.00 | 18.47 | 5.72 |
| 234 | Q14BN3 | 1.32 | 1.58 | 1.23 | 1.29 | 1.55 | 1.33 | 1.38 | 1.39 | 0.99 | 0.93 | -0.01 | 0.03 | PKP1 | PKP1 protein OS=Homo sapiens GN=PKP1 PE=2 SV=1 - [Q14BN3_HUMAN] | 3.00 | 4.19 | 1.00 | 334.00 | 36.79 | 8.56 |
| 235 | B4E241 | 1.54 | 1.35 | 1.47 | 1.37 | 1.66 | 1.35 | 1.45 | 1.46 | 0.99 | 0.95 | -0.01 | 0.02 | SRSF3 | Serine/arginine-rich-splicing factor 3 OS=Homo sapiens GN=SFRS3 PE=2 SV=1 - [B4E241_HUMAN] | 16.43 | 15.32 | 2.00 | 124.00 | 14.19 | 10.08 |
| 236 | O60250 | 1.76 | 1.76 | 1.84 | 1.79 | 2.16 | 1.36 | 1.79 | 1.77 | 1.01 | 0.96 | 0.01 | 0.02 | | Ribosomal protein L13 (Fragment) OS=Homo sapiens PE=2 SV=2 - [O60250_HUMAN] | 3.88 | 20.00 | 1.00 | 65.00 | 7.39 | 10.27 |
| 237 | Q9Y2R5 | 1.56 | 1.40 | 1.25 | 1.33 | 1.68 | 1.23 | 1.40 | 1.41 | 0.99 | 0.96 | -0.01 | 0.02 | MRPS17 | 28S ribosomal protein S17, mitochondrial OS=Homo sapiens GN=MRPS17 PE=2 SV=1 - [RT17_HUMAN] | 3.33 | 8.46 | 1.00 | 130.00 | 14.49 | 9.85 |
| 238 | Q2Z195 | 1.39 | 1.34 | 1.45 | 1.31 | 1.60 | 1.29 | 1.39 | 1.40 | 1.00 | 0.96 | -0.01 | 0.02 | HLA-A | MHC class I antigen (Fragment) OS=Homo sapiens GN=HLA-A PE=2 SV=1 - [Q2Z195_HUMAN] | 2.23 | 12.36 | 1.00 | 89.00 | 10.27 | 6.05 |
| 239 | I3L3P7 | 1.75 | 1.87 | 2.00 | 1.86 | 2.07 | 1.67 | 1.87 | 1.87 | 1.00 | 0.96 | 0.01 | 0.02 | RPS15A | 40S ribosomal protein S15a OS=Homo sapiens GN=RPS15A PE=2 SV=1 - [I3L3P7_HUMAN] | 1.68 | 8.00 | 1.00 | 100.00 | 11.47 | 10.15 |
| 240 | Q9Y5B2 | 1.59 | 1.54 | 1.62 | 1.55 | 1.74 | 1.46 | 1.59 | 1.58 | 1.00 | 0.96 | 0.00 | 0.02 | | Junction adhesion molecule OS=Homo sapiens PE=2 SV=1 - [Q9Y5B2_HUMAN] | 10.01 | 13.13 | 2.00 | 259.00 | 28.10 | 8.29 |
| 241 | Q7Z434 | 0.99 | 1.12 | 1.38 | 1.10 | 1.45 | 0.96 | 1.16 | 1.17 | 0.99 | 0.97 | -0.01 | 0.01 | MAVS | Mitochondrial antiviral-signaling protein OS=Homo sapiens GN=MAVS PE=1 SV=2 - [MAVS_HUMAN] | 4.71 | 5.19 | 1.00 | 540.00 | 56.49 | 5.52 |
| 242 | P06733 | 1.75 | 1.43 | 2.11 | 1.69 | 1.82 | 1.76 | 1.76 | 1.76 | 1.00 | 0.97 | 0.01 | 0.01 | ENO1 | Alpha-enolase OS=Homo sapiens GN=ENO1 PE=1 SV=2 - [ENOA_HUMAN] | 15.10 | 7.83 | 3.00 | 434.00 | 47.14 | 7.39 |
| 243 | E7EX53 | 2.21 | 1.88 | 2.09 | 1.91 | 2.29 | 1.96 | 2.06 | 2.05 | 1.00 | 0.97 | 0.00 | 0.01 | RPL15 | Ribosomal protein L15 (Fragment) OS=Homo sapiens GN=RPL15 PE=2 SV=1 - [E7EX53_HUMAN] | 2.88 | 6.77 | 1.00 | 133.00 | 15.71 | 11.00 |
| 244 | J3QRP6 | 1.53 | 1.22 | 1.55 | 1.29 | 2.01 | 1.02 | 1.43 | 1.44 | 0.99 | 0.98 | -0.01 | 0.01 | SLC9A3R1 | Na(+)/H(+) exchange regulatory cofactor NHE-RF1 (Fragment) OS=Homo sapiens GN=SLC9A3R1 PE=4 SV=1 - [J3QRP6_HUMAN] | 2.86 | 12.09 | 1.00 | 215.00 | 22.86 | 4.82 |
| 245 | Q02539 | 1.92 | 1.79 | 2.32 | 1.87 | 2.31 | 1.85 | 2.01 | 2.01 | 1.00 | 1.00 | 0.00 | 0.00 | H1-1 | Histone H1.1 OS=Homo sapiens GN=HIST1H1A PE=1 SV=3 - [H11_HUMAN] | 16.36 | 12.09 | 1.00 | 215.00 | 21.83 | 10.99 |
| 246 | F5GWP8 | 1.48 | 1.60 | 2.05 | 1.63 | 1.91 | 1.60 | 1.71 | 1.71 | 1.00 | 1.00 | 0.00 | 0.00 | KRT17 | Junction plakoglobin OS=Homo sapiens GN=JUP PE=2 SV=1 - [F5GWP8_HUMAN] | 17.82 | 12.69 | 1.00 | 591.00 | 66.31 | 5.19 |
| 247 | P62269 | 1.74 | 1.80 | 1.92 | 2.01 | 1.98 | 1.48 | 1.82 | 1.82 | 1.00 | 1.00 | 0.00 | 0.00 | RPS18 | 40S ribosomal protein S18 OS=Homo sapiens GN=RPS18 PE=1 SV=3 - [RS18_HUMAN] | 1.91 | 9.87 | 2.00 | 152.00 | 17.71 | 10.99 |
