## Supplementary Figure 1 for "CMTM6 drives cisplatin resistance in OSCC by regulating AKT mediated Wnt signaling"

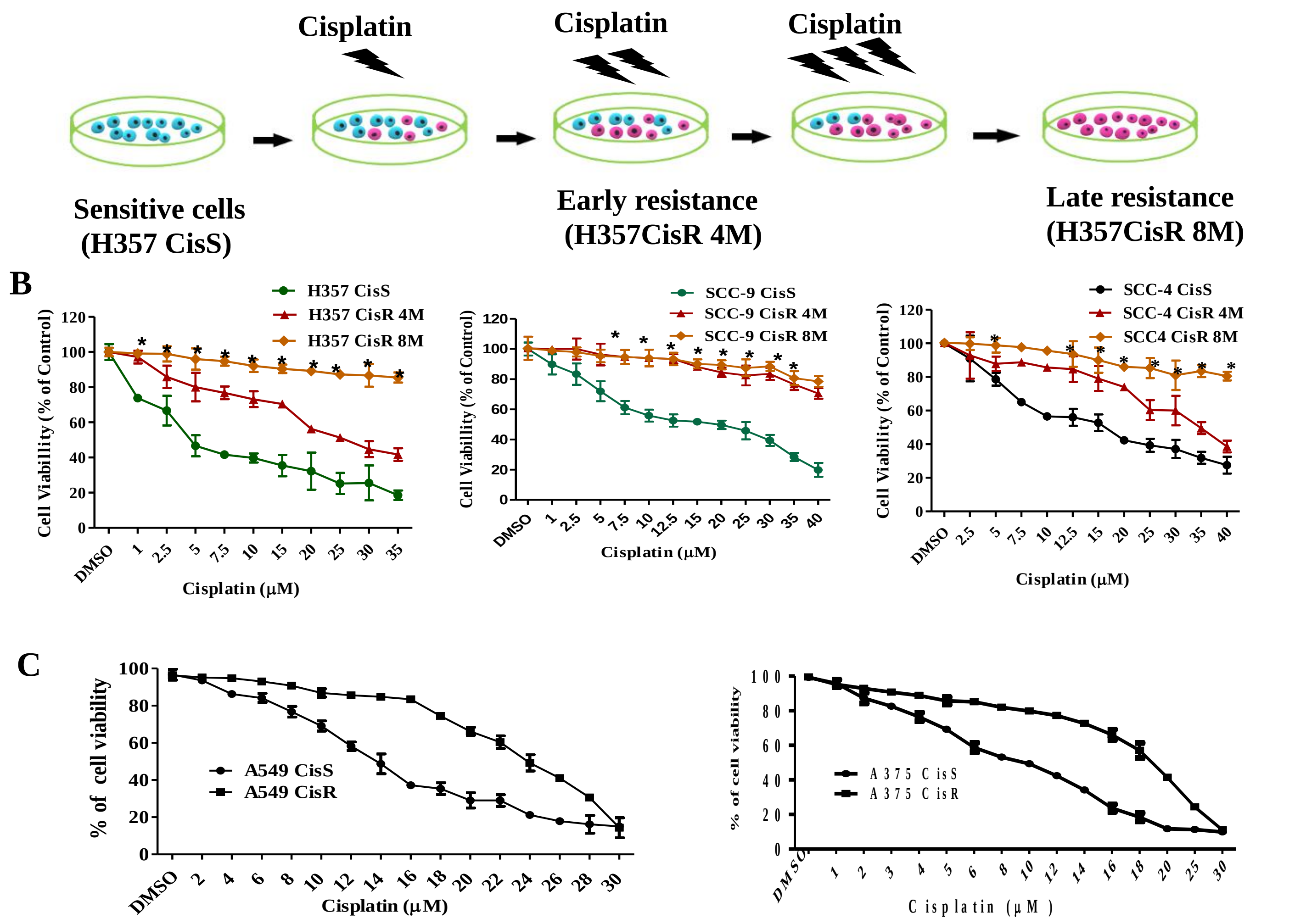


**Supplementary figure 1: Characterization of sensitive, early and late cisplatin resistant lines: A)** Schematic presentation of establishing sensitive, early and late cisplatin resistant cancer lines **B)** Sensitive, early and late cisplatin resistant pattern (CisS, CisR4M and CisR8M) of H357, SCC-9 and SCC-4 cells were treated with indicated concentration of cisplatin for 48h and cell viability was determined by MTT assay (n=3, *: $P < 0.05$). **C)** Sensitive and late cisplatin resistant lung cancer (A549) and melanoma (A375) lines were established as described in method section. Sensitive and resistant cells were treated with cisplatin with indicated concentration for 48h and cell viability was determined by MTT assay (n=3).
