## Supplementary Figure 2 for "CMTM6 drives cisplatin resistance in OSCC by regulating AKT mediated Wnt signaling"

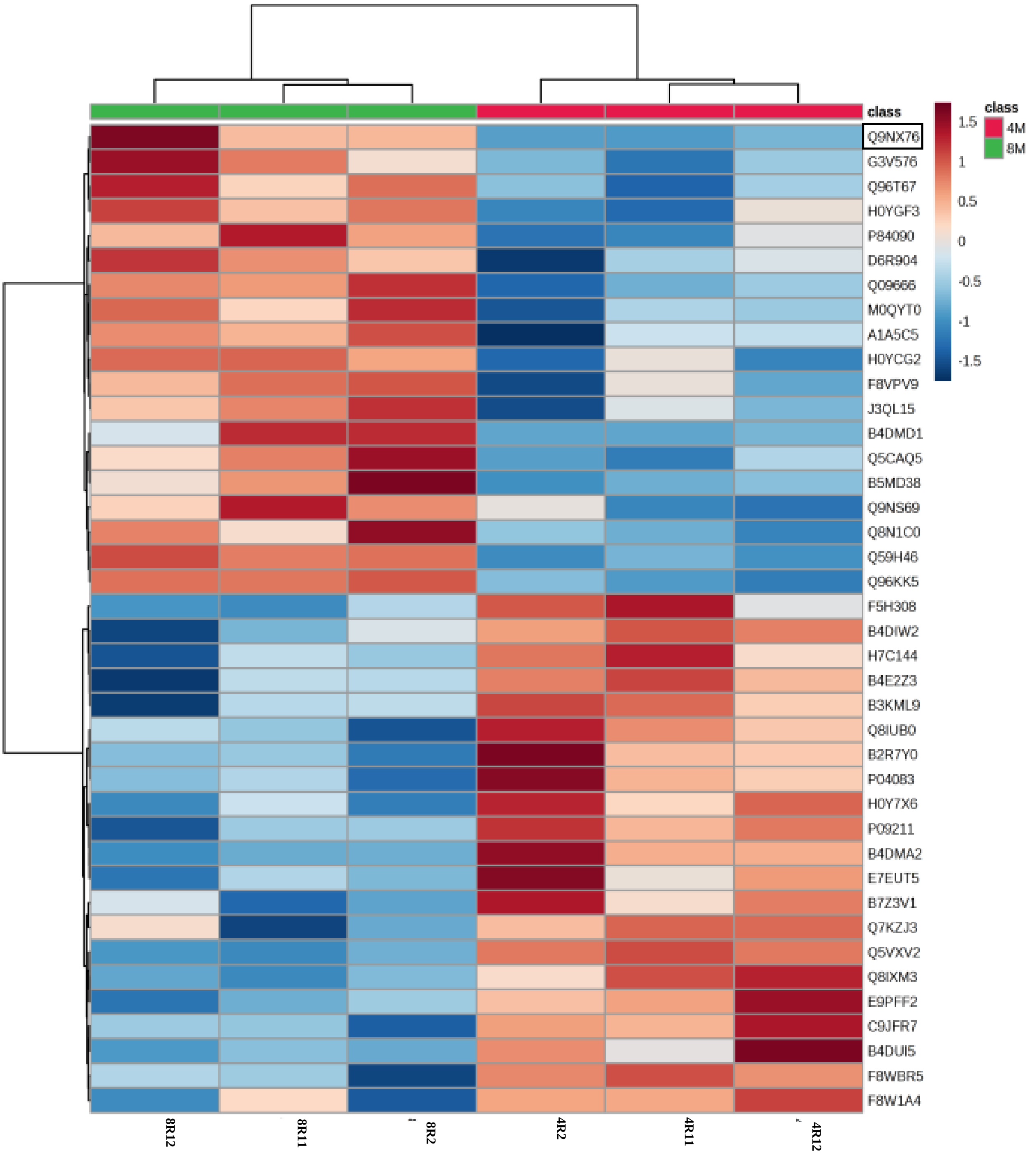


**Supplementary figure 2: Global proteomic profiling of sensitive, early and late chemoresistant cells.** The lysates were isolated from parental sensitive (H357CisS), early (H357CisR4M) and late (H357CisR8M) cisplatin resistant cells and subjected to global proteomic profiling. The dendrogram represents the deregulated genes from proteomic analysis.
