## Supplementary Figure 3 for "CMTM6 drives cisplatin resistance in OSCC by regulating AKT mediated Wnt signaling"

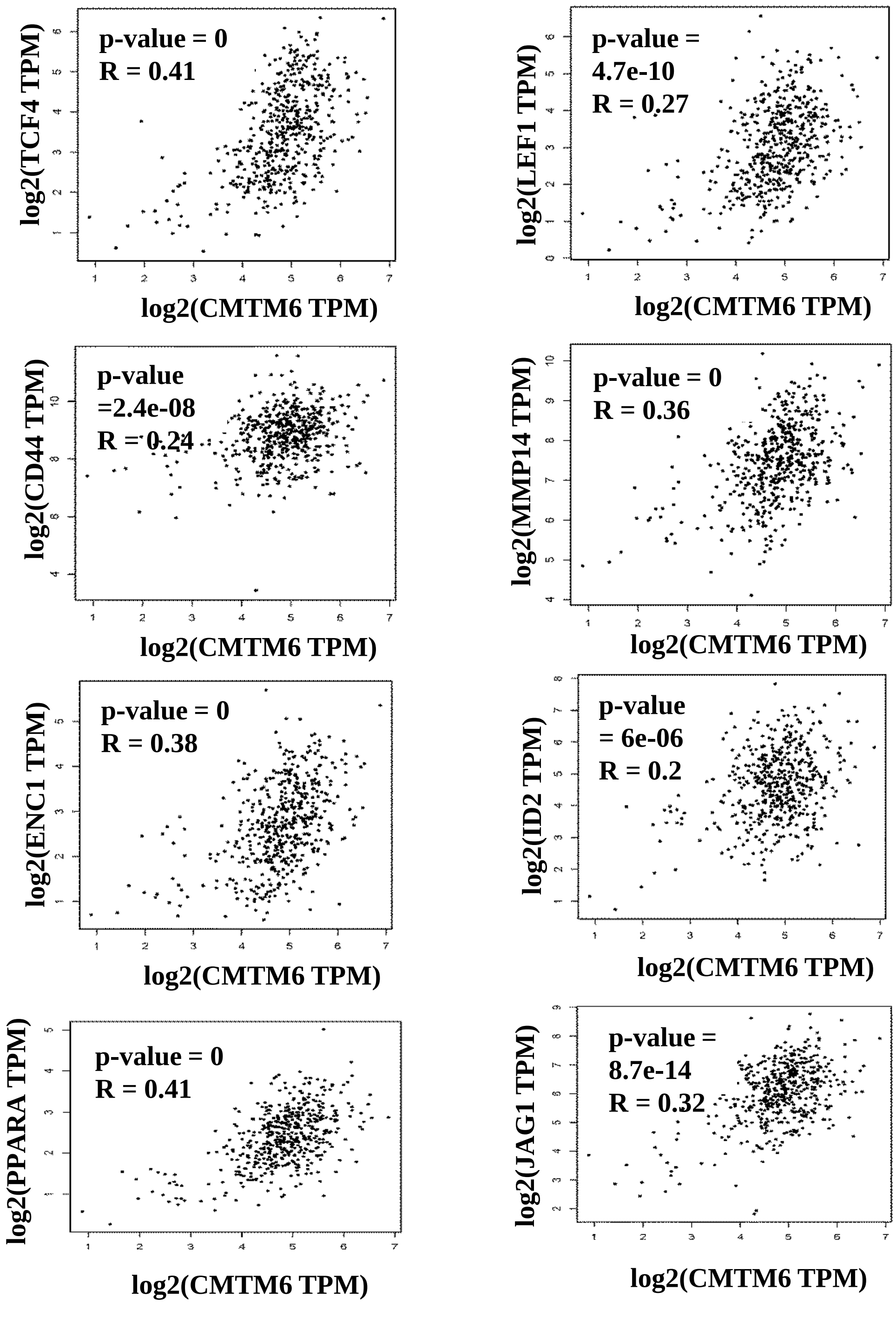


**Supplementary figure 3: CMTM6 correlates with Wnt signalling target genes in tumor of HNSCC patients:** Expression correlation between CMTM6 and Wnt target prosurvival genes (TCF4, LEF1, CD44, MMP14, ENC1, ID2, PPARA and JAG1) RNA expression in the TCGA HNSCC database. Correlation was analyzed using Spearman's correlation coefficient test, n = 520. The analysis was performed in Gene expression profiling interactive analysis (GEPIA) platform.
