## Supplementary Figure 4 for "CMTM6 drives cisplatin resistance in OSCC by regulating AKT mediated Wnt signaling"

A

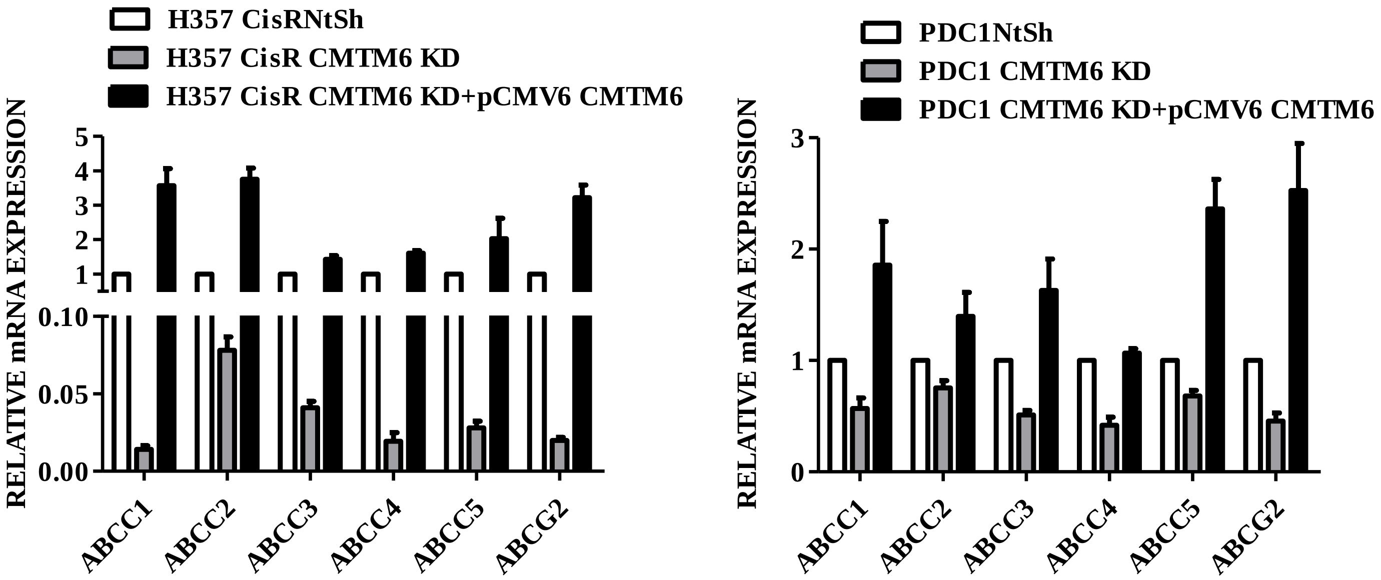


B

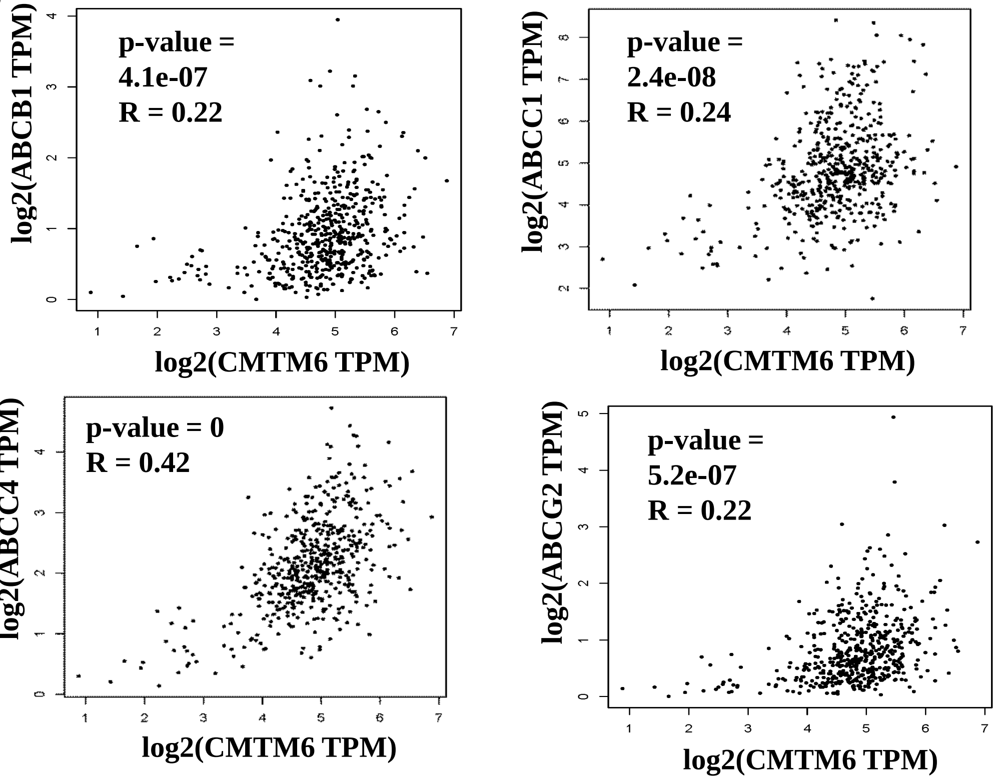


**Supplementary Fig 4: CMTM6 regulates the expression of ABC transporters**: A) CMTM6 was transiently overexpressed in chemoresistant cells stably expressing CMTM6ShRNA#2 and qRTPCR was performed for indicated genes to evaluate relative mRNA expression (mean ±SEM, n=3). **B)** Expression correlation between CMTM6 and ABC transporter genes (ABCB1,ABCC1, ABCC4 and ABCG2) mRNA in the TCGA HNSCC database. Correlation was analyzed using Spearman's correlation coefficient test, n = 520. The analysis was performed in Gene expression profiling interactive analysis (GEPIA) platform.
